# DecoDiPhy: Deconvolving Phylogenetic Distance Mixtures

**DOI:** 10.64898/2026.01.18.700179

**Authors:** Shayesteh Arasti, Ali Osman Berk Şapcı, Eleonora Rachtman, Mohammed El-Kebir, Siavash Mirarab

**Affiliations:** Department of Computer Science and Engineering, UC San Diego, CA 92093, USA; Bioinformatics and Systems Biology Graduate Program, UC San Diego, CA 92093, USA; Siebel School of Computing and Data Science, University of Illinois Urbana-Champaign, Urbana, IL 61801, USA; Department of Electrical and Computer Engineering, UC San Diego, CA 92093, USA

**Keywords:** Mixture deconvolution, Phylogenetic placement, Metagenomics, Phylogenetic mixture analysis, Differential Abundance

## Abstract

Mixtures of multiple organisms are sequenced in many applications, including metagenomics, but characterizing their composition faces two challenges: constituents are evolutionarily related, and individual short reads provide limited signal. Here, we introduce phylogenetic distance deconvolution (PDD), a framework that jointly addresses these challenges by using distances from mixtures to reference genomes to place samples on multiple branches of a phylogeny with associated abundances. We develop both exact and heuristic algorithms, implemented in DecoDiPhy. In metagenomic analyses, DecoDiPhy consolidates overdispersed per-read placements into a small set of phylogenetic features, thereby improving interpretability and enhancing downstream phenotype differentiation and differential abundance analysis.

## Background

In many modern applications, DNA is sequenced not from a single organism but from a mixture. Examples abound and include sequencing of microbiome communities [1], heterogeneous cancer tumors [2], and true biological mixtures, such as admixtures [3] and recent hybrid species [4]. A first step in analyzing these mixtures is to characterize their contents relative to a reference dataset of known organisms, which presents several challenges. The constituents of a “query” mixture *and* the references are the product of an evolutionary process. Thus, query organisms are at some evolutionary distance from references (“novelty” hereafter), and while denser reference sets reduce novelty, in many applications, high levels of novelty remain, making simple sequence matching incapable of fully characterizing a sample [5–7]. Relatedly, the shared evolutionary history creates dependencies, and as a result, mixture identification should go beyond classifying into independent buckets. What makes the identification even more challenging is that mixture data most often consists of short reads with limited signal per read. The rich literature on mixture identification [8] and particularly microbiome analysis [9, 10], which is our focus, has sought to address these challenges by designing methods to capture dependencies among species or among reads (Table S1).

Despite the rich literature, existing methods typically model either evolutionary dependencies or read dependencies, but not both. Most methods crudely model evolutionary histories by grouping reference genomes into predefined hierarchical taxonomic groups or data-derived [11–13] operational taxonomic/genomic units (OTUs/OGUs), and treat groups within a higher-rank category as independent. This provides limited modeling of dependencies, and the resulting taxonomic profiles have less resolution than those using phylogenetic trees [14, 15]. Methods for phylogenetic characterization also exist [16, 17], especially phylogenetic placement [18–22]. A rich toolkit uses phylogenetic profiles in downstream analyses such as comparing and clustering samples [23, 24], extracting phylogeny-aware features [25] for supervised learning [26–28], and differential abundance across phenotypes [29]. Separately, whether they use taxonomy [30– 33] or phylogeny [18–22], most methods characterize individual reads and summarize results, often using marker genes [34–36]. The summarization step can be as simple as averaging [35] or can further attempt to denoise the profile using prior expectations [37]. An alternative is to characterize the entire sample without separating reads [38]. For example, *k*-mer sketching methods find which reference genomes seem present in a sample as a whole [39–41]. However, these methods often ignore evolutionary dependency, with limited modeling in some cases [42]. Thus, while the field has recognized the roles of both evolutionary *and* across-read dependencies, existing methods are limited in their ability to jointly address both challenges.

We address both challenges simultaneously by *i*) using a reference phylogeny instead of a taxonomy, while *ii*) placing the entire sample instead of placing each read on the phylogeny. We take advantage of the metric space of trees defined as the pairwise path length between leaves. has focused on input matrices in which each row or column corresponds to a single leaf. The goal of this work is to go beyond this setting and enable distance-based phylogenetic placement using input distances computed from mixtures. Using these calculated mixture distances, obtained with external tools, we seek a *small* number of placements that best explain mixture distances; the output, a multi-placement on the tree, deconvolves the input distance vector into its constituents and their relative abundances. This approach, to our knowledge, has not been pursued directly, though deconvolution without trees [38] and placement on exactly two edges [43] have been. We test the resulting method, DecoDiPhy, on metagenomic data and show that by consolidating over-dispersed placements, it improves the accuracy of profiles and elucidates features associated with environments and clinical outcomes.

## Results

### Phylogenetic Distance Deconvolution (PDD) and DecoDiPhy

We first pose phylogenetic mixture analyses as the Phylogenetic Distance Deconvolution (PDD) problem (see Methods for formal definition). We are given a reference tree *R* and a mixture query (Fig. 1a). The standard placement, adding the query as a single leaf [18], is insufficient because the input is a mixture. Instead, PDD seeks to insert the query into *k* edges of the tree. For any set of *k query* leaves *Q* on a tree and a relative abundance vector **p**, we define their total mixture distance to a non-query *reference* leaf *r* as:

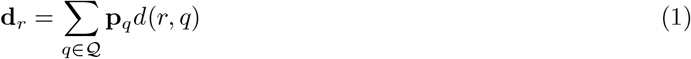

where *d*(*r, q*) is the path length between between reference leaf *r* and query leaf *q* on the tree (Fig. 1b). On real data, we do not have access to true mixture distances **d**, but we can *estimate* it from sequence mixtures to obtain 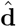. PDD can work with any estimator, but we explore two here (see Methods): a sequence-based approach based on the Jukes-Cantor (JC69) [44] model and a 2-step approach that first places each read on one or multiple edges (to account for uncertainty) and then uses initial placements to compute mixture distances using Eq. (1) (Fig. 1b). In the second mode, PDD consolidates noisy placements into *k* positions.

**Figure 1:**
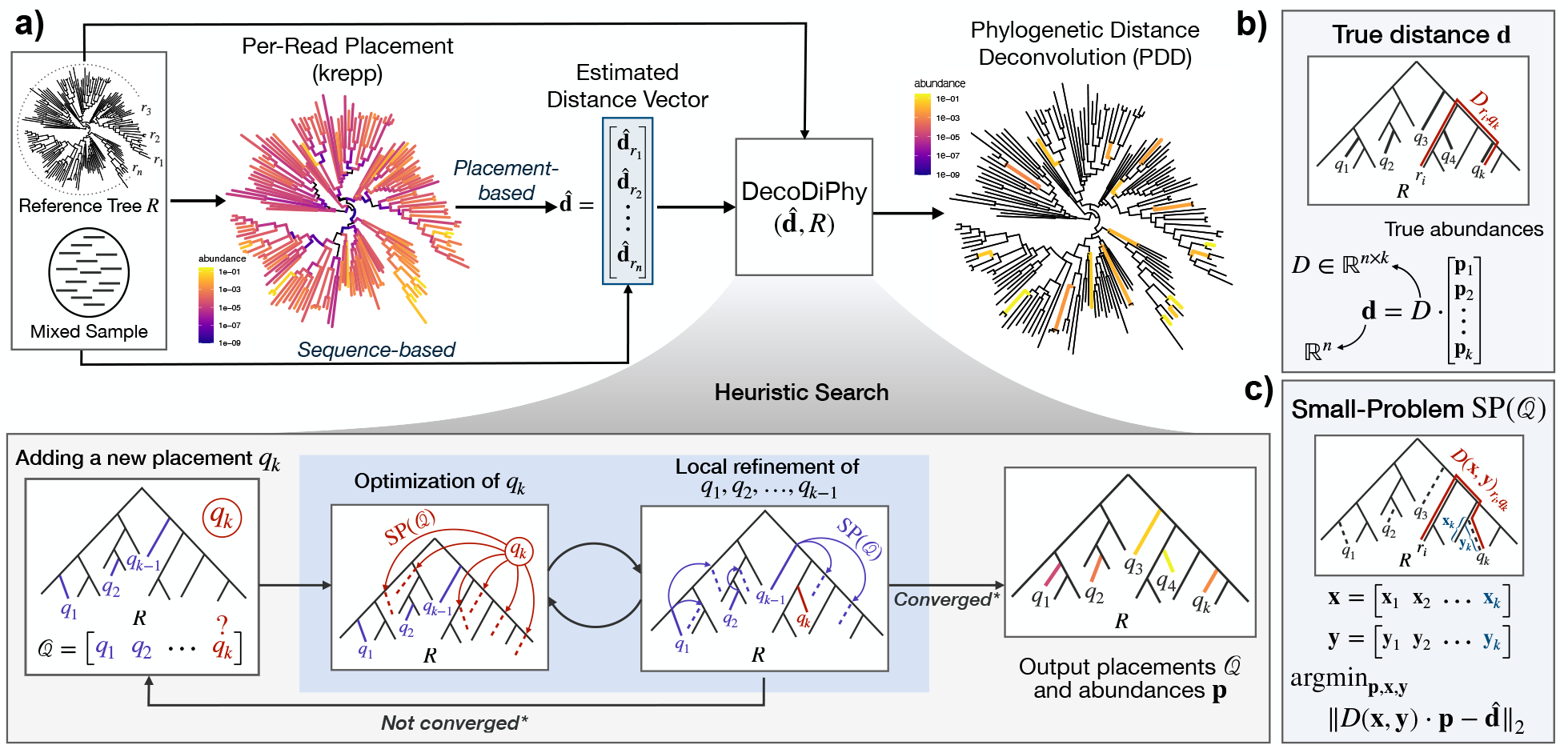
**a)** The phylogenetic Distance Deconvolution (PDD) problem solved by DecoDiPhy. Given a reference tree and a mixed sequence sample, mixture distances to references, 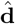, are calculated either directly from sequences or by placing all reads using an external tool. PDD deconvolutes the mixture distances by placing the sample on *k* places. DecoDiPhy solves PDDs using a heuristic search with greedy additions followed by local refinements (Alg. S1). **b)** PDD assumes that the true distance vector **d** follows Eq. (1); the distance from the mixed sample to each query is a linear combination of the individual distances of the query taxa, weighted by their abundances. Here, *n* denotes the number of reference taxa. **c)** The search repeatedly solves the small-problem of optimizing continuous variables (abundance **p**, pendant **y**, and distal **x** lengths with fixed placements) for fixed placements, which we turn into a quadratic programming problem.

The goal of PDD is to recover the placement of *k* unknown queries and their **p** given the reference tree and 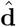. The problem can be posed as a *ℓ*_2_ optimization. The exact placement of any query leaf *q* can be characterized with three variables : the placement edge **e**_*q*_, the distal edge length **x**_*q*_, and the pendant edge length **y**_*q*_ (Fig. S1a); therefore, any set **e** of *k* placement edges, together with pendant and distal edge lengths **y, x**, and relative abundances **p** define a multi-placement, which implies a mixture distance **d**(**e, p, x, y**) through Eq. (1) (Fig. 1c and Fig. S1). Then, the objective is to find

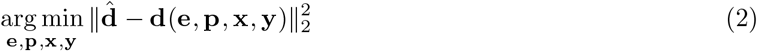

The PDD has a discrete component (placement edges, **e**) and several continuous ones. For any given **e**, optimizing the continuous variables is called the “small-problem”, and through a series of algebraic manipulations, we can turn the small-problem into a quadratic optimization with linear constraints (Problem 2 in Methods). As a result, the small problem can be solved efficiently using existing optimizers such as OSQP [45], which we use. For finding placement edges, the brute-force approach of examining all possible *k* edges (Definition 1, Methods) does not scale. Thus, we developed a heuristic algorithm (Alg. S1) that adds one query at a time, optimizing Eq. (2) for all possible placements and retaining the optimal solution, then performs local refinements of the previous placements (Fig. 1a). Letting *n* denote the number of leaves of *R*, the algorithm solves the small-problem *O*(*nk*) times, and empirically scales quadratically with respect to *n* if 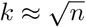 in time and memory (Fig. 2a). We have also developed heuristic stopping criteria to decide what *k* is sufficient for a given sample (see Methods).

**Figure 2:**
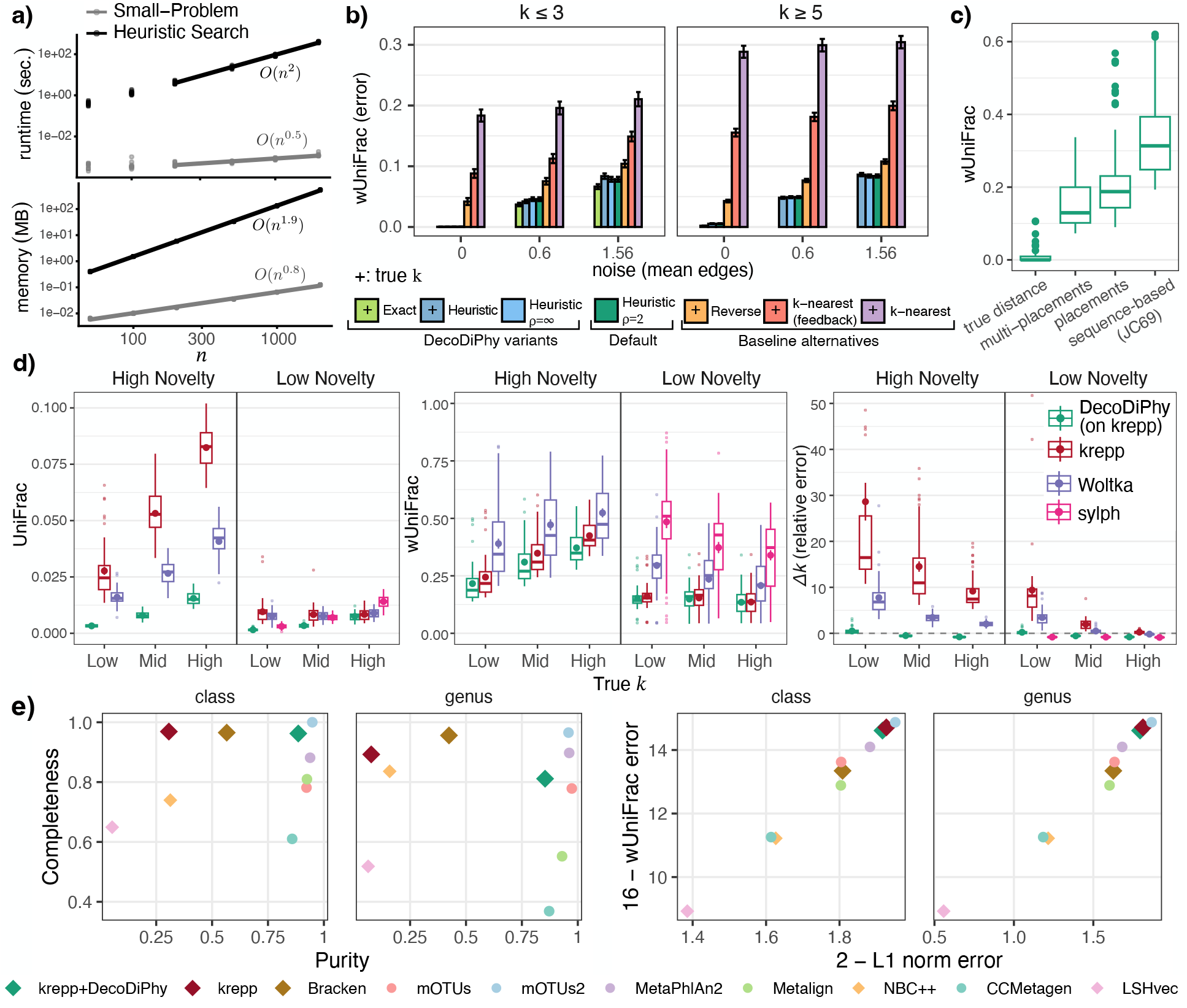
**a)** Empirical running time and memory versus *n* for 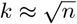, showing log-log plots; line slopes estimate the asymptotic scaling. **b)** Impact of search algorithm on E1 dataset. Bars show the mean and standard error (across 120 data points for noise=0 and 240 data points for other noise levels) for variants of DecoDiPhy and three baseline methods, which use our small-problem. The “Exact’ variant could be run only for *k <* 4. *X*–axis: noise in input distance, measured as the mean number of branches by which each placement is moved. +: the method is given the true *k. ρ*: refinement radius (Methods). See also Fig. S4. **c)** Impact of input mixture distances, comparing true distances, distances from placement (krepp), or from sequences (Jukes-Cantor correction), on two small microbial subtrees (E2). See also Fig. S5. **d)** Metagenome simulations (E3). For six model conditions (two novelty levels × three true *k*s), 50 replicates each, we compare independent read placement (krepp), DecoDiPhy run on krepp multiplacements, single read mapping (Woltka), and joint sample mapping (sylph). On high novelty, sylph found no matches. Δ*k* shows estimated − true *k* normalized by true *k*. Average true *k* for low, med, and high model conditions is 33, 102, and 214, respectively. We limited UniFrac and Δ*k* to placements with abundance at least 10^*−*4^. For each metric, boxplots as well as mean and standard error (dot and error bars) are shown for each method. See Fig. S3. **e)** CAMI-II Dataset (E4). Performance of taxonomic profiling on marine dataset for class and genus taxonomic levels. Panels show purity versus completeness (left), and 2−L1 norm versus 16−weighted UniFrac (right). Note that L1 norm and weighted UniFrac are by design bounded to [0, 2] and [0, 16], respectively. Closer to top right corner is better. *k*-mer-based methods as well as krepp+DecoDiPhy are shown with a diamond shape, while alignment-based methods are shown with circles. See Fig. S10 for all taxonomic levels.

### Simulation studies

#### Effectiveness of DecoDiPhy on simulated mixture distances

We first evaluate the effectiveness of our heuristic search algorithm using a set of 12 simulated datasets in a leave-out experiment (**E1**; Methods). We primarily report accuracy using weighted UniFrac (wUniFrac), which incorporates the reference tree, placements, and abundances, and additionally report UniFrac (which ignores abundances) and the Jaccard index (which further ignores the reference tree); see Methods.

Given true distances, DecoDiPhy recovers the true placements in all cases using the exhaustive search (Definition 1), indicating that cases in which PDDs are not identifiable are rare (Fig. 2b). The heuristic search achieves near-perfect accuracy when provided with the true values of *k* and **d**, demonstrating that the search procedure itself is effective. The heuristic used to select *k* occasionally underestimates the true value (Fig. S4), resulting in an approximately 10% reduction in Jaccard accuracy. Nevertheless, wUniFrac error remains low (Fig. 2b), indicating that missed placements tend to be of low abundance and that inferred placements are phylogenetically close to the true ones. Increasing the aggressiveness of the search (see Methods) does not improve accuracy.

Because no existing method directly addresses PDD inference, we designed three alternative approaches for comparison (Methods). None achieved accuracy comparable to DecoDiPhy. The nearest-*k*-neighbor approach exhibited high error; although a feedback mechanism reduced this error, it did not make the method competitive. The most accurate alternative was a reverse search based on our small-problem formulation, which substantially outperformed nearest neighbors but remained less accurate than DecoDiPhy. When noise was added to the input distances (Methods), Jaccard accuracy decreased for all methods, and other error measures increased (Fig. S4). In contrast, wUniFrac error increased only modestly (Fig. 2b), indicating that errors under noise tend to involve phylogenetically close and low-abundance placements. Under noisy input, the exact and heuristic versions of DecoDiPhy achieved nearly identical wUniFrac accuracy.

We next examined strategies for computing the mixture distance 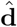using a small simulated microbiome dataset consisting of two trees with 34 and 111 taxa (**E2**; Methods). When distances were inferred from simulated reads, we observed a substantial reduction in Jaccard accuracy compared to true distance (Fig. S5), whereas degradation measured by the tree-aware wUniFrac metric was more modest (Fig. 2c). We observed clear differences among methods for estimating 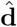, with per-read placements (using krepp) consistently outperforming sequence-based methods, and multi-placements, which account for uncertainty, better than single placements. Placement-based estimates of 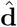 also showed stronger correlation with the true mixture distance **d** than sequence-based estimates; Pearson correlation across replicates: 0.89 for multi-placements and 0.69 for sequence-based with JC69 correction distances (Fig. S6). JC69 distances had an underestimation bias, which F81 correction reduced substantially (Fig. S6); however, JC69 and F81 had similar correlation levels (0.70 for F81), and when used as input to DecoDiPhy, accuracy did not improve (Fig. S5). Generally, the estimation error in 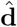 can significantly impact the accuracy of DecoDiPhy (Fig. 2c). The impact of the input error on DecoDiPhy placements is especially clear when the error is judged by wUniFrac, while UniFrac error shows robustness to input error (Fig. S7). Finally, our heuristics for estimating *k* recovered values in the correct range but exhibited a modest tendency toward overestimation under noisy inputs, particularly for smaller true values of *k* (Fig. S5).

Moving forward, we use multi-placements using krepp to obtain 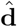, thus, testing DecoDiPhy as a means to consolidate and denoise read placements.

#### Comparison to other methods on large-scale simulated metagenomes

We next simulated metagenomic datasets (**E3**; Methods) using leave-out experiments on the WoL-2 [46] reference tree with 15,953 leaves (see Methods), controlling novelty (distance of queries to closest reference), and complexity (number of queries, *k*), producing a range for both (Fig. S3). We first revisited the question of DecoDiPhy’s search effectiveness and identifiability using true distances (Table S2). Because exhaustive search was computationally infeasible in these settings, we evaluated DecoDiPhy using the search heuristic while providing the true value of *k*. Across 480 replicates, DecoDiPhy failed to recover all true placement branches in only 26 cases (Table S2). Even in these cases, accuracy remained high (mean Jaccard index of 92%), while UniFrac and wUniFrac errors were low (mean 0.000443 and 0.00153, respectively), indicating that inferred placements remained phylogenetically close to the true ones (Fig. S2). Thus, the search algorithm is effective, and identifiability is not a practical issue, leaving us to next examine the impact of distances estimated using krepp’s multi placements.

Compared to its input (krepp), DecoDiPhy+krepp dramatically reduces the placement error for novel and low complexity conditions, with smaller improvements for low novelty but high complexity input (Fig. 2d). Since no other method is available for scalable phylogenetic placement of genome-wide reads, we next compared DecoDiPhy to alternatives designed for other purposes, with their assignments mapped to leaves of the phylogeny. In this setting, DecoDiPhy is far more accurate than read mapping using Woltka, which internally uses bowtie2, and the joint sample method sylph, which is based on *k*-mer containment (Fig. 2d). Improvements are also observed for abundance-free metrics UniFrac and Jaccard (Fig. S8).

The accuracy of all methods depended on novelty. With higher novelty, sylph, which was quite accurate on lower novelty data, was unable to match any queries. Note that sylph is designed to find highly similar genomes contained in a sample, not for the phylogenetic placement of novel species; thus, this behavior is not surprising. DecoDiPhy had the lowest degradation in accuracy as novelty increased. The most dramatic improvements with DecoDiPhy are observed for UniFrac for the high novelty condition, where error is reduced by 9× and 4× compared to krepp and Woltka, respectively (improvements are even larger if we don’t filter low abundances; see Fig. S8). The improvements in wUniFrac are smaller than UniFrac, consistent with the notion that DecoDiPhy consolidates krepp’s placements, reducing noisy low-abundant placements.

The complexity of the sample (true *k*) also mattered, though less than novelty (in our setup, the two are not fully independent; Fig. S3). UniFrac error increases with complexity, but wUniFrac only increases in high novelty conditions. The detected number of queries was too high by 5–30× in most conditions for krepp and Woltka. Both DecoDiPhy and sylph estimated *k* in the correct range (Fig. S8), with sylph detecting slightly fewer queries than present, while DecoDiPhy detected slightly too many for low complexity and too few for high complexity cases.

#### Taxonomic profiling of CAMI-II dataset

While DecoDiPhy is designed for phylogenetic placement, we asked whether it can also produce accurate taxonomic profiles. We studied the 10-sample marine dataset from the CAMI-II challenge [47] with 777 distinct genomes appearing as queries across all ten samples (**E4**; Methods). To translate DecoDiPhy’s phylogenetic placements to taxonomic labels, we simply chose the least common ancestor (LCA) of the taxonomic labels below a placement edge (details in Methods). Across all 10 replicates of this dataset, DecoDiPhy produced 1383 distinct placements (409 terminal and 974 internal), whereas krepp produced 101, 481 distinct placements. Using DecoDiPhy on top of krepp drastically improved the purity (Fig. 2e) of the profiles (an *increase* of 58% for class and 78% for genus), with minimal decreases in completeness (0.7% for class and 8.2% for genus). The only exception was at the species level, where DecoDiPhy was still far purer but also substantially less complete than krepp (43% vs. 84% for krepp; Fig. S10a). Bracken, a *k*-mer-based method specifically designed for taxonomic profiling, had the highest overall completeness compared to krepp and DecoDiPhy, with 95% average completeness compared to 91% and 83% for krepp and DecoDiPhy, respectively (Fig. S10a). However, DecoDiPhy outperformed Bracken in terms of purity, with profiles that were on average 36% more pure across all taxonomic levels (Fig. S10a). In terms of abundance-weighted metrics, weighted UniFrac and L1 norm error, both krepp and DecoDiPhy outperformed Bracken on both metrics across all levels (Fig. 2e and Fig. S10b). Overall, pairing DecoDiPhy with krepp dramatically improves its purity at little cost to completeness and abundance profile accuracy. It outperforms all other *k*-mer-based methods and is outperformed only by one marker-based method, mOTU2.

### Biological metagenomic analyses (IBD and EMP)

#### Sample-level phylogenetic consolidation improves separation with compact feature sets

We evaluated DecoDiPhy on two real metagenomic datasets representing contrasting levels of reference novelty: an inflammatory bowel disease (IBD) cohort of 220 human gut samples [48] and 210 free-living microbial communities from the Earth Microbiome Project (EMP) [49] (See supplementary section C.7.6). Both datasets were analyzed against the 15,953-taxon WoL2 reference tree [46]. Compared to the environmentally diverse EMP samples, gut-associated samples exhibit lower novelty relative to the reference.

Across both datasets, DecoDiPhy consolidated krepp’s read-level placements into two to three orders of magnitude fewer branches, creating fewer phylogenetic features while retaining the majority of mapped signal (Fig. 3a). On the less novel IBD dataset, all methods except for Woltka were able to map a large number of reads, with krepp (and by extension DecoDiPhy) placing slightly more. On EMP, krepp and Kraken mapped orders of magnitude more reads than Woltka, sylph, or sourmash.

**Figure 3:**
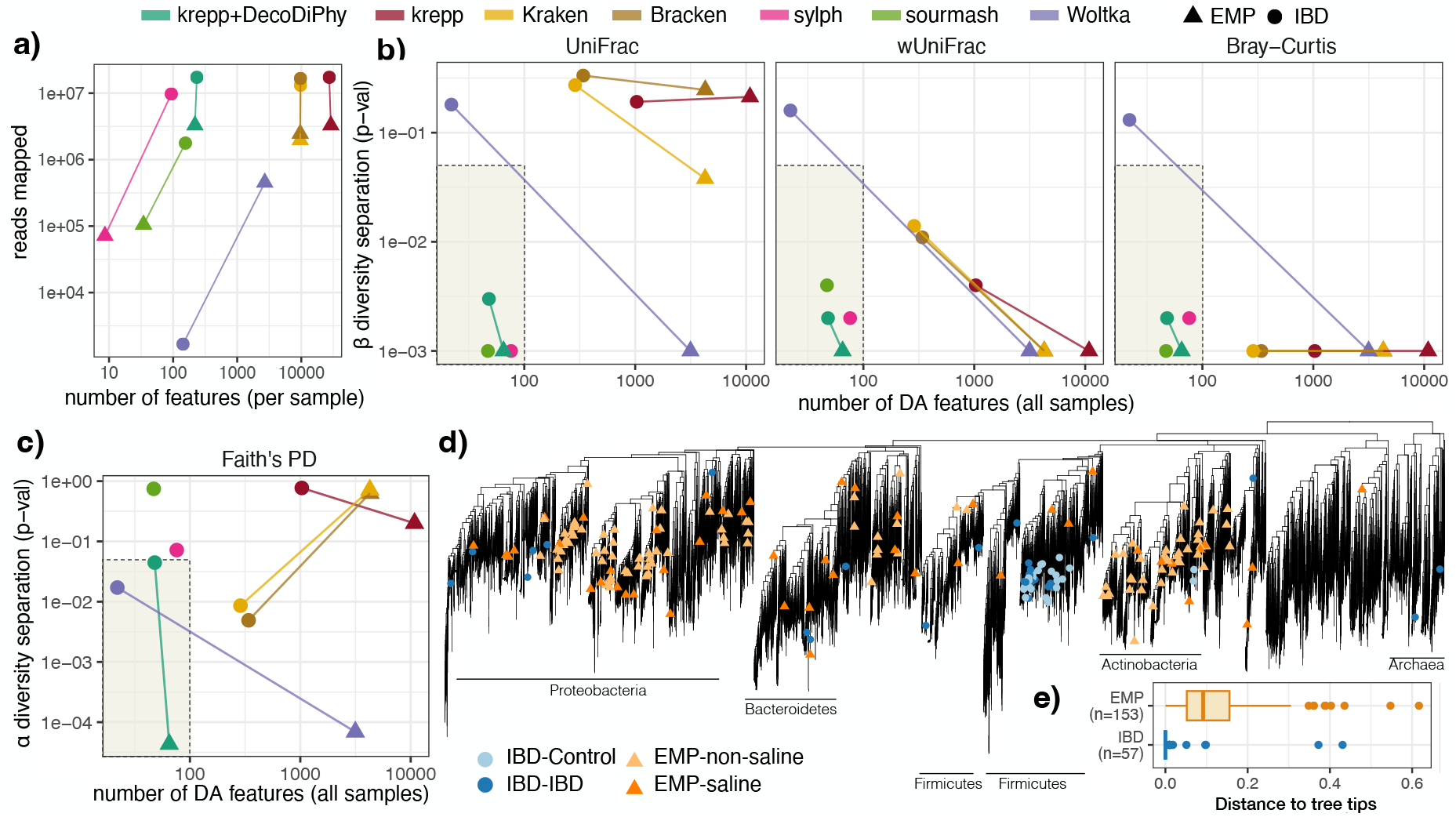
IBD and EMP datasets. **a)** The average number of reads in each sample that were assigned to a reference and the average number of features (branches with placements). For DecoDiPhy, the number of mapped reads is the same as krepp, which it uses. **b**,**c)** The number of total features marked as differentially abundant (DA) across categories (healthy/diseased for IBD, saline/non-saline for EMP), versus the *p*-value testing separation between the categories according to **(b)** *β* diversity metrics and the PERMANOVA test, and **(c)** *α* diversity using the Wilcoxon rank-sum test. Bray-Curtis *β* diversity metric only takes into account feature abundances, UniFrac only focuses on phylogenetic distance between a pair of placements, and weighted UniFrac (wUniFrac) captures both. While *ε* diversity metrics compare two samples, *α* diversity metrics quantify the diversity within one sample. For Kraken and Bracken, we use taxonomy with all branches set to 1 for (w)UniFrac and Faith’s PD. DA fails for sourmash and sylph on EMP due to too few placements recovered. Shaded: desired regime (p-value *<* 0.05) with few features that separate categories. **d)** The DA features for each dataset marked at the tip of each branch on the 16k-taxon WoL2 reference tree. Light and dark colors show the direction of association for each dataset. **e)** Distribution of the distance from each DA feature to leaves of the tree (mean); most IBD features are leaves, while for EMP are deeper in the tree.

To assess the utility of these features for downstream analysis in the absence of ground truth, we tested whether different methods could identify a compact set of differentially abundant (DA) features on a training cohort that generalizes to held-out test samples. DecoDiPhy was the only approach that achieved statistically significant separation across all tested *β* and *α* diversity measures (Fig. 3b,c) on *both* datasets. On the IBD dataset, DecoDiPhy identified 48 DA features on the training set, comparable in number to tree-agnostic joint profiling approaches (sylph and sourmash), yet achieved stronger and more consistent separation of traits (healthy/diseased) than read-level placements alone (krepp and Woltka) and read taxonomic classifiers (Kraken and Bracken). Here, krepp identified the highest number of DA features (1032), followed by Kraken and Braken, which had substantially fewer features than krepp, but much higher than joint profiling tools and DecoDiPhy (287 and 340 for Kraken and Bracken, respectively; Fig. S12). Even when the number of features used was kept constant by picking the same number of features as DecoDiPhy for other methods, separation remained significant according to UniFrac only for DecoDiPhy (Fig. S13).

Performance differences were amplified on the more novel EMP dataset. Similar to simulations, joint profiling methods identified few or no features for a substantial fraction of samples, preventing reliable DA analysis (Fig. S12). The lack of matches is not surprising as these methods were not designed for a dataset with high phylogenetic novelty. Among the remaining methods, DecoDiPhy achieved strong separation between saline and non-saline environments using only 65 DA features, whereas alternative approaches required orders of magnitude more features and showed degraded performance on abundance-free phylogenetic metrics. These results emphasize that sample-level phylogenetic consolidation enables robust, interpretable separation across biological datasets with markedly different novelty profiles.

### Differential abundance reveals dataset-specific phylogenetic resolution driven by novelty

We next asked whether the differentially abundant (DA) features identified by DecoDiPhy reflect intrinsic differences between datasets with low versus high reference novelty. To isolate the dataset-specific signal, we performed differential abundance analysis on the combined training and test cohorts for each dataset. This yielded 57 DA features for the IBD dataset and 153 for the EMP dataset.

Despite comparable sample sizes, the phylogenetic distribution of DA features differed strikingly between datasets. In the IBD cohort, DA features were concentrated within a small number of clades of the 15,953-taxon reference tree, whereas EMP features spanned substantially broader subtrees (Fig. 3d). Consistent with their origin from the human gut, IBD-associated features were largely restricted to the phyla Firmicutes, Proteobacteria, Bacteroidetes, and Actinobacteria, all of which are well represented in the reference and have established links to gut-associated microbial communities and inflammatory bowel disease [50].

In contrast, DA features in the EMP dataset occupied deeper phylogenetic branches and encompassed more diverse regions of the tree (Fig. 3e), reflecting the higher novelty and incomplete reference coverage characteristic of free-living microbial communities. Moreover, 77% of IBD DA features corresponded to terminal branches (genome-level placements), compared to only 7% in the EMP dataset. Together, these results show that DecoDiPhy naturally adapts the phylogenetic resolution of features based on reference representation and dataset novelty.

### Consolidation reveals dysbiosis patterns and recovers known IBD-associated taxa for IBD

We next asked whether the features obtained through consolidation with DecoDiPhy capture characteristic patterns of dysbiosis associated with IBD [51]. Examining the *α*diversity measure, we observed that typical IBD samples have lower than average diversity, with a tail of abnormally high diversity, consistent with dysbiosis (Fig. S14a). The number of DecoDiPhy placements (*k*) is between 100 and 400 in most cases. Moreover, there is a strong difference (p-value ≈ 0.001 by a Welch two-sample t-test) between IBD and control samples (Fig. S14b), with IBD samples showing lower *k*, pointing to lowered diversity. Impact of diversity can also be observed by comparing abundance-free UniFrac and the abundance-weighted UniFrac (wUniFrac) *β* diversity metrics (Fig. 4a). The strongest separation between healthy and IBD samples was achieved by differentially abundant (DA) features using the abundance-agnostic UniFrac metric (Fig. 4ab). However, without DecoDiPhy, UniFrac had the worst performance, especially with DA features (Fig. S14c). These results indicate that, within the consolidated feature space, differences in the presence/absence of phylogenetic lineages (rather than their abundance) are informative for disease discrimination, a pattern that is lost when *β*-diversity is measured using noisy placements.

**Figure 4:**
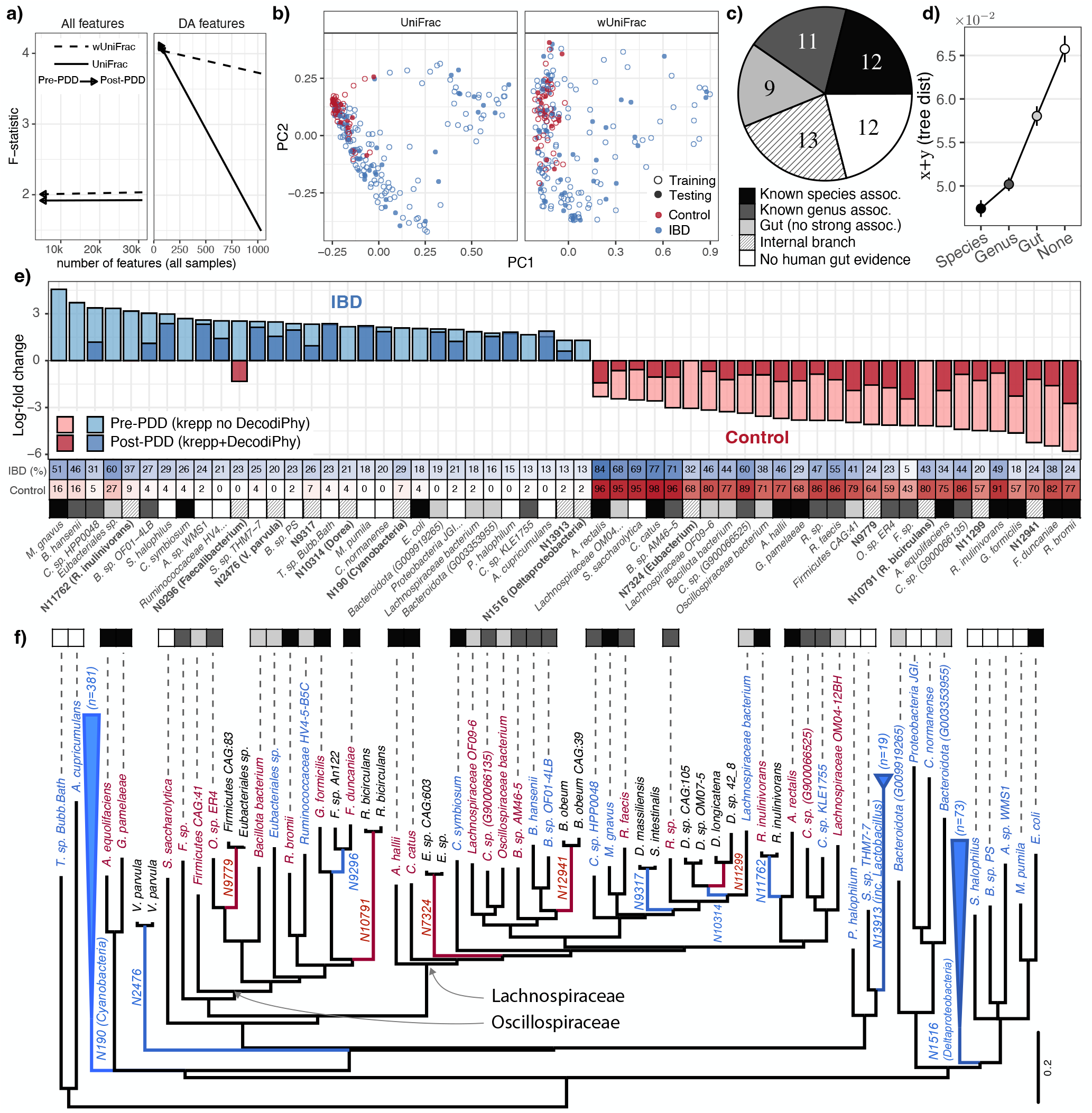
IBD dataset. **a)** The total number of DA or all features versus the pseudo-F test statistic on test samples. Higher pseudo-F corresponds to stronger separation. DA features are selected based on the training set. **b)** Principle coordinates of UniFrac and wUniFrac for DecoDiPhy, showing separation between IBD and control using DA features obtained on the train set. **c)** Distribution of prior evidence for each of the 57 combined DA features. Combined DA features correspond to the DA features obtained by combining training and test sets. **d)** Average tree distance between DecoDiPhy placement and the reference taxa per each category of prior evidence. Average is over the features in each category and all samples. Only terminal branches are shown. **e)** Log-fold change (LFC) in abundance of combined DA features; LFC of krepp is shown if available. For each feature, the proportion of IBD and control samples containing that feature in non-zero abundances are shown. For the DA leaves, we show whether they are supported by the literature (see also Table S4). **f)** The tree of all DA features. For internal features with more than 10 leaves in their subtree, the size of the subtree below them is shown in the figure.

Consistent with this observation, analysis of presence–absence patterns revealed a marked asymmetry between health- and disease-associated features (Fig. 4e). Features enriched in healthy controls tended to be present in a larger fraction of both control and IBD samples, albeit often at low abundance in IBD. In contrast, features enriched in IBD were generally less prevalent overall and were frequently absent from healthy controls. This pattern aligns with established models of gut dysbiosis in IBD, in which disease can arise through both *loss-of-function* (loss of widespread beneficial taxa) and *gain-of-function dysbiosis* (the expansion of less common, potentially pathogenic or pro-inflammatory taxa)[51].

We next examined the literature to assess the biological validity of the differentially abundant (DA) features identified by DecoDiPhy (Fig. 4c–e). Most DA features were assigned to two well-studied families, *Lachnospiraceae* and *Oscillospiraceae*, both of which are widely implicated [52–55] in gut homeostasis and inflammatory bowel disease (Fig. 4f).

Among 44 genome-level DA features, 12 species have prior evidence of association with IBD in the same direction observed here (Fig. 4c; Table S4). Of these, eight species were depleted in IBD (*Ruminococcus bromii, Faecalibacterium duncaniae, Gemmiger formicilis, Roseburia inulinivorans, Adlercreutzia equolifaciens, Gordonibacter pamelaeae, Anaerobutyricum hallii, Coprococcus catus*, and *Agathobacter rectalis*), while three species (*Mediterraneibacter gnavus, Clostridium symbiosum*, and *Escherichia coli*) were enriched. Several of these species were also detected by joint profiling methods; however, three validated species were not present among the 1,772 DA features identified prior to consolidation, indicating that consolidation can recover biologically relevant signal that is diluted at the read level without PDD.

In addition to genome-resolved features, DecoDiPhy identified 11 unclassified species from genera with well-documented roles in IBD. For these features, the direction of association and log-fold changes were consistent with those obtained by other methods when detected (Fig. S15). As expected, taxa from genera such as *Faecalibacterium, Ruminococcus, Oscillibacter, Clostridium*, and *Blautia* were identified as depleted in IBD, consistent with prior reports [56–59], while four taxa from genera *Coprococcus, Blautia*, and *Clostridium*) were enriched. *Blautia* and *Clostridium* genera are known to exhibit cohort-dependent behavior in IBD, since they contain both anti-inflammatory and pro-inflammatory members [60–62]. While *Coprococcus* is generally reported as depleted in IBD, the specific species identified here (*Coprococcus sp. HPP0048*) has been reported to increase in related postinfection irritable bowel syndrome cohorts, providing a plausible context for its enrichment [63].

Beyond genome-level features, DecoDiPhy identified three internal edges corresponding to subtrees of at least ten genomes that were enriched in IBD (Fig. 4f). One such feature represents a long branch containing a tightly clustered group of *Lactobacillus* genomes. Although *Lactobacillus* species are often regarded as beneficial commensals or probiotics, several studies have reported increased abundance of specific *Lactobacillus* strains in IBD cohorts [55, 64, 65]. A second internal feature corresponds to the class *Deltaproteobacteria*, which includes sulfate-reducing bacteria such as *Desulfovibrio* and *Desulfomicrobium*, taxa repeatedly associated with mucosal inflammation and IBD [66, 67]. The third feature maps to the least common ancestor of the phylum *Cyanobacteria*, for which evidence of association with IBD remains limited and indirect [57, 68].

Notably, the phylogenetic distance produced by DecoDiPhy between DA placements and their corresponding reference genomes (**x**_*q*_ + **y**_*q*_) was correlated with prior evidence of IBD association, suggesting that more distant placements may reflect missing or underrepresented taxa in the reference tree that are captured through placements on related branches (Fig. 4d; see also Fig. S16). Together, these results show that consolidation recovers known IBD-associated taxa while also exposing higher-level phylogenetic signals that are not accessible through genome-level features alone.

### Consolidated features capture salinity-associated structure in free-living communities

We next asked whether DecoDiPhy provides comparable interpretability on the EMP dataset by identifying features associated with salinity in free-living environments. Differential abundance analysis of DecoDiPhy placements identified 153 features, of which 106 were enriched in non-saline and 47 in saline samples (Fig. 5a).

**Figure 5:**
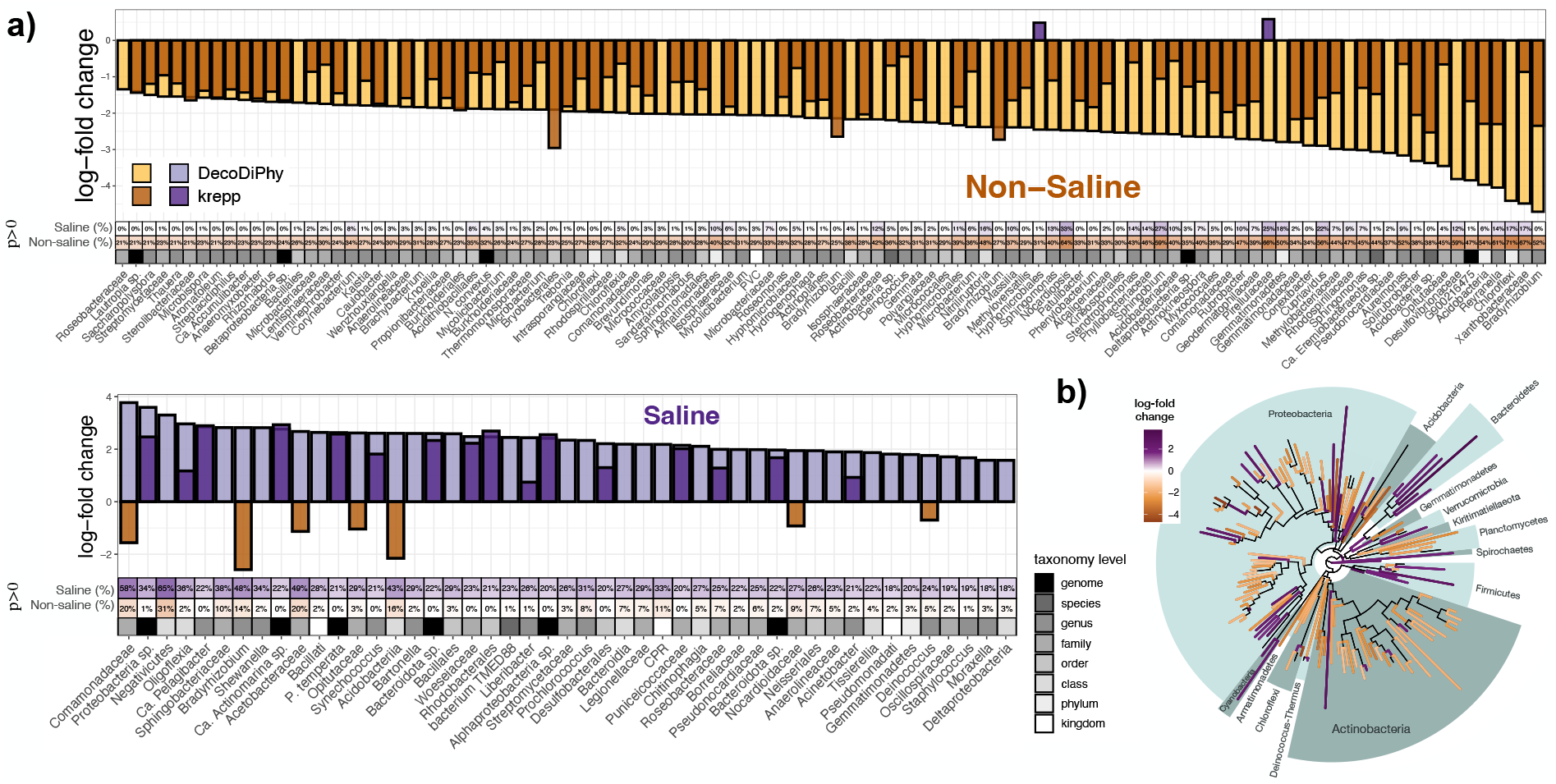
EMP dataset. **a)** Log-fold change (LFC) in abundance of combined DA features; LFC of krepp is shown if available. Combined DA features correspond to the DA features obtained by combining training and test sets. For each DA feature, we show the proportion of saline and non-saline samples with non-zero abundances for that feature. We also show the corresponding taxonomic rank for each feature. **b)** The WoL2 reference tree is restricted to the combined DA features. Colors indicate the log-fold change and the direction of association for each feature. Present phyla are highlighted in the phylogeny.

This imbalance is consistent with prior observations that microbial diversity tends to decrease with salinity, as elevated salinity imposes strong physiological constraints that limit community membership [69, 70].

The identified features span multiple taxonomic levels, ranging from species to kingdoms, and show strong group specificity: features enriched in saline environments were largely absent from non-saline samples, and vice versa. Among non-saline–associated features, *Bradyrhizobium*—a genus commonly found in forest and agricultural soils [71]—and the family *Xanthobacteraceae*, which is prevalent in freshwater and sedimentary environments [72], exhibited the largest abundance shifts relative to saline samples. Conversely, saline-associated features including *Comamonadaceae*, a family previously reported to be more abundant in saline and brackish habitats [73], as well as an unclassified *Proteobacteria* species (*Proteobacteria bacterium JGI 0000113-P07*), showed the largest increase in abundance compared to non-saline samples. This organism originates from a study of microbial gene expression in oxygen-minimum zones of the Northeast subarctic Pacific Ocean and was present in 34% of saline samples, consistent with its marine origin.

At higher taxonomic levels, several phyla showed strong salinity-specific associations (Fig. 5b). Branches from *Spirochaetes, Firmicutes, Bacteroidetes*, and *Cyanobacteria* were predominantly associated with saline environments, whereas *Actinobacteria, Acidobacteria, Planctomycetes*, and *Armatimonadetes* were largely associated with non-saline samples. These patterns are consistent with prior reports linking these phyla to saline or non-saline habitats, respectively [74–79] (Table S5).

Finally, we observed a systematic difference in the phylogenetic depth of salinity-associated features. DA features enriched in saline samples tended to occur at greater phylogenetic heights (mean height = 0.17), whereas features enriched in non-saline samples were, on average, closer to the leaves of the reference tree (mean height = 0.10; Fig. S17b). We also observed that the number of DecoDiPhy placements (*k*) is significantly higher for non-saline samples than saline samples (p-value *<* 0.001; Fig. S17c). Notably, this difference in phylogenetic height and the number of features suggests that metrics sensitive to phylogenetic diversity may be particularly informative for distinguishing saline and non-saline communities. Consistent with this interpretation, DecoDiPhy achieved the strongest separation among all methods based on sample phylogenetic diversity (Faith’s PD), indicating that consolidation enhances the recovery of phylogenetically structured signal relevant to environmental differentiation (Fig. 3c).

## Discussion

We introduced phylogenetic distance deconvolution (PDD), a framework for decomposing biological mixtures that combines two capabilities not previously integrated: phylogenetic placement and joint characterization of reads. Given the availability of many well-established tools, one may ask, what is the benefit of DecoDiPhy? Compared to taxonomic assignment, phylogenetic placement provides finer resolution [12], and unlike methods that assign reads only to leaves [12, 17, 41], PDD allows insertion on internal edges, a feature that was highly consequential on the EMP dataset and to a lesser degree on IBD. The phylogenetic output of DecoDiPhy can be translated to taxonomic profiles, which showed high levels of purity while retaining completeness on CAMI-II. In our limited exploration, it outperformed *k*-mer-based and most alignment-based methods across most taxonomic levels, even with the simple LCA mapping we used.

A separate concern is whether reads should be jointly analyzed. Methods for read-level phylogenetic placement (krepp), taxonomic classification (e.g., Kraken), or genome mapping (e.g., bowtie2/Woltka) produce over-dispersed placements, adding noise and reducing interpretability. DecoDiPhy consolidates over-dispersed placements into a smaller number of informative phylogenetic features while retaining phenotype separation. When fewer features are as powerful in distinguishing samples as more features, the reduced set increases the interpretability of the results; the small subset may also enable testing hypotheses generated through metagenomics in follow-up experimental analyses. While a compact set of features can be obtained from any method by selecting the top features based on a threshold, we observe that even when using the same number of features, DecoDiPhy achieves better separation significance, especially according to UniFrac (Fig. S13).

Methods that jointly model reads (e.g., sylph, sourmash, Quikr) enjoy the same benefit of having compact set of features, but suffered on highly novel data. Compared to these, we expected and observed that PDD performs better when the input is phylogenetically novel relative to the reference. On datasets with limited novelty, such as human gut, sourmash, and sylph, perform well and in some cases match or outperform DecoDiPhy. In contrast, on more phylogenetically novel data (e.g., EMP), these methods fail to find any matches for many samples and very few for others, reducing their ability to profile the samples. This is expected because these methods are not designed to identify novel species in metagenomic samples.

While PDDs are more robust to novelty than non-phylogenetic methods, completeness of reference trees continues to matter. Increased novelty does increase error for all methods, including DecoDiPhy, on simulations (Fig. 2d). The high-novelty scenario in our simulations captures 90% novelty at the species level (Fig. S3c). However, if the tree is missing entire groups (e.g., viruses or eukaryotes) that are present in the sample, those species cannot be placed. Moreover, the prevalence of internal branches in outputs for novel samples (e.g., in the EMP dataset; see 5) makes actionable experiments more challenging. Similarly, while most features detected in the IBD dataset are consistent with prior studies, additional internal and terminal features with limited existing annotation highlight targets for future biological investigation and expose gaps in current reference collections. When such features show high distance to samples (Fig. S16a), our results highlight a lack of sampling in the reference set. Thus, using more complete references remains desirable even with DecoDiPhy.

DecoDiPhy is designed to be scalable (quadratic with the size of the reference tree; see Fig. 2a); however, it is still more expensive to run than some of the alternative state-of-the-art. The running time of sylph per sample is far better than all other methods (Table S6) and outperforms krepp+DecoDiPhy by more than two orders of magnitude. Kraken2 and sourmash are slower than sylph but still by one to two orders of magnitude faster than krepp. DecoDiPhy adds only 17% to the running time of krepp. By far the slowest method is Woltka, used with bowtie2, which is an order of magnitude slower than krepp+DecoDiPhy. Overall, results point to a substantially higher running time for DecoDiPhy, but not to the extent that analyses of datasets with tens of thousands of references become impossible. Further scaling to many hundreds of thousands of references will require further heuristic improvements.

Our PDD framework raises theoretical and algorithmic questions about identifiability, robustness to distance error, and the optimality of search strategies, which can be addressed in future work. Improving scalability through alternative optimization approaches or better heuristics may be possible. Here, we used a single tree as a reference, but prokaryotic evolution exhibits high rates of horizontal gene transfer (HGT), which the current method does not account for. HGT can create incongruence between the metric spaces for various parts of the genome, making the use of a single tree imprecise. Other processes, such as simple rate variation across genes, can also have the same impact. Our future work will explore the theoretical impacts and try to incorporate such complexities by allowing the use of multiple (gene) trees or a reference network. We note that our current results (both on simulated and real metagenomes) are based on real genomes that have experienced complexities such as HGT and rate variation, and thus, their impacts are already reflected in our empirical results.

Several practical extensions may further increase the utility of PDD. Adaptive strategies for selecting the number of mixture components, incorporation of genome coverage as a regularization term, and integration with alternative distance estimators (e.g., Containment Jaccard [80]) could improve performance in specific applications. We used a simple criterion to choose *k* to maintain generalization; however, a more complex criterion may perform better for specific types of samples, a topic to be explored in the future. We anticipate that combining PDD with advances in distance estimation and reference construction will enable more accurate and interpretable analysis of complex biological mixtures. Finally, although we focused on metagenomics, the PDD framework is applicable to other mixture problems defined on phylogenies, including identifying progenitor species of hybrids [4, 43], distance-based admixture analysis on population trees [3], and deconvolving cancer samples relative to existing ones [2]. In each of these applications, mixed data need to be identified with respect to a phylogeny, and thus, PDDs may prove useful.

## Conclusion

PDD provides a general framework for decomposing biological mixtures on phylogenies by jointly modeling reads and allowing placements on both terminal and internal branches. Across simulations and empirical datasets, DecoDiPhy produced accurate and interpretable representations of mixtures, particularly in settings with substantial phylogenetic novelty where existing approaches were less effective. We anticipate that combining PDD with continued advances in distance estimation and reference construction will enable more accurate and interpretable analysis of complex biological mixtures and broaden the use of phylogeny-aware mixture deconvolution across biological domains.

## Methods

### Notation and Problem Formulation

Let *T* = (*V*_*T*_, *E*_*T*_) be a tree with nodes *V*_*T*_ and edges *E*_*T*_ . For a node *v* ∈ *V*_*T*_, we let *C*_*T*_ (*v*) denote the set of all the leaves in the subtree below *v*, and *e*(*v*) denote the incoming edge to *v*. For any edge *e* ∈ *E*_*T*_, let *l*_*T*_ (*e*) denote the length of the branch *e* in *T* . We let *d*_*T*_ (*a, b*) be the path length between two nodes *a, b* ∈ *V*_*T*_ . We often omit the subscript *T* when it is the reference tree *R*, defined below.

Consider a phylogenetic tree *F* on a leaf set labeled by *R* ∪ *Q* where *Q* represents a set of *k* query taxa and *R* represents a set of *n* reference taxa, indexed by [*k*], [*n*] = {1,. .., *n*}, resp. We assume such a tree *F* exists, though we do not have access to it. Instead, we have access to a reference tree *R*, which we assume is *F* induced down to reference leaves *R* (i.e., by pruning *Q* from *F* and suppressing degree-2 nodes). We refer to *R* as the *reference tree* and *F* as the *full tree*. We let *m* ≤ 2*n* − 2 denote |*V*_*R*_|. We assume *F* is rooted arbitrarily and *R* keeps the same arbitrary rooting. Note that while we assume *Q* ∩ *R* = ∅, branch lengths of *F* can be arbitrarily short; thus, if the query *Q* includes an organism *q* that is exceedingly similar to some reference *r* ∈ *R*, we can include *r* and *q* in *F* with arbitrarily small branch length separating them. In addition to *R*, we have access to some data points (e.g., reads) *X* gathered from a *phylogenetic mixed query* (*Q*, **p**), where each *q* ∈*Q* is associated with an abundance **p**_*q*_ ≥ 0 such that 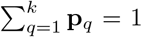 (i.e., **p** ∈ Δ^*k−*1^ where Δ denotes a (*k* − 1)-simplex). Each *x* ∈ *X* belongs to an (unknown) constituent *q* ∈ *Q*, and the proportion of *X* belonging to *q* follows **p**_*q*_ (i.e., *X* can be modeled as a random draw according to **p**). Let the query-reference distance matrix *D*(*F*) ∈ ℝ^*n×k*^ be defined by *D*(*F*)_*r,q*_ = *d*_*F*_ (*r, q*). We define the *(true) mixture distance* vector **d** ∈ ℝ^*n*^ as **d** = *D*(*F*) *·* **p** (Fig. 1a). If we do not have access to **d**, we assume we can estimate 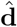 from *X* such that (ideally) 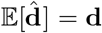. The goal is to recover *F* and **p** from *R* and either **d**, or 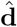. Note that in this general form, *F* can have splits (i.e., unrooted edges) made entirely of subsets of *Q*. Placing a query *q* on an edge *e* = (*u, v*) ∈ *E*_*R*_ results in dividing *e* into two new edges *e*_1_ = (*u, w*) and *e*_2_ = (*w, v*) by adding a new node *w*, and adding *q* as a child to *w*. We call *e* the *true placement edge* of *q* on *R* if the resulting tree is compatible with *F* (see [81] for the definition of compatibility). We also define the distal (**x**_*q*_) and pendant (**y**_*q*_) length of *q* on the resulting tree as the length of *e*_2_ relative to the length of *e* 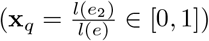, and the length of the edge (*w, q*), respectively (see Fig. S1a).

We begin with the case where the true **d** is given to study the identifiability of the problem.

#### Problem 1

(Phylogenetic Distance Deconvolution (PDD)). *Given a reference tree R on R* = [*n*], *a mixture distance vector* **d** ∈ ℝ^*n*^, *and k, find a full tree F with n* + *k leaves that is compatible with (i*.*e*., *induces) R and includes k additional query leaves and their proportions* **p** ∈ Δ^*k−*1^ *such that*

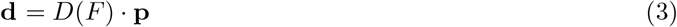

Even with error-free input **d**, for *F* to be recoverable, we need identifiability; that is, if (*F*, **p**) satisfies Eq. (3), there should exist no other (*F* ^*′*^, **p**^*′*^) that also satisfies it. However, PDD is only partially identifiable.

#### Claim 1.

*If two queries q and q*^*′*^ *share the same placement edge on R, both queries can be replaced with a single query with abundance* 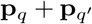 *without changing the distance mixture* **d**.

#### Claim 2.

*Problem 1 is not identifiable if* **x**_*q*_ = 0 *or* **x**_*q*_ = 1 *are allowed. Thus, we only allow* **x** ∈ (0, 1)^*k*^. *Also, any two vectors* **y** *and* **y**^*′*^ *with* 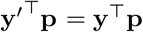 *lead to the same distance mixture* **d** *in Problem 1. Thus, we can only identify the mean terminal branch length* 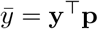 *and not the individual values of* **y**.

#### Claim 3.

*Even for fixed (true) placement edges, the relative positions x*_*i*_ *and proportions p*_*i*_ *are unidentifiable under some corner conditions for queries placed on adjacent edges*.

All claims are proved in supplementary Section C.6. Because of these negative results, we need to disallow two queries being placed on the same edge of *R*. Essentially, we can identify the total **p** subtending from each branch of *R*, but cannot deconvolve the relationship per branch. More formally:

#### Assumption 1.

*No two queries share a placement edge. Thus, a solution to Problem 1 can be expressed as k distinct placements on R represented by vectors* **e, x, p** *indexed by* [*k*] *and* 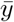; **e**_*q*_ ∈ *E*_*R*_ *indicates the placement edge*, 0 *<* **x**_*q*_ *<* 1 *is the distal length of q, and* **p**_*q*_ *is its abundance (Fig. 1a). We only seek* 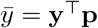 *and not* **y**.

Even with Assumption 1, the problem is still *not always* identifiable, as shown by a counter-intuitive counterexample, showing two very different query sets with identical **d** (Fig. S1f). Note that the lack of identifiability requires exactly identical branch lengths and very specific proportions. Small deviations from this strict symmetry would make the problem identifiable. Given expected errors in inputs to PDD, identifiability should be considered a theoretical question, not a practical one. Despite the lack of strict identifiability, we empirically observed that conditions that break identifiability rarely happen empirically and when they do, they seem to be explained by floating point numerical resolution (Table S2 and Fig. 2b). We leave a full theoretical characterization of identifiability to future work.

Problem 1 can be expressed linear algebraically using auxiliary variables. For a query leaf *q* with placement edge **e**_*q*_ = (*u, v*), distal length **x**_*q*_, and pendant length **y**_*q*_, the tree distance between *q* and any leaf *r*_*i*_ ∈ *C*(*v*) can be written as *d*(*q, r*_*i*_) = *d*(*v, r*_*i*_)+ *l*(**e**_*q*_) *·* **x**_*q*_ + **y**_*q*_ (see Fig. S1b). Similarly, for any leaf *r*_*o*_ ∉ *C*(*v*), we have *d*(*q, r*_*i*_) = *d*(*v, r*_*i*_) − *l*(**e**_*q*_) *·* **x**_*q*_ + **y**_*q*_ (Fig. S1c). We combine these distances with Eq. (3) as follows. Let *A* : (*V*_*R*_)^*k*^ ⟶ {0, 1}^*m×k*^ be a binary representation of **e** defined by *A*(**e**)_*v,q*_ = 1 iff query *q* is placed above node *v* (i.e., *e*(*v*) = **e**_*q*_). Note that ∀*q* : ∑_*e∈E*_ *A*_*e,q*_ = 1 and ∀*e* : ∑_*q*_ *A*_*e,q*_ ≤ 1, where the second constraint follows from Assumption 1. We let the *anchor node a*_*q*_ ∈ *V*_*R*_ of *q* be the tail of its true placement edge. Let **w** ∈ (0, 1)^*k*^ be **w** = **x** ∘**p** where ∘ is element-wise product and note that **w**_*q*_ ≤ **p**_*q*_. Let *C* ∈ {−1, 1}^*n×m*^, where *C*_*r,v*_ = 1 iff *r* ∈ *C*(*v*) (*r* descends from *v*) and *C*_*r,e*_ = −1 otherwise; *L* ∈ ℝ^*m×m*^ = diag(*l*(*e*(*v*_1_)),. .., *l*(*e*(*v*_*m*_))) where *V*_*R*_ = *{v*_1_ .. . *v*_*m*_}. Finally, *D* ∈ ℝ^*n×m*^ is a distance matrix equivalent to tree *R* where *D*_*r,v*_ = *d*_*R*_(*r, v*) for *r* ∈ *R,v* ∈ *V*_*R*_.

#### Claim 4.

*Under conditions of Assumption 1, we can rewrite Eq*. (3) *as*

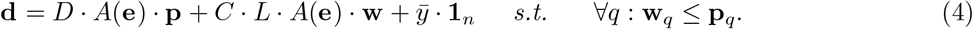

We can rewrite Problem 1 using Eq. (4), noting that *D, C, L* are given and *A*(**e**), **w, p** 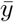 are unknown variables. **e** gives placement edges. Attachment positions on each can be recovered as **x**_*q*_ = **w**_*q*_*/***p**_*q*_, provided **p**_*q*_ *>* 0. While we can only recover 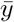 (Claim 2), in practice, we can use heuristics to estimate individual **y**_*q*_ values (see Section C.5).

On real data, due to noise in 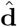, Problem 1 is not sufficient. To accommodate error, we form an optimization problem. We need a metric to quantify the similarity of two distance vectors. A natural choice is *ω*_2_, which is easy to optimize. We can now define:

#### Problem 2

(PDD optimization). *Given a reference tree R and an estimated mixture distance vector* 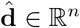, *infer a multiplacement* (**p, e, w**, 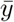*) that adds k queries to R by minimizing:*

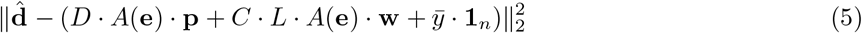

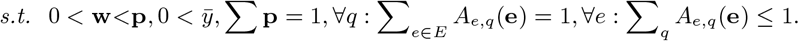

Among variables of Problem 2, *A*(**e**) and *k* are combinatorial in nature and hard to optimize directly.

### DecoDiPhy Algorithm

#### Exact solution

If we fix *A* in Problem 2, we get a “small-problem” of minimizing (**p, w**,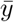) in Eq. (5), which is a simple quadratic objective function due to the linearity of Eq. (4). Since the objective function is convex, the small-problem has a unique minimization cost; however, the objective function is not strictly convex and can have multiple minimizers. Nevertheless, convexity of Eq. (5) allows an exact algorithm:

##### Definition 1

(Exhaustive search). *For every possible k, for every possible matrix A, optimize the quadratic objective function of Eq*. (5) *using gradient descent; choose* (*k, A*, **p, w**, 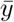) *that minimizes objective value*.

This exhaustive algorithm is not practical. Even if we limit ourselves to a constant range 2 ≤ *k* ≤ *K*, we need to explore 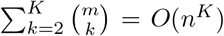possible *A* matrices. Ignoring the time for each optimization, we will need *O*(*n*^*K*^) steps, which is practical only for small *K* (e.g., *K* = 3), even for moderately sized trees. A further complication is that on noisy data, as *k* increases, the objective function tends to decrease. Thus, we cannot simply rely on the optimization score to choose the right *k*; instead, we need a heuristic to select *k*.

#### Heuristic search

We developed a heuristic search algorithm (Algorithm S1) adopting a greedy search with local refinements. Briefly, we add one placement at a time, starting from *k* = 1 (Line 15 in Algorithm S1). For each *k*, we start from the optimal solution found for *k* − 1, and test all possible placement edges (Line 3) for the *k*th query (Line 6), solving the small-problem (i.e., quadratic optimization) to minimize Eq. (5) for each edge visited and selecting the placement with the minimum residue. We then revisit all prior *k* − 1 queries and test if their placement can improve (Line 18), fixing all other queries; we accept any move that reduces the residue. When revisiting prior nodes, we search a constant number of edges (Line 5), limiting the search to edges that are up to *ρ* edges away from the best previous placement (i.e., limited radius). We stop for any *k* only when no improvement is possible. Given the finite number of possible *k*-placements and the optimality of the small-problem solutions, this approach guarantees convergence to a global minimum within the search space defined by the radius *ρ*. Also, note that with this search strategy, no two placements will move at the same time; therefore, the optimizer will never explore different permutations of the same set of placements that would have identical scores. The first small problem for a given *k* is initialized with 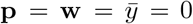, whereas subsequent small problems are initialized using the estimated values of **p, w**, and 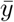from the previous small problem; therefore, the DecoDiPhy algorithm does not rely on any random initialization. We use Algorithm S2 to handle boundary cases of Claim 2. For large backbones (*n >* 10^3^), we use divide-and-conquer (supplementary Section C.4) to solve PDDs on several subtrees.

#### Selecting *k*

Let us examine an example dataset (Fig. S1d). With *k* = 2, some placements are correct, and others are phylogenetically close, as reflected in low weighted UniFrac error. The observation that selecting fewer edges than *k* still leads to partially correct results motivated our semi-greedy heuristic. All metrics improve as *k* approaches its true value. When we reach the true *k*, the residue drops dramatically. However, this pattern becomes blurred with noisy data (Fig. S1e), so we do not use a loss-based criterion by default. Instead, note that for *i > k*, the extra placements tend to lead to many placements *q* with **p**_*q*_ ≈ 0. Thus, we stop exploring higher *k* when at least one **p**_*q*_ ≈0, defined as 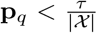(default *τ* = 1000). Intuitively, this stopping criterion will tend to avoid adding placements if they explain fewer than *τ* reads in the query set *χ*; therefore, it automatically adjusts the minimum allowed abundance as sequencing depth changes. While the number of inferred placements by DecoDiPhy depends on the choice of the stopping criterion and *τ*, accuracy and error of inferred placements are more robust to this selection based on most metrics (Fig. S9a,b). Results on the real datasets (IBD and EMP) also show that as the number of placements increases, the solution converges to a final set of placements with respect to wUniFrac (Fig. S11), with additional placements only contributing low abundances, local moves, or both, in ways that lead to low wUniFrac distances.

#### Asymptotic time

*D* and *C · L* can be easily pre-computed once in *O*(*n*^2^) time, and multiplying with *A*(**e**) is simple column selection. Because the greedy refinement is allowed to continue moving queries until no improvement is observed, the asymptotic running time is not strictly bounded. For exposition, let us bound the number of rounds of optimization in Line 18 by *C* and bound *k* (Line 15) by *K*; then, we will need ɵ (*K*) outer loops (Line 15), each with *O*(*C*) rounds of improvement, each with *O*(*K*) queries moved across a constant number of edges, and the *k*^th^ query moved across ɵ(*n*) edges. Thus, in total, we run the small-problem *O*(*C*(*K*^2^ + *Kn*)) = *O*(*CKn*) times. In our experiments, *C* is nearly constant versus *n* and *k* (Fig. S18c,d), resulting in *O*(*Kn*) runs of the small-problem, compared to *O*(*n*^*K*^) for the exact algorithm. We implemented the small-problem using the OSQP [45] package for convex optimization, which uses a first-order iterative method (ADMM).

Empirical tests show that the small problem with ɵ (*K*) variables and ɵ (*n*) equations scales with *O*(*n*^0.3^*k* + *k*^2.4^) (Fig. S18a,b). Sublinear dependency on *n* may be due to redundancies and/or sparsity in matrices and the highly optimized implementation within OSQP.

### Estimating distance vector 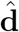

In theory, we need an unbiased estimator of the additive tree distances from *χ* however, in practice, De-coDiPhy can take any 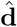 as input. In practice, we explore two methods. *i*) Sequence-based 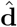: Assume genomes evolve down *F* under a Markov model such as the JC69 [44] or Felsenstein 1981 (F81) model [82]. Let *t*(*h*) be the phylogenetically corrected distances for appropriate input; for example, for JC, we can use 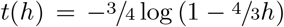 when *h*_*x,r*_ is the Hamming distance between an error-free read *x* from query *q* and a reference genome *r* if *x* is aligned correctly to *r*. If *t*(*h*_*x,r*_) is an unbiased estimator of *d*_*F*_ (*r, q*), we can easily average them across reads to obtain a consistent estimate for **d**_*r*_. When the estimator is consistent but biased, due to fixed read length, the mean of *t*(*h*_*x,r*_) over *x* ∈ *χ* does not converge to mixture distance **d**_*r*_ as |X| → ∞. The widely-used phylogenetic corrections do have a bias given finite-length reads [83]. However, these biases are relatively small (e.g.,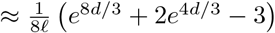 for branch length *d* and read length *l* for JC69, which is less than 1.6% of *d* for *d <* 1 with *l* = 150) and are dominated by variance (which often shrinks with 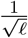). Accepting the small bias, we use the average of *t*(*h*_*x,r*_) as 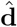 in practice, noting that the estimator’s variance and the impact of model misspecification are likely far greater than this bias. Computing *t* for JC69 is trivial and is relatively easy for F81, which allows unequal base frequencies (*t*_*F* 81_(*h*) = −(1 − *S*) log(1 − ^*h*^*/*_(1 *− S*)_), where 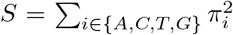 is the sum of squared base frequencies). Computing all necessary parameters for more complex models from *k*-mers alone is possible for all models nested within the TK4 model [84]; however, these make calculations far more challenging in practice. In our experiments, we explored JC69 and F81 models, simply using base frequencies for each reference genome for the F81 correction. For computing distances for genome-wide reads, we can use read alignment or the more scalable alignment-free method, krepp [22]. *ii*) Placement-based 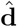: We also use PDDs to consolidate outputs of per-read placement methods. We use an existing method to place each read *x* ∈ *χ* on *R* and let 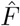 be the resulting tree, creating polytomies for multiple placements on the same branch. We simply use the mean distance 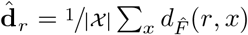. While individual placements may be wrong, if 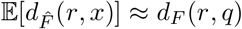 (expectation is over all *x* that belong to *q*), then 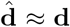. We use krepp [22] to obtain per-read placements, setting **x** = ½, **y** to the height of the sister clade. krepp can be configured to output one or multiple placements per read. We explore both modes, and for multi-placements, we take the average distance per read.

### Experiment Design

**E1:** Used to evaluate the effectiveness of our heuristic, it contains 12 biological trees with *n* ≤ 363 (Table S3). We compare Alg. S1 to the exact version (Definition 1), feasible for *k* ∈ {2, 3} and *n <* 120, and a version of Alg. S1 with true *k* given or with *ρ* set to ∞ to perform a more thorough search. We randomly selected *k* ∈ {2, 3, 5, 7, 10} query species (only keeping *k* ≤ ^*n*^*/*_3_ cases) and assigned a random abundance vector **p** ∈ Δ^*k*−1^ (Section C.7.2). The query taxa were pruned to obtain *R* (10 replicates). We tested both true distances **d** and a noisy estimate 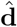 meant to reflect errors in placing individual reads on *R*. The procedure (Section C.7.2) essentially divides queries into 10^5^ pieces and moves each piece around on the tree stochastically, computing 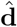 from the noisy placements. We use two settings with pieces moving by an average of 0.6 or 1.56 edges. Since no other method is specifically designed for PDDs, we compared DecoDiPhy to three baseline algorithms, all given true *k*: reverse-search (apply the small PDD on all edges, remove low **p**_*q*_ branches, and recurse), *k*-nearest leaves by distance, and a greedy *k*-nearest leaves with feedback (removing the contribution of already selected leaves before selecting the next leaf); see Section C.7.2.

**E2:** To test the impact of mixture distance calculation, we used two relatively small subtrees with *n* = 34 and 111 (see Section C.7.3). For each subtree, we simulated five sets of query taxa for *k* ∈ {3, 5, 10} together with randomly drawn abundances. We simulated 10^5^ short 150bp reads with errors from query genomes using ART [85] and ran DecoDiPhy with both true and estimated distances. For distance computation, we used both sequence-based and distance-based methods described earlier, implemented using krepp [22], which is a *k*-mer-based method that estimates read-to-reference (Hamming) distances as well as either a single placement branch or multiple equally likely placement branches (multi-placements) for each read.

**E3: Simulated metagenomes**. We simulated realistic metagenomes using leave-out experiments on the WoL-2 [46] reference tree with 15,953 leaves to compare DecoDiPhy to existing metagenomic profiling methods. Zhu *et al*. [12] have mapped 210 HMI (Human Microbiome Initiative) samples across 7 human body sites to WoL2 genomes using Woltka [12], producing a vector **p**^(*i*)^ of normalized OGU counts for each sample *i*. To mimic these microbial communities, we randomly selected 50 samples, and for each, drew a count for each genome from Multinomial(*N* ; **p**^(*i*)^) and normalized these counts to obtain the ground-truth abundances **p**. For each sample, we used *N* ∈ {10^2^, 10^3^, 10^4^} to control true *k* (low, med, high; Fig. S3a), and left out either some or all queries from *R* to create either low or high novelty levels (Section C.7.4). This process resulted in 33, 102, and 214 average queries per sample, for low, med, and high true *k*, respectively.

For each case, we simulated 10^6^ short reads with errors using ART [85] and analyzed these using krepp (multiplacements) [22] with and without DecoDiPhy. Since no other method performs phylogenetic placement on such a scale, we compared with Wotlka, which maps reads to genomes using bowtie2 [86], and sylph [41], which assigns the entire sample to genomes (Section C.7.5), treating results as placing on leaf branches of *R*. We were able to only test methods that scale to the WoL-v2 reference set, assign abundances to genomes as opposed to taxonomic units (incongruent with *R*), and can easily leave out chosen “query” genomes.

#### E4: CAMI-II analysis

We evaluated DecoDiPhy as a taxonomic profiling tool on the CAMI-II challenge [47]. We chose the ten-sample marine dataset for its high reported novelty (61% of genomes without close taxa present). We benchmarked krepp+DecoDiPhy against the *k*-mer-based methods krepp and Bracken [37], as well as other tools studied by CAMI-II, including metagenomic operational taxonomic units (mOTUs [87] and mOTUs2 [88]), Metalign [89], MetaPhlAn2 [90], CCMetagen [91], NBC++ [92], and LSHvec [93]. For krepp and DecoDiPhy, we used a reference tree with 50,752 taxa published previously [22] as input. To increase scalability of DecoDiPhy, we divided the reference tree into 32 subtrees of on average 1,586 taxa using TreeCluster [94], ran DecoDiPhy on each subtree independently, and combined all placements (see Section C.4). To obtain taxonomic profiles from the phylogenetic placement outputs of DecoDiPhy and krepp, we applied the following procedure. For each placement, we first corrected for genome lengths by dividing the abundance of each placement by the average length of the genomes under the placement edge and renormalizing the abundances to sum to one. We then assigned a taxonomic label to each placement as the lowest taxonomic rank shared among all the genomes below the placement edge and obtained a taxonomic profile at each level by summing the abundances corresponding to that level. We computed purity, completeness, weighted UniFrac, and L1 norm error for each method using the open-community profiling assessment tools (OPAL [95]). Purity and completeness are abundance agnostic (only considering presence or absence of taxonomic labels), whereas weighted UniFrac and L1 norm error take abundances into account.

#### Metrics

We used four widely used metrics from ecology to compare true and estimated multi-placements. *Jaccard index* (intersection*/*union) measures the similarity of placement edges alone, ignoring **p, x**, *y*, and *R. Bray-Curtis (BC)* [96] is similar to *ℓ*_2_ norm; it incorporates abundances **p** but ignores the tree. *UniFrac* [23] and *weighted UniFrac (wUniFrac)*take into account *R* (unshared spanned branch length over total spanned branch length); UniFrac ignores **p** while wUniFrac considers it. To compute (w)UniFrac, we place true and estimated queries on *R* but set all **y**s (but not **x**s) to 0 for true and estimated placements because the alternative methods cannot compute **y** and our method can compute only 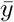.

### Biological Experiments

We studied two metagenomic datasets representing contrasting scenarios: an IBD (Inflammatory Bowel Disease) dataset with 220 human gut samples from diseased subjects and controls [48], and a subset of EMP (Earth Microbiome Project) dataset with 210 samples collected from free-living communities [49] (Section C.7.6). We analyzed both datasets using the 15,953-taxon WoL2 [46] reference set. We compared DecoDiPhy, with krepp multiplacements as input, to six methods, including those from E3 plus sourmash [39], Kraken-II [31], and Bracken [37], which we could not include in simulations because query leave-out is expensive for these methods. To evaluate accuracy, we examined the separation between diseased and control samples for IBD and between saline and non-saline free-living samples in EMP. For each dataset, samples were split into training (70%) and testing (30%) sets. Differentially abundant (DA) features were identified on the training set using ANCOM-BC [97], implemented in QIIME 2, given class labels, and selecting those with a *q*-value *<* 0.005. Note that each feature is an internal or terminal branch on the phylogeny for krepp and DecoDiPhy, a genome or terminal branch on the phylogeny for sylph, sourmash, and Woltka, and a taxonomic label for Kraken and Bracken. Pairwise sample distances on the test set were computed using either all features or the DA features based on BC and (w)UniFrac, using the taxonomic tree (all branch lengths set to 1) for Kraken and Bracken. We used Faith’s Phylogenetic Diversity metric [98] to compute the *α* diversity of each sample. We used the PERMANOVA test of whether the separation of labels is significantly better than random for *β* diversity (pairwise) metrics, and the Wilcoxon test for *α* diversity metrics. See Section C.7.6 for commands used.

## Declarations

### Ethics approval and consent to participate

Not applicable.

### Consent for publication

Not applicable.

### Availability of data and materials

The software is available at github.com/shayesteh99/DecoDiPhy. The datasets generated and/or analyzed during this study are available on GitHub github.com/shayesteh99/DecoDiPhy-Data.

### Authors’ contributions

SA, SM, and MEK developed the methodology. SA implemented the method. SA, AOBS, and ER conducted the analysis. All authors contributed to writing and revising the manuscript and approved the final manuscript.

## Acknowledgements

This project was initiated as part of the Warren Alpert fellowship to S.A. from UCLA. This work used Expanse at San Diego Supercomputing Center through allocation ASC150046 from the Advanced Cyberinfrastructure Coordination Ecosystem: Services & Support (ACCESS) program, which is supported by U.S. National Science Foundation grants #2138259, #2138286, #2138307, #2137603, and #2138296.

