## Supplementary Material for "DecoDiPhy: Deconvolving Phylogenetic Distance Mixtures"

### A Supplementary figures

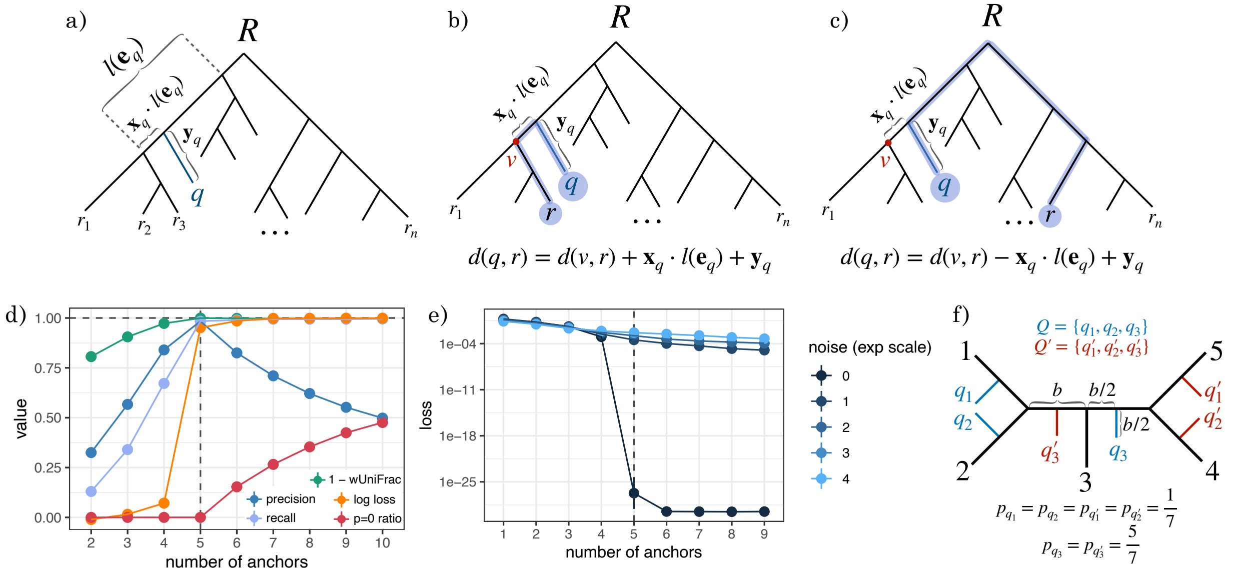

Figure S1: **a)** Definition of the placement edge $\mathbf{e}_q$, distal length $\mathbf{x}_q$, and pendant length $\mathbf{y}_q$ for a query leaf $q$. Note that $l(\mathbf{e}_q)$ denotes the length of the edge $\mathbf{e}_q$. **b,c)** An example of how the pairwise distance $d(q,r)$ is computed from the distance $d(v,r)$ when (b) $r$ is a descendant of $v$ ($r \in C(v)$) and (c) $r$ is not a descendant of $v$ ($r \notin C(v)$). **d)** Exploration of the search algorithm on an example dataset with $n = 100$ and true $k = 5$. We show the proportion of placement edges that are correct (precision) and correct placements found (recall); 1−wUniFrac, a measure of accuracy incorporating $\mathbf{p}$, $\mathbf{x}$, and $\mathbf{e}$; the normalized residue $\frac{\max_k (\log \ell_k)}{\log \ell}$ (log loss); and proportion of abundances $< 0.01$ ($p = 0$ ratio). Metrics improve for $k \leq 5$; using $k > 5$ yields many $\mathbf{p} < 0.01$ values and small changes in loss. **e)** Exploration of the search algorithm on an example dataset with $n = 100$ and true $k = 5$, and different noise levels. As noise increases, the loss at the correct $k$ increases, making any stopping criteria based on loss value fail on higher noise levels. **f)** Unidentifiable example. All $\mathbf{y}_{q_i} = \mathbf{y}_{q'_i} = \frac{b}{2}$, and $\mathbf{x}_{q_i} = \mathbf{x}_{q'_i} = \frac{1}{2}$ such that $Q$ and $Q'$ produce the same distance vector $\mathbf{d}$. Note that in this example, breaking the symmetry by changing the branch lengths, changing abundances, or adding new taxa to the backbone tree, makes the two distance vectors unequal and therefore, the placements become identifiable.

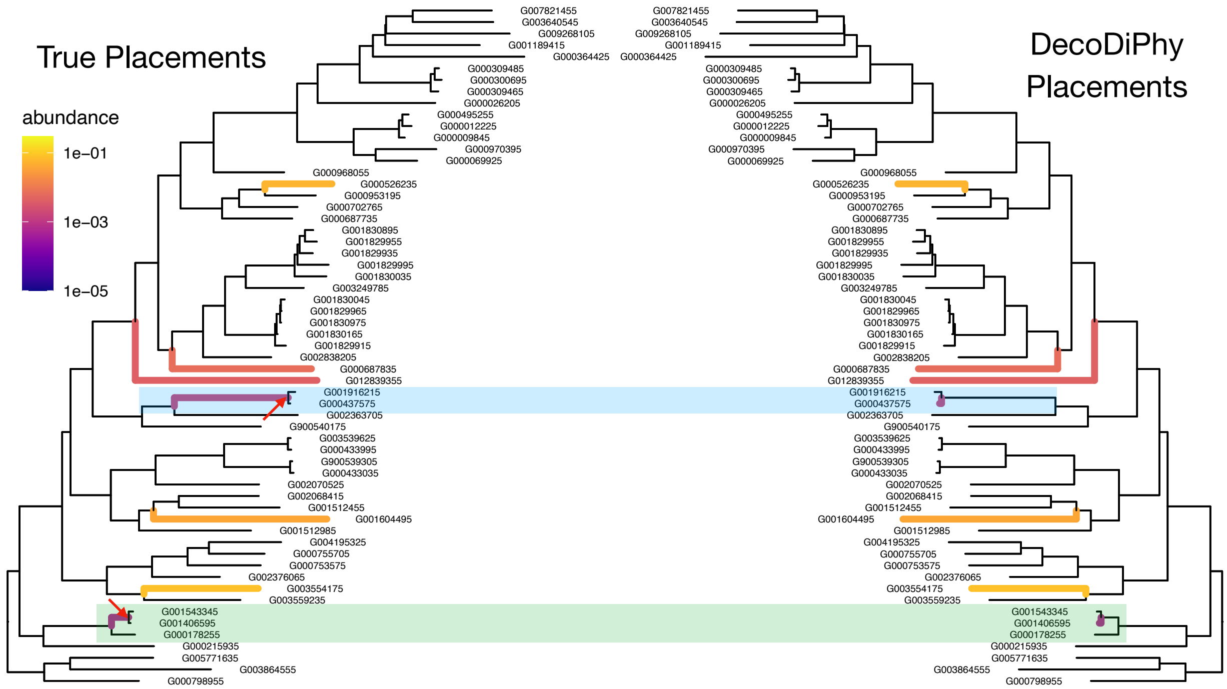

Figure S2: An example of DecoDiPhy failing to identify correct placements for all queries (last column of Table S2). Two out of 28 queries are placed on adjacent branches of the true placements, and the rest are placed on correct branches (we only show the region of the input tree with the two wrong placements). The Jaccard similarity to the true placements is 87%, and UniFrac and weighted UniFrac with respect to true placements are 0.000147 and 0.000154, respectively. Red arrows point to the approximate placements of the queries on the placement edges for these two queries (true distal lengths ($\mathbf{x}$) are 0.002 and 0.0007 for the blue and green placements, respectively), showing that the true placements are phylogenetically close to the estimated placements. Also, note that the estimated abundances of all placements shown in the figure are similar to the true abundances. Also, see Table S2 for all identifiability results.

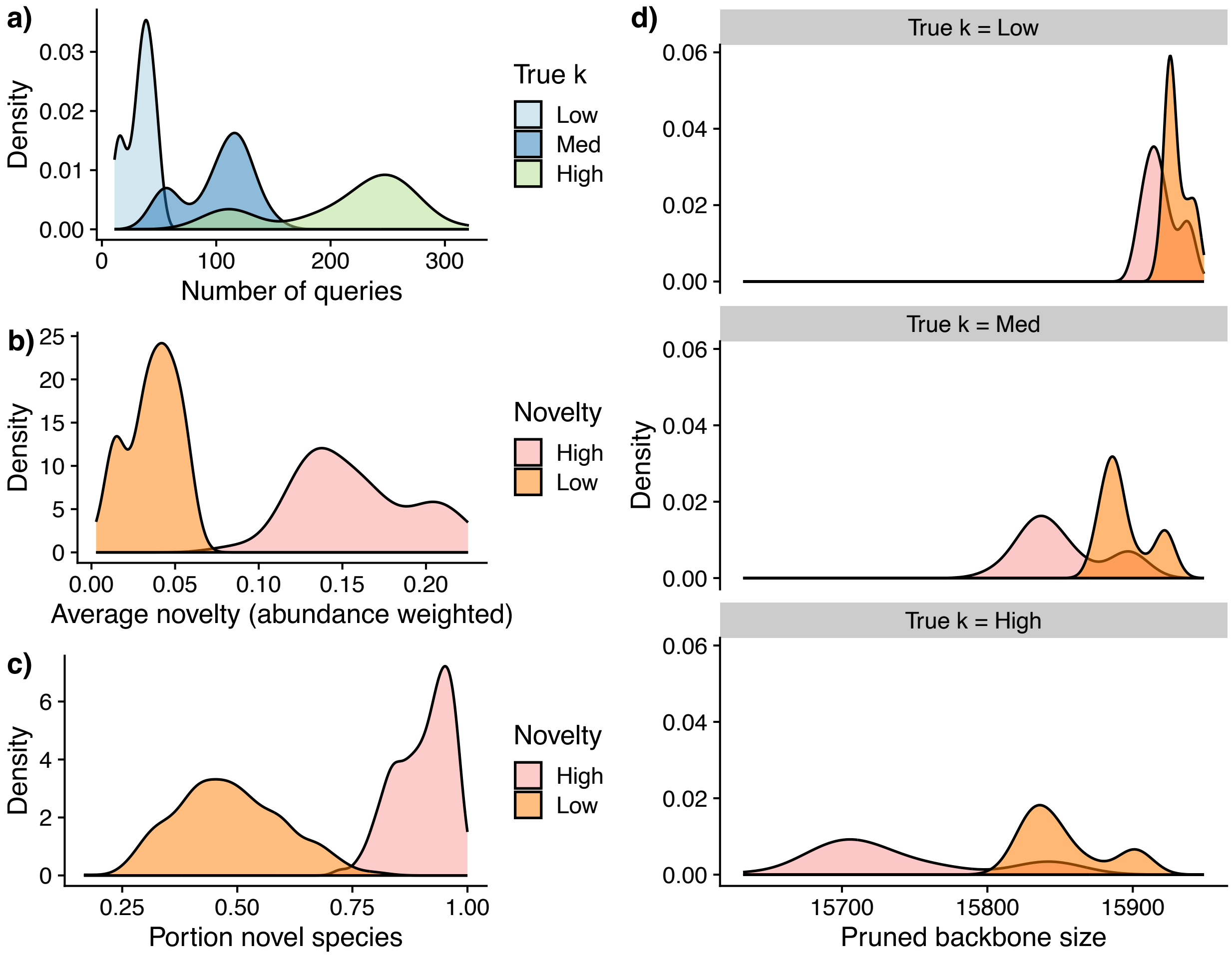

Figure S3: E3. **a)** The number of queries across 50 replicates, for low, med, and high true k and low novelty. The average number of queries for low, med, and high true k is 33, 102, and 214, respectively. **b)** The average novelty across replicates for different novelty levels, quantified by the average distance of the queries to the closest corresponding reference on the pruned tree, weighted by their abundance. **c)** The average portion of the queries without any taxa from the same species present in the backbone tree. **d)** The size of the pruned backbone trees for the two novelty levels across varying numbers of draws.

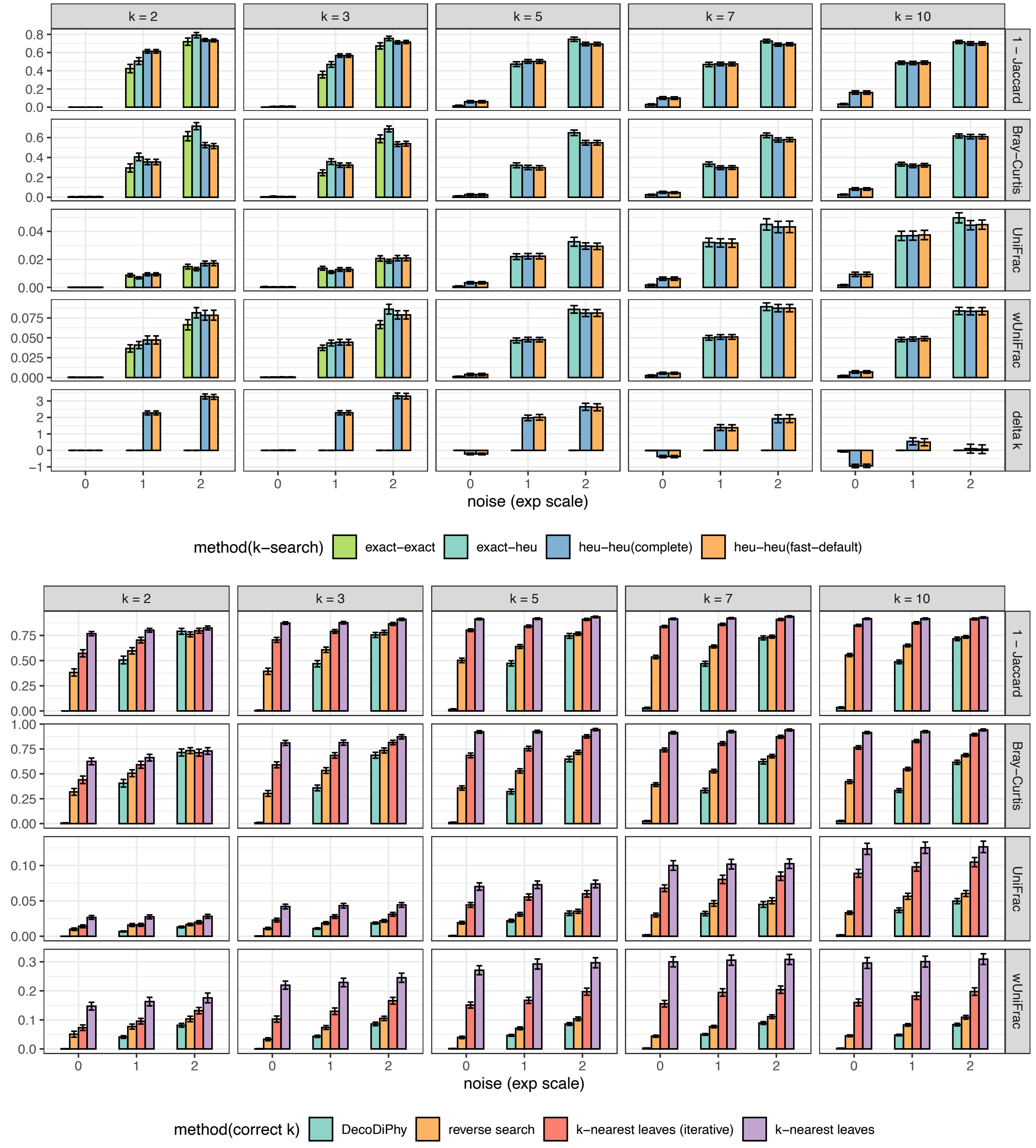

Figure S4: Error on 12 biological trees in the E1 dataset. Bars show the mean and standard error (120 points for noise=0 and 240 points for other noise levels) across four measures of error for three versions of DecoDiPhy and two baseline methods. Note that `exact-exact` variant could be run only for $k \leq 3$. $x-$axis: noise in input distance. Metrics consider different aspects: Jaccard only placement edges, Bray-Curtis only placements and abundance, UniFrac only placements and branch lengths, wUniFrac: placement, branch length, and abundance.

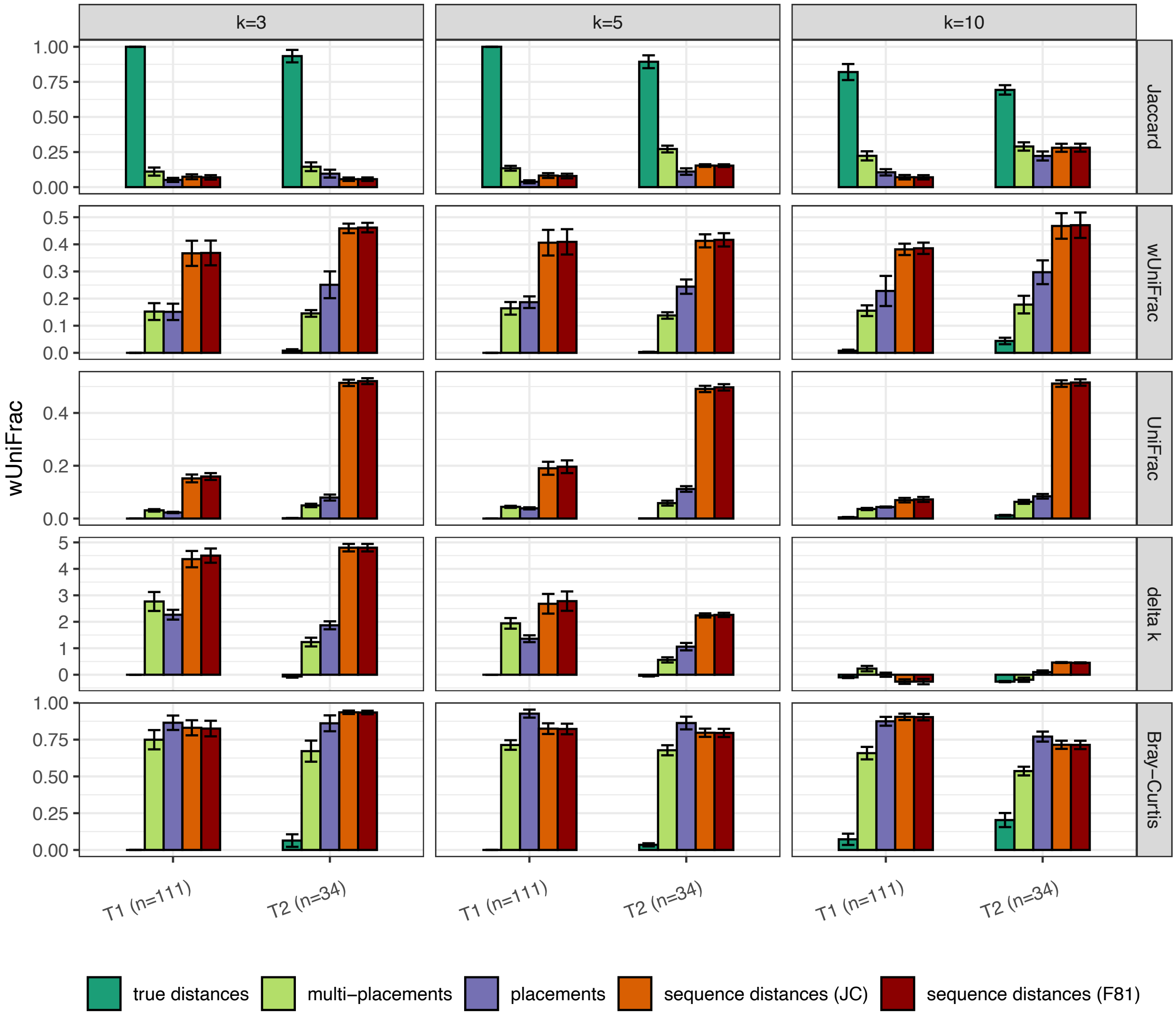

Figure S5: E2. Impact of distance calculation method, comparing true distances, distances from a placement method, or from sequences, on two small subtrees. For sequence-based distances, we show estimates based on two sequence evolution models, Jukes-Cantor (JC) and Felsenstein's 1981 model (F81). See *Methods* for details. Each row corresponds to a different true k. Metrics consider different aspects: Jaccard only placement edges, Bray-Curtis only placements and abundance, UniFrac only placements and branch lengths, and wUniFrac: placement, branch length, and abundance.

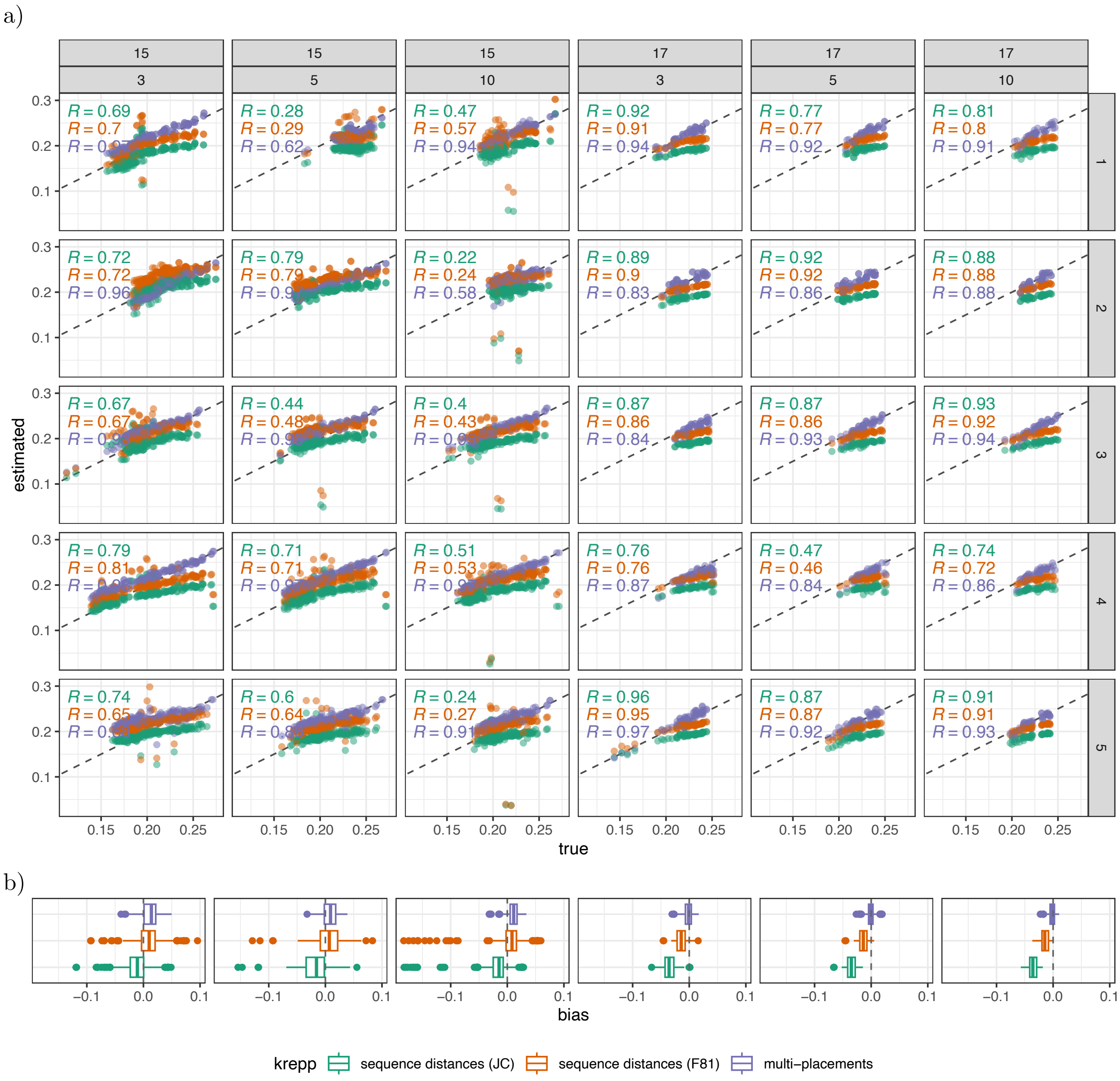

Figure S6: Distance accuracy in E2. **a)** We show the true distance, defined by (3) on the true tree versus estimated distances using krepp sequence distances and krepp multi-placements. For sequence-based distances, we show estimates based on two sequence evolution models, Jukes-Cantor (JC) and Felsenstein's 1981 model (F81). See *Methods* for details. Each row is a replicate. Each dot is one query mixture against one reference species. Pearson correlation coefficients for each method are shown. Dashed black line: the unity line. **b)** We show bias for each estimate defined as estimated − true, averaged across 5 replicates per subtree and true $k$. On average, estimated distances from krepp multi-placements have lower bias, and between the sequence-based methods, Jukes-Cantor has a higher underestimation bias than F81.

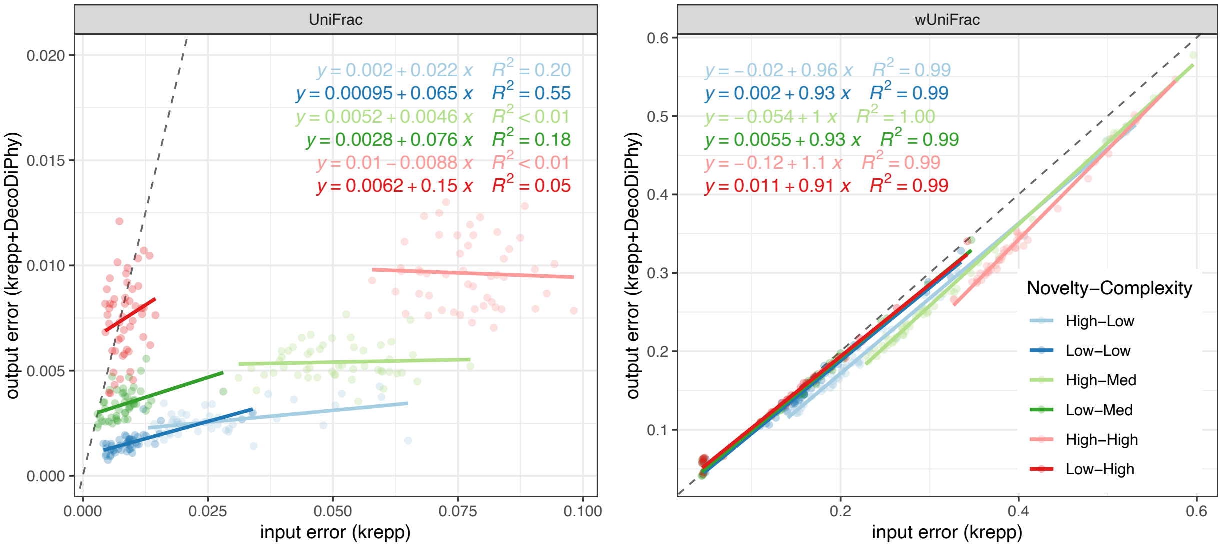

Figure S7: E3. Error of input (krepp) placements versus output (krepp+DecoDiPhy) placements for two levels of novelty (Low/High) and three levels of complexity denoting true $k$are (Low/Med/High). UniFrac (left) and weighted UniFrac (right) are shown. For each condition, a line is fitted to the data, and the slope and the intercept is shown, as well as the Pearson correlation coefficients between input and output errors. The dashed line shows the unity line. According to the UniFrac measure, which is impacted by noisy low-abundance placements, the error in input (krepp) has no to moderate correlation to output error (note low $R$ and slopes), showing that krepp's noise does not propagate directly to DecoDiPhy error (i.e., DecoDiPhy successfully denoises the krepp placements). According to the wUniFrac measure, which is not sensitive to low-abundance noise, the input error translates to output error, but the overall error is reduced (note that most data points are below the unity line). Thus, some forms of input error (not related to low-abundance noise) do carry over from input (krepp) to DecoDiPhy.

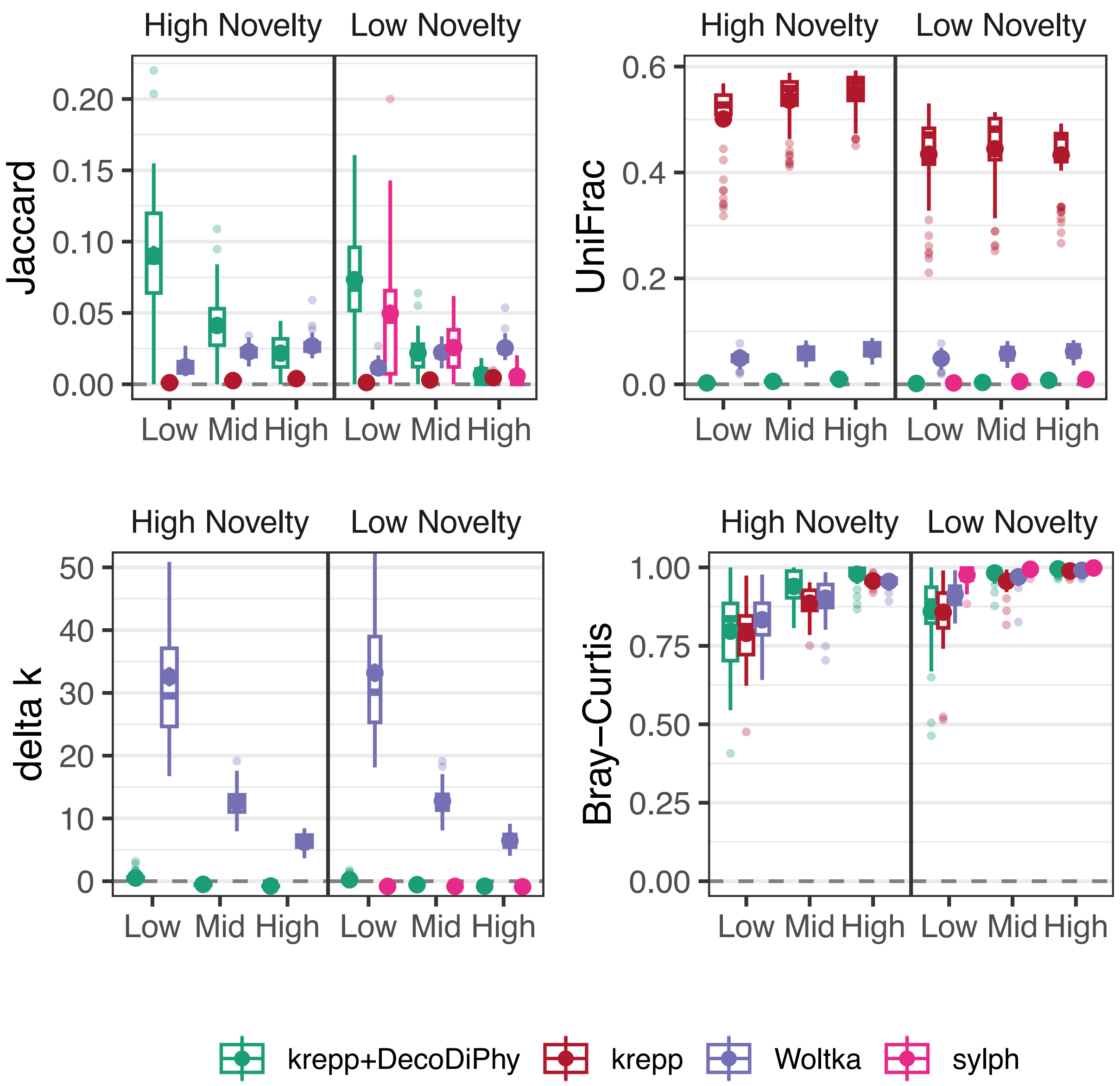

Figure S8: E3. For six model conditions (two novelty levels $\times$ three levels for true $k$), each with 50 replicates, we compare single read placement (krepp), DecoDiPhy run on krepp, single read mapping (Woltka), and joint sample mapping. Average true $k$ for low, med, and high $k$ model conditions is 33, 102, and 214, respectively. On high novelty, sylph found no matches. All placements are considered in the computation of all metrics. For each metric, boxplots as well as mean and standard error are shown for each method. krepp is removed from the delta $k$ figure for a better comparison of DecoDiPhy and sylph since krepp has a much higher error and would dominate the scale. See $\Delta k$ metric in Fig. 2d for a comparison of all methods.

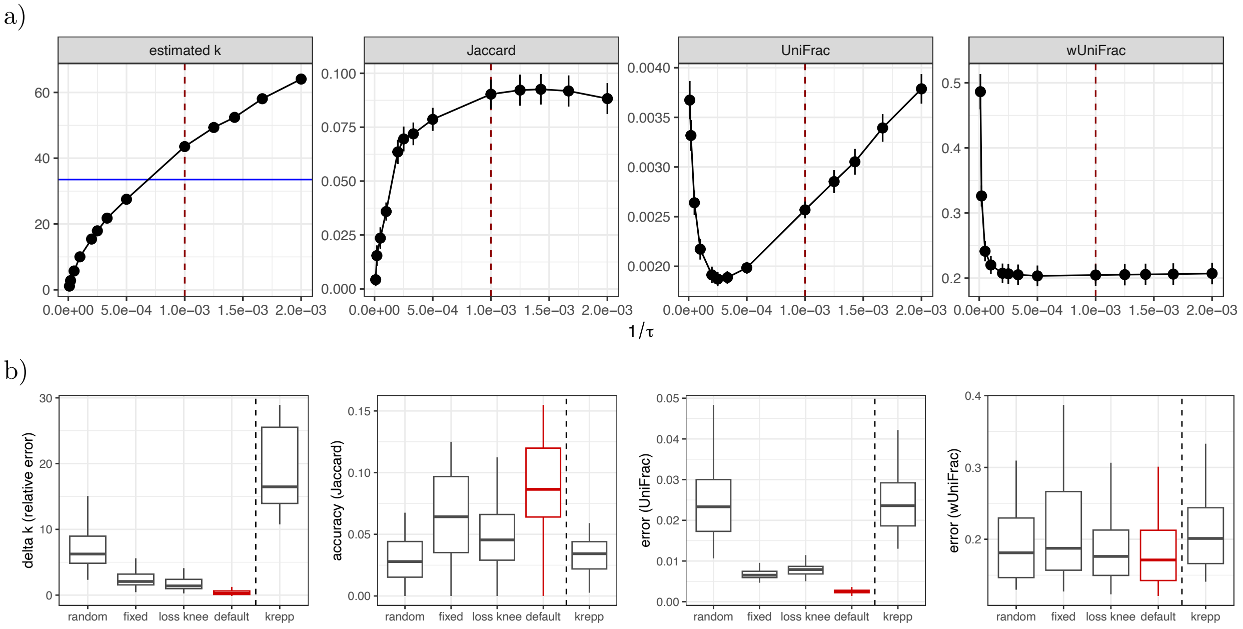

Figure S9: E3. **a)** For the high novelty and low complexity (true $k$) model condition (50 replicates), we ran DecoDiPhy with the default stopping criteria ($\mathbf{p}_q < {}^{\tau}/_{|\mathcal{X}|}$; $|\mathcal{X}|$ is the total number of reads), but different thresholds ($\tau \in \{500, 600, 700, 800, 1000, 2000, 3000, 4000, 5000, 10000, 20000, 50000, 10000\}$); the default is 1000. Lower threshold (higher ${}^{1}/_{\tau}$) results in a higher number of inferred placements. For each threshold, we computed the number of inferred placements (estimated $k$), accuracy (Jaccard), and error (UniFrac and wUniFrac) to the true placements. The average true number of placements is shown with the blue line. The defaults threshold ($\tau = 1000$) is shown with a red dashed line. Note that while the number of placements is a direct function of $\tau$, Jaccard and wUniFrac are to some extent robust to the choice of $\tau$. On the other hand, UniFrac, which ignores abundances, changes substantially with the choice of $\tau$. Our default threshold of 1000 is not selected to be optimal according to any of the metrics; it simply reflects our intuition that detection of a genome with less than 1000 reads ($\approx$ 150Mbp) mapped to it is hard to justify. **b)** On the same dataset, we explored different stopping criteria for DecoDiPhy as well as the default with $\tau = 1000$. For each sample, we ran DecoDiPhy for $max\ k = 500$ rounds. The tested stopping criteria are: random ($k$ is chosen uniformly from $[1, max\ k]$), fixed ($k = 100$), and loss knee (we used an automatic knee locator algorithm called kneed [99] on loss values to find the convergence point). We also show krepp results as a baseline for comparison. While various stopping criteria produce differences in delta $k$, UniFrac, and Jaccard, they all outperform krepp in most cases (even random selection of $k$). Note that wUniFrac is highly robust to the choice of $k$. The default stopping criteria performs the best among all tested criteria and across all metrics.

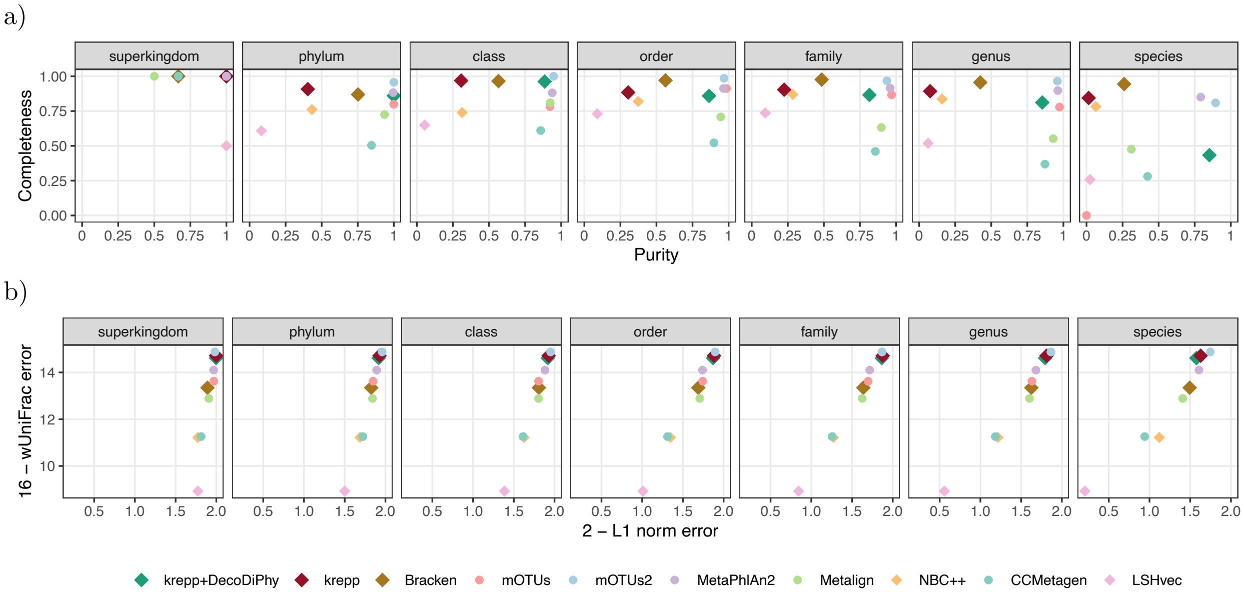

Figure S10: E4 (CAMI-II dataset). Performance of taxonomic profiling on marine dataset across all taxonomic levels. Panels show purity versus completeness (**a**), and 2−L1 norm versus 16−weighted UniFrac (**b**). Note that L1 norm and weighted UniFrac are by design bounded to $[0, 2]$ and $[0, 16]$, respectively. Closer to top right corner is better. $k$-mer-based methods as well as krepp+DecoDiPhy are shown with a diamond shape, while the alignment-based methods are shown with circles.

a)

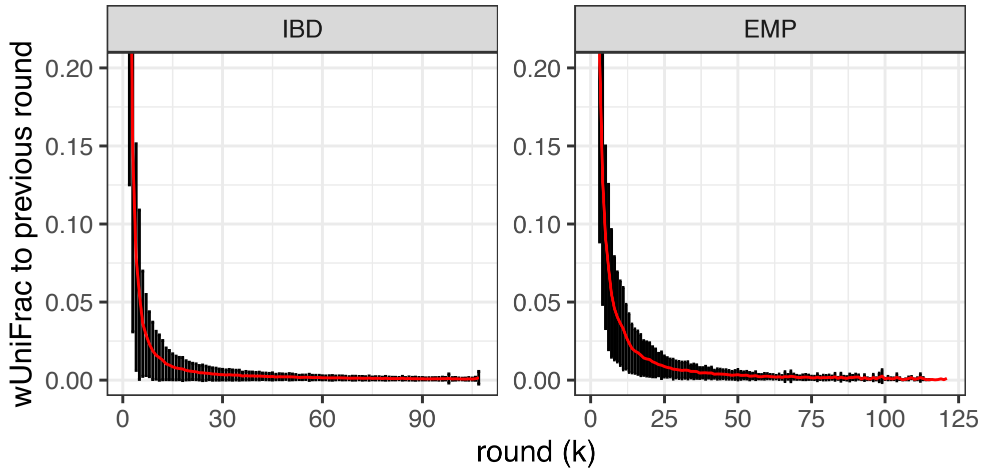

b)

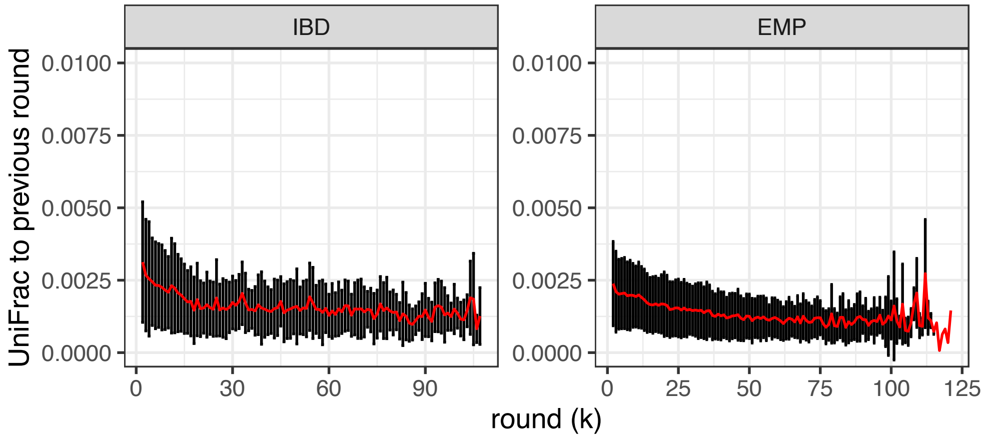

Figure S11: IBD and EMP datasets. The distribution of wUniFrac **(a)** and UniFrac **(b)** to the previous round $(k-1)$ across all samples. The mean (red) and standard deviation (black error bars) across $220 \times 16$ (subtree per sample) replicates are shown for each $k$. wUniFrac shows convergence as $k$ increases, indicated by a close to zero distance to the previous round after roughly $k > 50$. Thus, new queries are either close to the previous ones or have low abundance, or both. UniFrac values do not converge to zero, likely due to the fact that UniFrac does not take into account abundances, and therefore, the addition of any new placements to the previous round impacts UniFrac more than wUniFrac. Nevertheless, the downward trends show that new queries tend to be placed phylogenetically close to the previous ones.

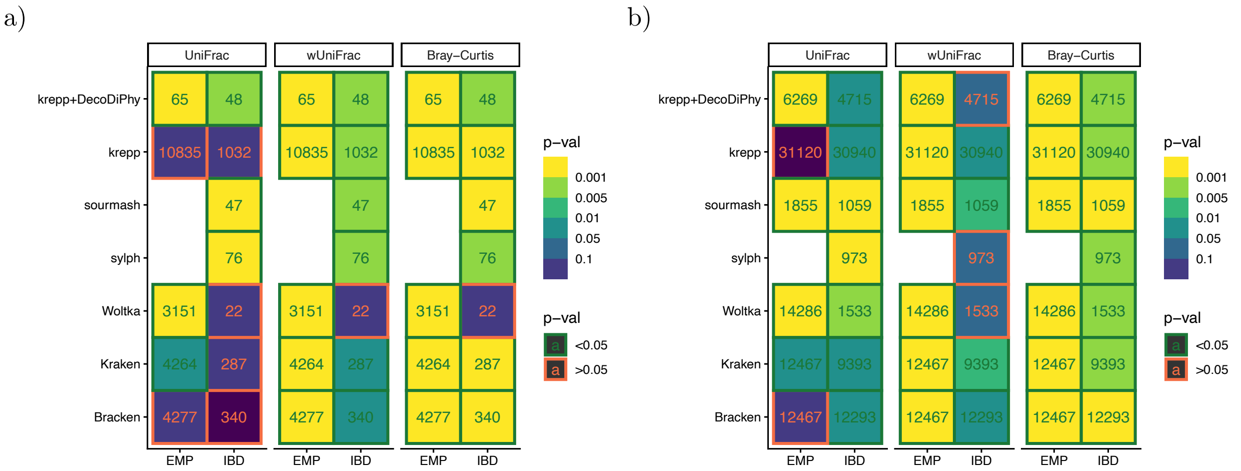

Figure S12: IBD and EMP datasets. The significance of separation between groups for **a)** DA and **b)** all features. Significant p-values (p-value $< 0.05$) are marked in green, and insignificant p-values (p-value $> 0.05$) are marked in red. The total number of features for each method is written in text. sylph was removed for the EMP dataset (both all and DA features), because it failed to report any placements for 79 out of 210 samples. Similarly, only 72 out of 210 samples have any of the 9 DA features found for sourmash on the EMP dataset; therefore, we only report the significance of separation for all features for this method.

a)

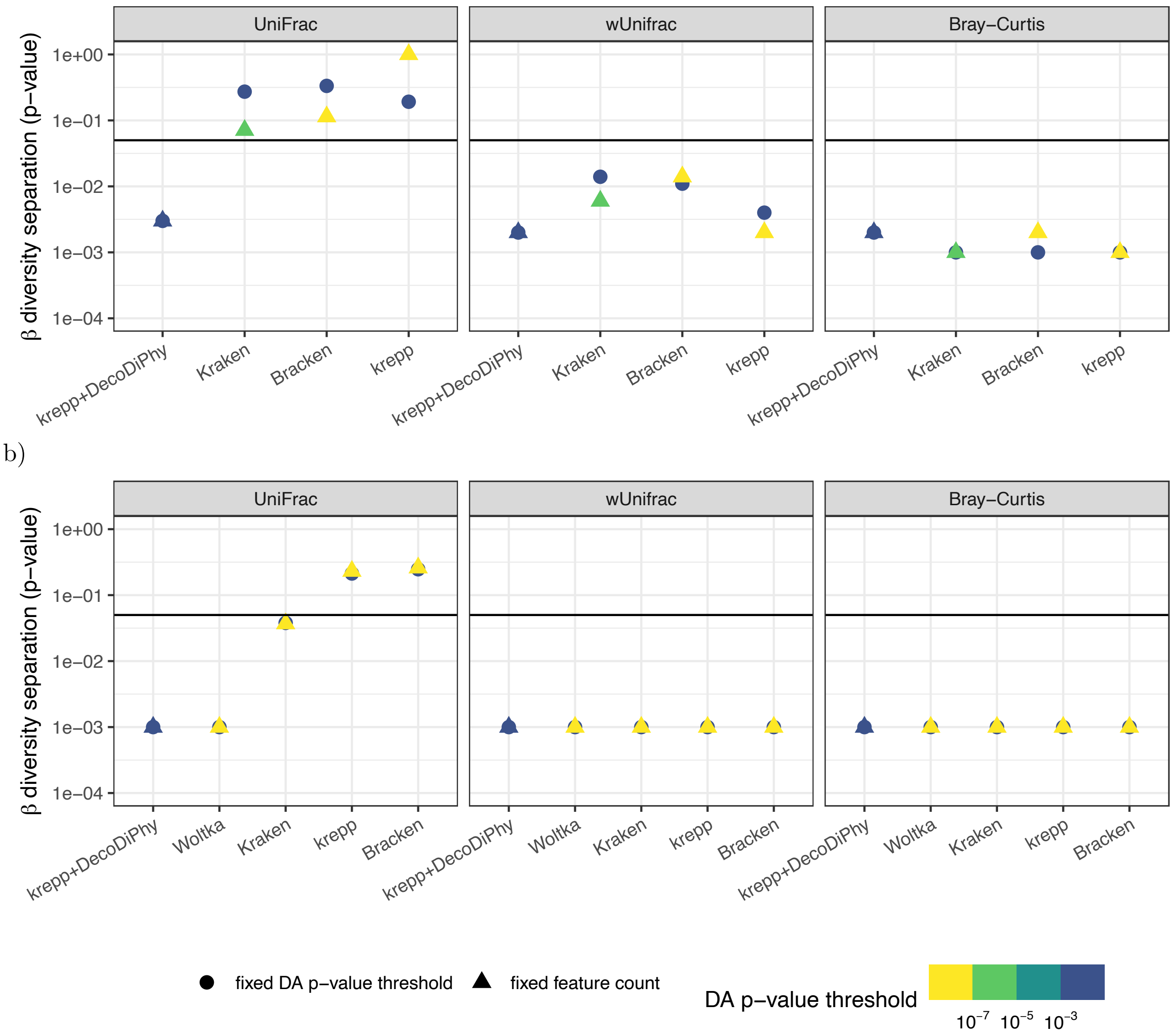

Figure S13: IBD and EMP datasets. The change in the significance of separation between groups for a fixed ANCOM-BC p-value threshold versus fixed number of DA features for IBD (**a**) and EMP (**b**) datasets. For the fixed number of features, the same number of DA features as DecoDiPhy were selected for each method (48 for IBD and 65 for EMP dataset). Only methods with more than 100 original DA features are shown. Colors show the ANCOM-BC p-value threshold that would have been used to achieve the selected number of features for each method. The $\alpha = 0.05$ significance level (for diversity separation) is marked with black line.

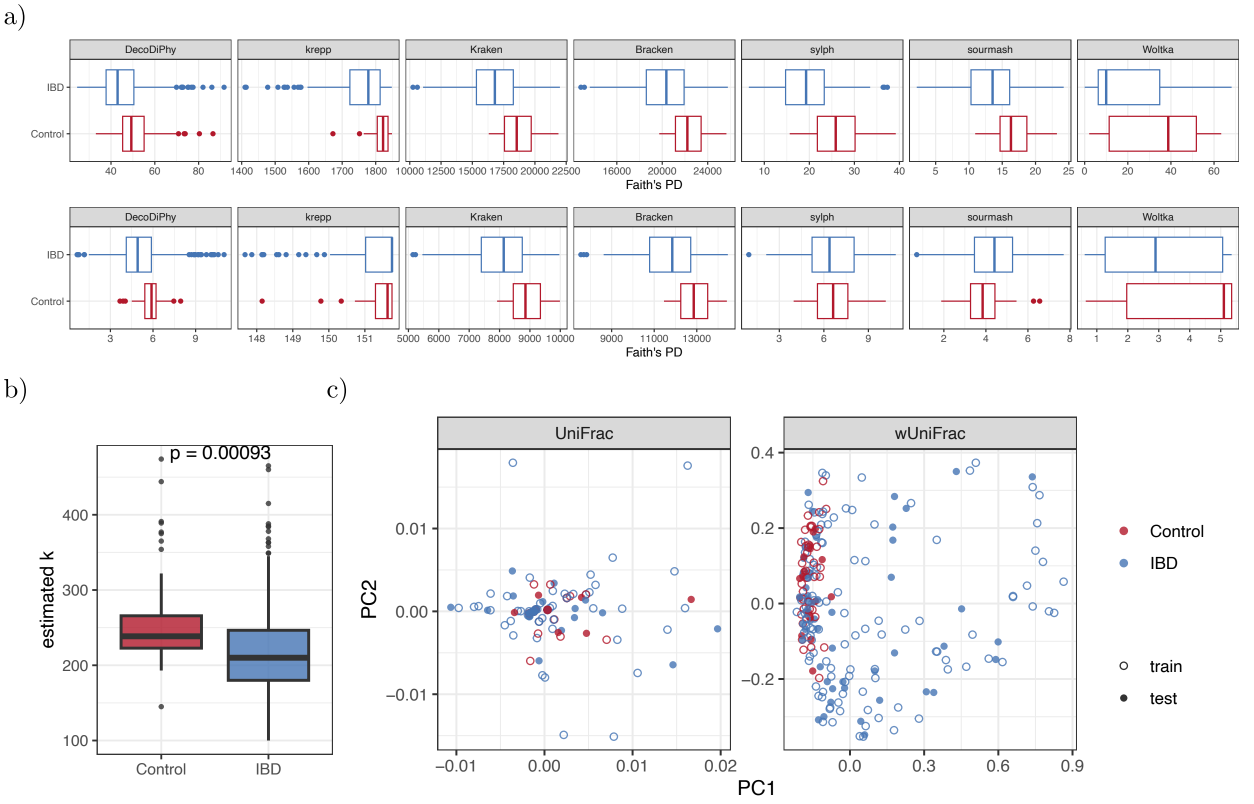

Figure S14: IBD dataset. **a)** Distribution of Faith's phylogenetic diversity for all (top) and DA (bottom) features for IBD versus control samples. **b)** The distribution of the estimated number of placements by DecoDiPhy (estimated $k$) for each group. The p-value for the Welch two-sample t-test is shown, testing if the estimated number of placements differs significantly between groups. **c)** Principle coordinates of UniFrac and wUniFrac on DA features for krepp, showing separation between IBD and control samples. Due to the high number of DA features, krepp fails to separate samples with UniFrac.

a)

b)

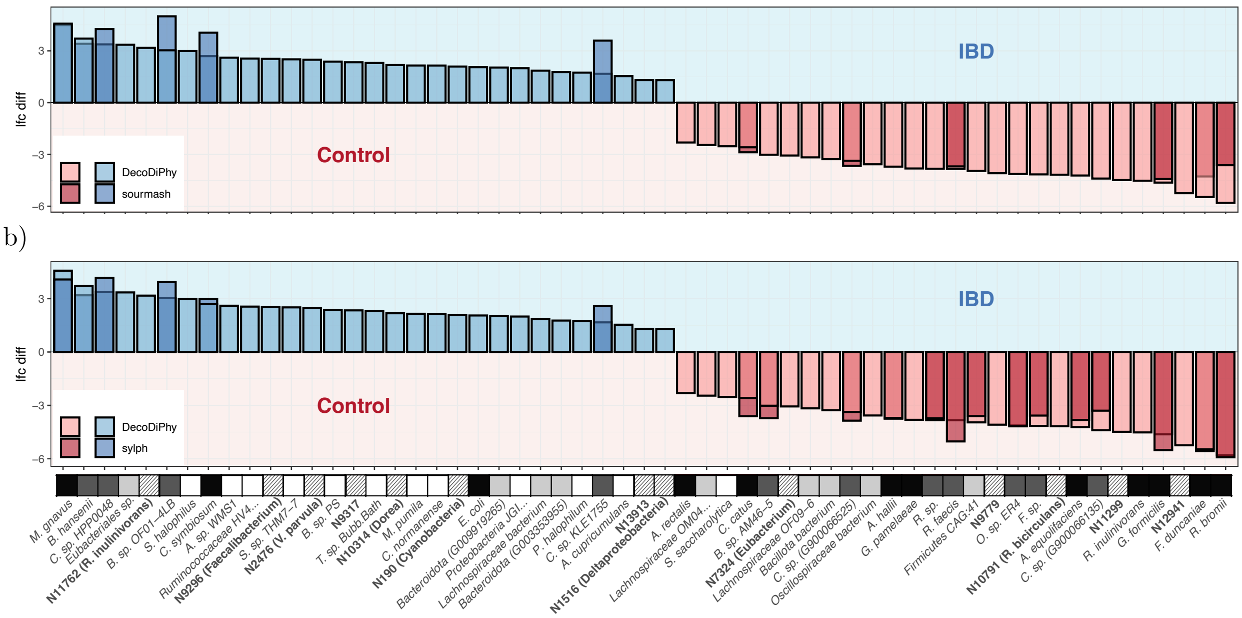

Figure S15: IBD dataset. DA features identified by DecoDiPhy on IBD data, together with their log-fold change (LFC) in abundance. sourmash **(a)** and sylph **(b)** LFCs shown for features that are also in their DA feature set. Colors associated with each feature show the level of prior association with IBD in the literature. See Fig. 4c for a description of each category. Many DA features identified by DecoDiPhy that have known prior associations with IBD on species or genus level, were also identified by the exact $k$-mer matching methods sylph and/or sourmash.

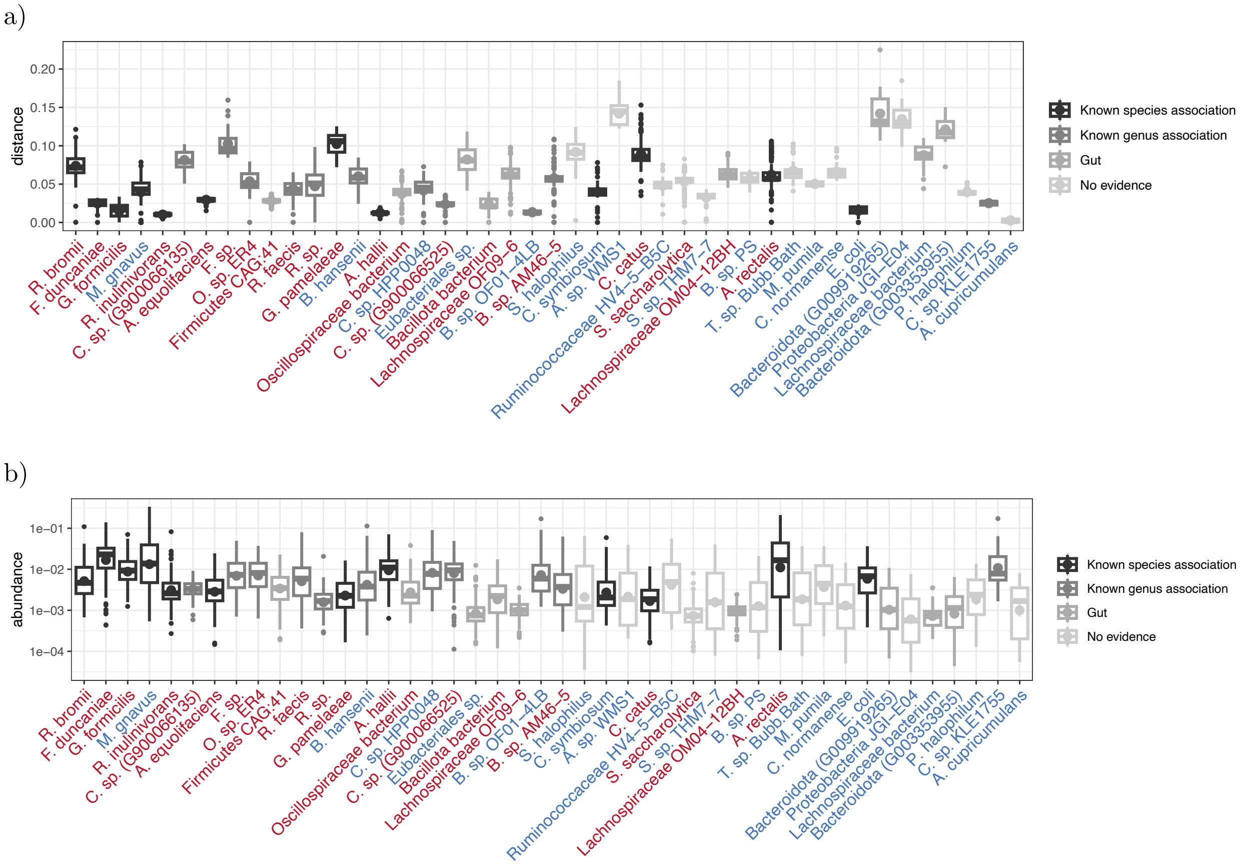

Figure S16: IBD dataset. Distribution of **(a)** tree distance of DecoDiPhy placement to the corresponding reference taxa defined as $x+y$, and **(b)** assigned abundance ($p$) by DecoDiPhy, for terminal DA features identified by DecoDiPhy. DA features are ordered by decreasing absolute log-fold change ($|\text{lfc}|$). Species with low literature support tend to have higher distances to closest taxa, except for *A. cupricumulans*, which has been implicated at least in one study on mice [100]. Similarly, these features tend to have lower abundances than features with stronger literature support.

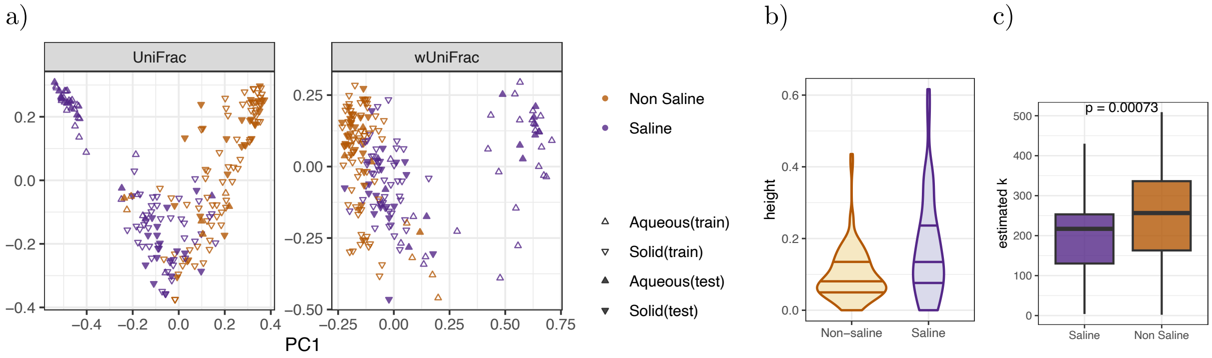

Figure S17: EMP dataset. **a)** Principle coordinates of UniFrac and wUniFrac for DecoDiPhy, showing separation between saline and non-saline free-living samples; In both metrics, PC1 mainly separates solid versus aqueous, though DA features were selected for saline/non-saline, not this categorization. **b)** Distribution of the height of the DA features associated with saline/non-saline. The height of a node $n$ is defined as the average distance of all tips below $n$ to $n$. **c)** The distribution of the number of DecoDiPhy placements ($k$) for each group across samples. P-values for the Welch two-sample t-test are shown, showing that the estimated number of placements differs significantly between groups.

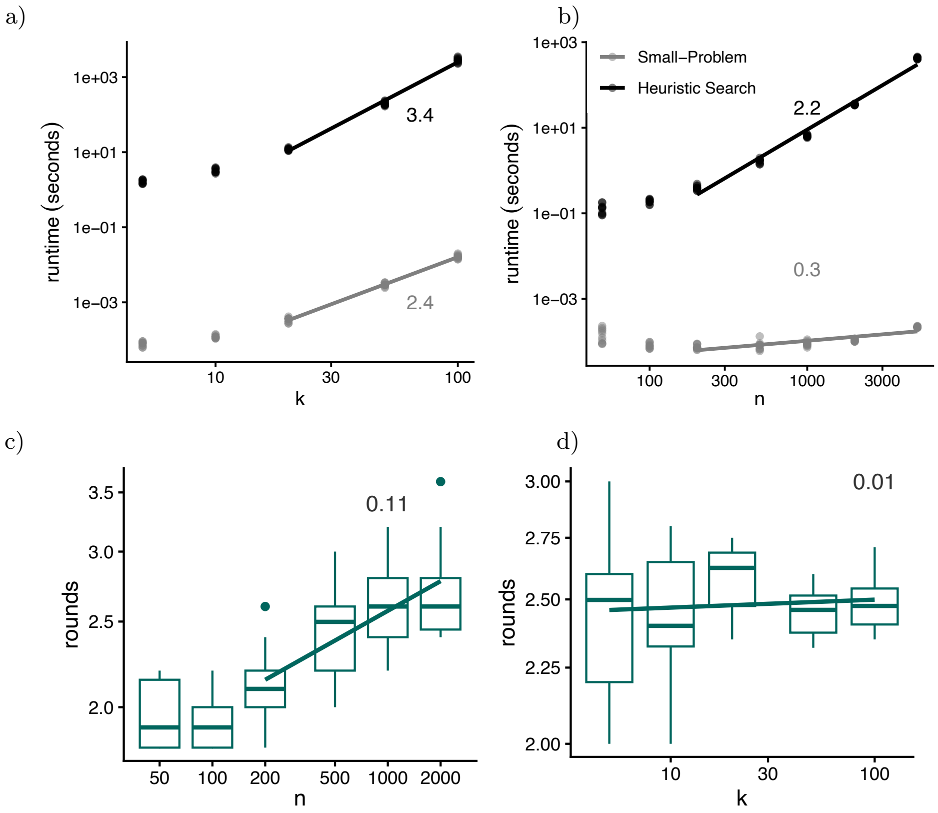

Figure S18: **a,b)** Empirical running time of small-problem and the heuristic search for **(a)** varying $k$ and **(b)** varying $n$, showing log-log plots; line slopes estimate the asymptotic power (excluding low $n, k$ values). For **(a)**, $n = 500$, and for **(b)**, $k = 5$ was used. **c,d)** In experiments described in Runtime analysis, we measure the number of rounds of optimization (Line 18) per each value of $k$ versus $n$ **(c)** or $k$ **(d)**. Both plots are on log-log scales, with the slope of the fitted line showing the power of asymptotic running time. For each replicate, the DecoDiPhy method is run with true $k$. Each dot is the total number of rounds for a replicate divided by $k$. Note that the number of rounds grows very slowly with $n$ and $k$, on average 2.3 across all values of $n$ and ever more than 3.6, motivating us to assume $C$ is a constant. For **(c)**, $k = 5$, and for **(d)**, $n = 500$ was used.

### B Supplementary tables

Table S1: Current microbiome methods do not jointly analyze reads or do not fully consider the phylogeny.

| Analysis Type | Problem Statement | Dependence* Reads | Dependence* Evolution | Example Software |
|---|---|---|---|---|
| Closed reference | Assign each read to references, grouped into operational taxonomic/genomic units (e.g., OTUs or OGU) | No | No | Woltka [12], DADA [11], krepp [22] |
| Read classification | Assign to each read a taxonomic label at the most precise rank possible | No | Limited | Kraken2 [31], Centrifuge [101], CONSULT-II [32], Ganon [33] |
| Per-read placement | Place each read on a phylogenetic tree | No | Full | pplacer [18], EPA [19], SEPP [21] DEPP [102], krepp [22], App-SpaM [20] |
| Profiling | Consolidate reads ids into sparser profiles | Limited | Limited | Bracken [37] |
| Joint taxon profiling | Deconvolve a mixed sample by assigning it jointly to independent units (e.g., species) | Yes | Limited/No | MetaPal. [38], sourmash [39], Mash Screen [40], sylph [41] |
| PDD | Deconvolve a mixed sample into a multi-placement, accounting for the tree metric | Yes | Full | DecoDiPhy (this paper) |

* Approaches that bin into independent buckets ignore evolutionary dependence; we consider taxonomic hierarchies as "limited" modeling of those dependencies. Tools that analyze reads one by one ignore read dependencies.

Table S2: Results of the identifiability analysis to test the limitations of the PDD problem and the heuristics used to achieve scalable running time.

(a) Number of replicates with exact solutions (average Jaccard similarity to ground truth for each case is also shown in percentage):

| | | | loss $> \epsilon$ | loss $\approx 0$ |
|---|---|---|---|---|
| $k$ | Total # replicates | Correctly Recovered† (Jaccard) | Sub-Optimal Solution‡ | Unidentifiable within $\epsilon^*$ |
| $\log_2(n)$ | 160 | 160 (100%) | 0 | 0 |
| $\sqrt{n}/2$ | 160 | 157 (100%) | 3 (88%) | 0 |
| $\sqrt{n}$ | 160 | 137 (100%) | 13 (91%) | 10 (93%) |

(b) UniFrac:

| | | loss $> \epsilon$ | loss $\approx 0$ |
|---|---|---|---|
| $k$ | Correctly Recovered | Sub-Optimal Solution | Unidentifiable within $\epsilon$ |
| $\log_2(n)$ | 0.00000196 | - | - |
| $\sqrt{n}/2$ | 0.0000112 | 0.000515 | - |
| $\sqrt{n}$ | 0.0000460 | 0.000579 | 0.000245 |

(c) weighted UniFrac:

| | | loss $> \epsilon$ | loss $\approx 0$ |
|---|---|---|---|
| $k$ | Correctly Recovered | Sub-Optimal Solution | Unidentifiable within $\epsilon$ |
| $\log_2(n)$ | 0.0000835 | - | - |
| $\sqrt{n}/2$ | 0.000166 | 0.00113 | - |
| $\sqrt{n}$ | 0.000295 | 0.00185 | 0.00123 |

**Experiment setup:** We used the subtrees from the WoL backbone tree as described in Section C.7.4. For each subtree, we resolved polytomies randomly and increased the length of super short branches (less than $1e-3$) by a positive number ($\sim Normal(0, 1e-3)$). We then created 10 mixtures by randomly selecting $k \in \{\log_2(n), \sqrt{n}/2, \sqrt{n}\}$, where $n$ is the size of the input tree ($n = 997$ on average), resulting in 160 total replicates per each choice of $k$. We used the true distance vector as input and ran the default version of DecoDiPhy on each sample. We divide replicates based on the accuracy of the placements and final loss residuals.

† Cases where all placement edges are correctly recovered (Jaccard = 100%) are considered as exact solutions.

‡ Cases with $loss > \epsilon$ are considered sub-optimal solutions, a scenario that may occur due to the limitations of our search heuristics.

$*$ All other cases are marked as non-identifiable; however, this inability to recover correct placements may be due to the optimizer's precision limits, and thus, should be interpreted as identifiability within those precision limits.

With $\epsilon = 1e-8$, only 16 and 10 out of 480 replicates resulted in sub-optimal and unidentifiable solutions, respectively; however, even for those scenarios, placement error (UniFrac and wUniFrac) remained extremely low. See Fig. S2 for an example of one of the unidentifiable cases.

Table S3: The biological trees used in simulation analyses

| Group | Publication | $n$ | Availability |
|---|---|---|---|
| bees | https://doi.org/10.1093/sysbio/syaa097 | 30 | CASTLES-pro branch length on ASTRAL tree from https://doi.org/10.5061/dryad.z08kprrb6 available on https://github.com/ytabatabaee/CASTLES-Pro-paper/tree/main/data/biological/bees |
| mammals (Song) | https://doi.org/10.1073/pnas.1211733109 | 36 | CASTLES-pro branch length available from https://github.com/ytabatabaee/CASTLES-Pro-paper/tree/main/data/biological/mammals |
| birds (Jarvis) | https://doi.org/10.1126/science.1253451 | 48 | The TENT.RAxML-GAMMA tree form Newick_tree_files.tar.gz of http://gigadb.org/dataset/101041 |
| tilapia | https://doi.org/10.1093/molbev/msae116 | 91 | Mzeb_nc_iqtree.min4.phy.treefile from https://doi.org/10.5061/dryad.p2ngf1w0c |
| plants (1kp pilot) | https://doi.org/10.1073/pnas.1323926111 | 103 | FNA2AA.trim50genes50sites.no3rd.partitioned.gamma.final.tre tree form https://datacommons.cyverse.org/browse/iplant/home/shared/onekp_pilot/PNAS_alignments_trees/species_level/trees |
| Pancrustacea | https://doi.org/10.1093/molbev/msad175 | 105 | Fig2A_Dataset2_C60LG.tre from https://doi.org/10.5061/dryad.dr7sqvb2h |
| beetles | https://doi.org/10.1016/j.ympev.2018.05.028 | 127 | exabayes_tree_with_bootstraps_rerooted.newick from https://datadryad.org/stash/dataset/doi:10.5061/dryad.5r6d5kc |
| fish (marine) | https://doi.org/10.1073/pnas.2122486119 | 152 | SI_Dataset_S6_IQTREE_Best_partition_scheme.part.contree.tre from https://datadryad.org/stash/dataset/doi:10.5061/dryad.z34tmpgfw |
| hemipteroid insects | https://doi.org/10.1073/pnas.1815820115 | 193 | Supplementary_Archive_4_phylogenies_bootstraps/ML_tree_inference_nucleotide_12/bestTreeWithSupport.newick from https://doi.org/10.5061/dryad.t4f4g85 |
| mammals (Foley) | https://doi.org/10.1126/science.abl8189 | 241 | Concatenation_HRA_neutral_241_10miss_rooted.nexus from https://zenodo.org/records/7958013 |
| fish | https://doi.org/10.1073/pnas.1719358115 | 305 | 1105_protein_ExaBayes.tre from https://datadryad.org/stash/dataset/doi:10.5061/dryad.5b85783 |
| birds (Stiller) | https://doi.org/10.1038/s41586-024-07323-1 | 363 | CASTLE-Pro on the original ASTRAL tree, available from https://github.com/ytabatabaee/CASTLES-Pro-paper/tree/main/data/biological/birds-stiller |

Table S4: IBD dataset. DecoDiPhy DA features. For each feature, taxonomic label, log-fold change in IBD abundance (LFC) compared to control, and association to IBD in prior studies is shown.

| Genome ID | Taxonomic Label | LFC | Association with IBD | other methods |
|---|---|---|---|---|
| G004554925 | *Ruminococcus bromii* | -5.8 | *Ruminococcus bromii* is a keystone species for the degradation of resistant starch in the human colon [103]. [55, 104–107] report more abundance in the healthy subjects than in the CD patients. | sourmash, sylph |
| G000162015 | *Faecalibacterium duncaniae* | -5.5 | Many studies have reported higher abundance of *F. duncaniae* in healthy samples compared to IBD samples [108, 109]. | sourmash, sylph |
| N12941 | *Ruminococcus obeum* (*Blautia obeum*) | -5.2 | Several studeis have reported enrichments of *Blautia obeum* in non-IBD controls [48, 58, 106, 110] | |
| G902363055 | *Gemmiger formicilis* | -4.6 | Reports of depletion in the gut microbiota of IBD samples [55, 59, 105, 106]. | sourmash, sylph |
| G000174195 | *Roseburia inulinivorans DSM 16841* | -4.5 | [57, 58, 106, 111] report significantly lower abundance in IBD patients as compared to healthy individuals. | |
| N11299 | parent of *Dorea longicatena* and *Dorea sp. 42_8* | -4.5 | *Dorea longicatena* is reported to be significantly reduced in the IBD group [57, 58, 107]. | |
| G900066135 | *uncultured Clostridium sp.* | -4.4 | Decreased abundance of *Clostridium* genus has been reported in IBD [55, 57, 62]. | sylph, sourmash |
| G000478885 | *Adlercreutzia equolifaciens DSM 19450* | -4.2 | [112, 113] reduction in *Adlercreutzia equolifaciens* in IBD cohort. [114] reports an increased abundance in *Adlercreutzia equolifaciens* after recieving composite probiotics. | sylph |
| N10791 | *Ruminococcus bicirculans* | -4.2 | [58, 107] show depletion of *Ruminococcus bicirculans* in IBD vs. healthy samples. | |
| G900542435 | *uncultured Faecalibacterium sp.* | -4.2 | *Faecalibacterium* genus is consistetly reported to have lower abundaces in IBD samples [108, 109]. | sylph |
| G000765235 | *Oscillibacter sp. ER4* | -4.1 | *Oscillibacter* genus is consistetly reported to have lower abundaces in IBD samples [68, 115, 116]. | sylph |
| N9779 | parent of *Firmicutes bacterium CAG:83* and *uncultured Eubacteriales bacterium* | -4.1 | *Firmicutes bacterium CAG:83* is reported to be depleted in IBD [106]. [55, 68] report lower abundance of Firmicutes phylum in IBD samples. | |
| G000431615 | *Firmicutes bacterium CAG:41* | -4.0 | [55, 68] report lower abundance of Firmicutes (Bacillota) phylum in IBD samples. [117] reports differentially higher abundances of *Firmicutes bacterium CAG:41* in healthy mice compared to depressed mice. | sylph |
| G001312505 | *[Ruminococcus] faecis JCM 15917* | -3.8 | The genera *Ruminococcus* in general in reported to be more abundant in healthy samples [59]. | sourmash, sylph |
| G900066095 | *uncultured Ruminococcus sp.* | -3.8 | The genera *Ruminococcus* in general in reported to be more abundant in healthy samples [59]. | sylph |
| G000210055 | *Gordonibacter pamelaeae 7-10-1-b* | -3.8 | [48, 106] report decrease of *Gordonibacter pamelaeae 7-10-1-b* in IBD cohort. | |
| G000173975 | *Anaerobutyricum hallii* (formerly *Eubacterium hallii*) DSM 3353 | -3.7 | [55, 57, 106, 118, 119] reported a significantly lower abundance of *E. hallii* in IBD patients compared to healthy individuals. *E. hallii* may be beneficial for IBD patients and could serve as potential probiotics for IBD treatment [120]. | sylph |

| Genome ID | Taxonomic Label | LFC | Association with IBD | other methods |
| --- | --- | --- | --- | --- |
| G902363685 | *Oscillospiraceae bacterium* | -3.6 | [54, 121] reports lower abundance in both CD and UC for *Oscillospiraceae* family. | |
| G900066525 | *uncultured Clostridium sp.* | -3.4 | The *Clostridium* genus, found in greater abundance in healthy patients [59]. | sourmash, sylph |
| G902387315 | *Bacillota bacterium* | -3.3 | [55] reports lower abundance of *Bacillota* phylum in IBD. Some key *Bacillota* species are fiber-degrading and produce SCFAs, thus generally exhibiting anti-inflammatory effects in the gut [122]. | |
| G003435055 | *Lachnospiraceae bacterium OF09-6* | -3.2 | The *Lachnospiraceae* family has been reported to be depleted in IBD [52]. | |
| N7324 | parent of *Eubacterium sp. CAG:603* and *uncultured Eubacterium sp.* | -3.1 | [111] reports lower abundance of *Eubacterium* genus in IBD. | |
| G003479885 | *Blautia sp. AM46-5* | -3.0 | Evidence on depletion of *Blautia* genus in IBD [52, 60, 123]. | sylph |
| G000210555 | *Coprococcus catus GD/7* | -2.6 | *Coprococcus catus* was reported to have lower abundance in IBD [58, 59, 106]. | sourmash, sylph |
| G005048345 | *Soehngenia saccharolytica* | -2.5 | No prior relatios to IBD or human gut. | |
| G003490145 | *Lachnospiraceae bacterium OM04-12BH* | -2.5 | The family *Lachnospiraceae* is reported to have lower abundance in IBD samples [59]. | |
| G000020605 | *Agathobacter rectalis ATCC 33656* | -2.3 | *Agathobacter rectalis* was found as the shared markers of CD and UC that are depleted in active IBD [55, 104–106, 124]. | |
| N1516 | *Deltaproteobacteria* class | 1.3 | Many *Deltaproteobacteria* in the gut are sulfate-reducing bacteria (SRB), such as *Desulfovibrio* and *Desulfomicrobium* [125], which have been reported to be associated with IBD [66, 67]. | |
| N13913 | *Lactobacillus* clade (including *Lactobacillus sp. HMSC24D01*) | 1.3 | Research on *Lactobacillus* is mixed. While some can be beneficial [126], some studies have reported over-abundance of some *Lactobacillus* species in IBD samples [55, 57, 58, 64]. | |
| G001402935 | *Acidiplasma cupricumulans* | 1.5 | No evidence on presence in human gut or relations to IBD. | |
| G000466465 | *Clostridium sp. KLE 1755* | 1.7 | Reports of increase of *Clostridium* species in IBD [55, 127]. | sourmash, sylph |
| G002245355 | *Paludifilum halophilum* | 1.7 | No link to human gut or IBD. | |
| G003353955 | *Bacteroidota bacterium* | 1.8 | [57] report an increase in abundance of another species from this phylum, *Bacteroides fragilis*, in IBD patients. | |
| G011960265 | *Lachnospiraceae bacterium* | 1.9 | [54] report higher abudance of some *Lachnospiraceae* species in IBD. | |
| G000428205 | *Proteobacteria bacterium JGI 0000113-E04* | 2.0 | [55] reports increase in *Proteobacteria* phylum in IBD samples. | |
| G009919265 | *Bacteroidota bacterium* | 2.0 | [128] reports a increase in *Bacteroidota* Phylum in IBD patients. | |
| G000944855 | *Escherichia coli* | 2.1 | Many studies have reported enrichment of *E. coli* in IBD samples [48, 55, 57, 104–107, 129–131]. | |

| Genome ID | Taxonomic Label | LFC | Association with IBD | other methods |
|---|---|---|---|---|
| N190 | *Cyanobacteria* Phylum | 2.1 | [57, 68] observed higher abundances of *Cyanobacteria* phylum in IBD samples compared to healthy samples. | |
| G001747495 | *Cloacibacterium normanense* | 2.1 | No link to IBD or human gut. | |
| G003596335 | *Motilimonas pumila* | 2.1 | No link to IBD or human gut. | |
| N10314 | *Dorea* Genus | 2.2 | [110, 123] found positive associations between *Dorea* genus and IBD. | |
| G012027595 | *Thermococcus sp. Bubb.Bath* | 2.3 | An archaea with no link to IBD or human gut. | |
| N9317 | parent of *Drancourtella massiliensis* and *Sellimonas intestinalis* | 2.3 | [132] identified *Sellimonas* genus as characteristic of IBD groups. [133] reported decrease in *Drancourtella massiliensis* in reaction to probiotic intervention. | |
| G000170715 | *Beggiatoa sp. PS* | 2.4 | No evidence on IBD or human gut. | |
| N2476 | *Veillonella parvula* | 2.5 | Reports of increase in IBD samples [55, 58]. | |
| G004153225 | *Sporolactobacillus sp. THM7-7* | 2.5 | No evidence on IBD or human gut. | |
| N9296 | *Faecalibacterium* Genus | 2.5 | - | |
| G900184925 | *Ruminococcaceae bacterium HV4-5-B5C* | 2.6 | No link to IBD or human gut. | |
| G001276625 | Achromatium sp. WMS1 | 2.6 | No IBD or human gut links. | |
| G000189595 | *[Clostridium] symbiosum WAL-14163* | 2.7 | Several studies have found enrichement of *Clostridium symbiosum* in IBD samples [48, 58, 134, 135]. | sourmash, sylph |
| G012999505 | *Salifodinibacter halophilus* | 3.0 | No evidence on IBD or human gut. | |
| G003478165 | *Blautia sp. OF01-4LB* | 3.0 | Studies have shown some *Blautia* species (not identified) are linked with IBS and UC [61, 136]. | sourmash, sylph |
| N11762 | *Roseburia inulinivorans* | 3.2 | See G000174195. | |
| G900552255 | *uncultured Eubacteriales bacterium* | 3.4 | - | sylph |
| G000411355 | *Coprococcus sp. HPP0048* | 3.4 | *Coprococcus sp. HPP0048* abundaces are reported to increase from baseline in high-PA PI-IBS subjects [63]. | sourmash, sylph |
| G000156675 | *Blautia hansenii DSM 20583* | 3.7 | Studies have shown some *Blautia* species (not identified) are linked with IBS and UC [61, 136]. | sourmash, sylph |
| G000526735 | *Mediterraneibacter gnavus AGR2154* | 4.6 | Several studies have shown increased *Mediterraneibacter gnavus* abundaces in IBD patients [48, 52, 55, 63, 105]. | sourmash, sylph |

Table S5: EMP dataset. DA features with available taxonomic labels. References of prior evidence are added for some of the features.

| Taxonomy level | Enriched in saline | Enriched in non-saline |
|---|---|---|
| Species | *bacterium TMED88, Planktomarina temperata [137, 138], Proteobacteria bacterium JGI 0000113-P07* | *Acidobacteria bacterium 13_1_40CM_2_64_6, Actinobacteria bacterium 21-64-8, Candidatus Eremiobacteraeota bacterium, Nocardioides convexus, Lautropia sp. SCN 69-89* |
| Genus | *Acinetobacter, Bartonella, Bradyrhizobium [139] , Candidatus Actinomarina, Candidatus Pelagibacter [140], Deinococcus, Liberibacter, Moraxella, Prochlorococcus, Shewanella [141], Staphylococcus, Synechococcus [142]* | *Actinokineospora, Actinoplanes, Amycolatopsis, Anaeromyxobacter [141], Aromatoleum, Aureimonas, Brachybacterium, Bradyrhizobium [71], Brevundimonas, Candidatus Accumulibacter [143], Caulobacter, Conexibacter, Corynebacterium, Cupriavidus, Deinococcus, Gemmata, Haliangium, Hydrogenophaga, Kaistia, Knoellia, Massilia, Methyloversatilis, Microbacterium, Microbispora, Mycolicibacterium, Nocardiopsis, Patulibacter, Phenylobacterium, Reyranella, Rhizorhabdus, Roseomonas, Rubrobacter, Saccharopolyspora, Sandarakinorhabdus, Solirubrobacter, Sphingobium, Sphingomonas, Stenotrophomonas, Streptacidiphilus, Thauera, Trebonia, Verminephrobacter, Wenzhouxiangella* |
| Family | Acetobacteraceae, Anaerolineaceae, Borreliaceae, Comamonadaceae [73], Legionellaceae, Nocardioidaceae, Opitutaceae, Oscillospiraceae, Pseudonocardiaceae, Puniceicoccaceae, Roseobacteraceae [144], Schleiferiaceae, Sphingobacteriaceae, Streptomycetaceae, Woeseiaceae | Acidobacteriaceae [77, 145], Alcaligenaceae [141], Anaerolineaceae, Burkholderiaceae [141], Comamonadaceae [141], Desulfovibrionaceae, Gemmatimonadaceae, Geodermatophilaceae, Hyphomicrobiaceae, Intrasporangiaceae, Isosphaeraceae, Lentisphaeraceae, Methylobacteriaceae, Microbacteriaceae, Micrococcaceae, Opitutaceae, Phyllobacteriaceae, Pirellulaceae, Polyangiaceae, Propionibacteriaceae, Pseudonocardiaceae, Rhodospirillaceae, Roseobacteraceae, Sterolibacteriaceae, Streptomycetaceae, Thermomonosporaceae, Xanthobacteraceae [72] |
| Order | Bacillales [146], Desulfobacterales [147], Neisseriales, Rhodobacterales [148] | Acidithiobacillales, Bacillales, Bryobacterales, Burkholderiales, Hyphomicrobiales, Kineosporiales, Micrococcales, Myxococcales, Sphingomonadales |
| Class | Acidobacteriia, Alphaproteobacteria [138], Bacteroidia, Chitinophagia, Deltaproteobacteria, Negativicutes, Oligoflexia, Tissierellia | Bacilli, Chloroflexia, Hydrogenophilalia, Nitriliruptoria |
| Phylum | Bacteroidetes [77], Gemmatimonadetes [77, 145] | Acidobacteria [77], Armatimonadetes [79], Chloroflexi [147], Gemmatimonadetes [147] |
| Kingdom | Bacillati, CPR, Pseudomonadati [149, 150] | PVC |

Table S6: IBD dataset. Average running time for each method for one sample. ($n \approx 16,000, |\mathcal{X}| \approx 10^7$).

| | Index Construction | | Per Sample Analysis | | |
|---|---|---|---|---|---|
| Method | Wall time | # Cores | Wall time × # Cores | Wall time | # Cores |
| krepp | 2.7 | 24 | 4.8 | 0.3 | 16 |
| DecoDiPhy ‡ | - | - | 0.83 | 0.05 | 16 |
| sylph | 0.01 | 48 | 0.015 | 0.01 | 1 |
| sourmash | 1.2 | 1 | 0.1 | 0.1 | 1 |
| Woltka | 6.1 | 32 | 57.6 | 1.2 | 48 |
| Kraken2 | 5.4 | 1 | 0.2 | 0.2 | 1 |
| Bracken † | 0.2 | 10 | 0.02 | 0.002 | 10 |

‡ The input to DecoDiPhy comes from krepp and thus, its running time needs to be added.
† Running time of Kraken (its input) should be added.

### C Supplementary text

#### C.1 Supplementary Algorithms

#### C.2 Main pseudo-code

**Algorithm S1** Heuristic search. Results $(\mathbf{p}_*, \mathbf{w}_*, \bar{y}_*, \mathbf{e}_*)$ are globally defined. Defaults: $\tau = {}^{10^3}/_{|\mathcal{X}|}; \rho = 3$.

SmallProb($\mathbf{e}$) := $\min, \arg\min_{\mathbf{p},\mathbf{w},\bar{y}}$ Eq. (5) fixing $\mathbf{e}$
**procedure** RePlaceOne($i, k$)
  $\mathcal{N} \leftarrow E_R$
  **if** $i < k$ **then**
    $\mathcal{N} \leftarrow$ edges within radius $\rho$ edge to $\mathbf{e}^*_i$
  **for all** $e \in \mathcal{N} \setminus \mathbf{e}$ **do**
    $\mathbf{e} \leftarrow \mathbf{e}^* \oplus_i e$
    $\ell, (\mathbf{p}, \mathbf{w}, \bar{y}) \leftarrow$ SmallProb($\mathbf{e}$)
    **if** $\ell < \ell^*$ **then**
      $\ell^*, \mathbf{e}^*, \mathbf{p}^*, \mathbf{w}^*, \bar{y}^* \leftarrow \ell, \mathbf{e}, \mathbf{p}, \mathbf{w}, \bar{y}$
      improved $\leftarrow$ True
  **return** improved
**procedure** Search($R, \mathbf{d}$)
  Compute $D$ and $C.L$ from $R$
  **repeat** (starting from $k = 1$)
    $k \leftarrow k + 1$
    $\ell^* \leftarrow \infty$
    **repeat**
      more $\leftarrow$ False
      **for all** $i \in \{k, k-1, \ldots 1\}$ **do**
        more $\leftarrow$ more $\vee$ RePlaceOne($i, k$)
    **until** $\neg$ more
  **until** $(\min(\mathbf{p}^*) < \tau) \vee (\ell^* \approx 0)$
  HandleX0X1($A, \mathbf{p}^*, \mathbf{w}^*$) ▷ See Alg. S2

#### C.3 Handling corner cases for $\mathbf{x}_q = 0$ or $\mathbf{x}_q = 1$

Although, theoretically, we are limiting $x_i \in (0, 1)$, the optimization algorithm might still assign $x_i \approx 0$ or $x_i \approx 1$. These cases might correspond to an invalid solution to the true placement (see Proof of claim 2 for an example scenario). While we have not exhaustively analyzed all potential scenarios where this might occur, empirical observations have been limited to the example provided in Proof of claim 2. To address this issue, when $x_i \approx 1$ $(1 - x_i < 1e - 3)$, we explore all edges adjacent to the placement edge $e_{a_i}$ until a solution with $x_i < 1$ is obtained. Similarly, when $x_i \approx 0$ $(x_i < 1e - 4)$, we examine all child nodes of $a_i$ as possible anchor nodes for $q_i$ and select a solution with $x_i > 0$, if such a solution exists.

**Algorithm S2** Handling corner cases when $\mathbf{x}_q = 0$ or $\mathbf{x}_q = 1$

**procedure** HandleX0X1( $A^{(\hat{k})}, \mathbf{p}^{(\hat{k})}, \mathbf{w}^{(\hat{k})}, \bar{y}^{(\hat{k})}$ )
    **if** $\max(\mathbf{w}^{(\hat{k})}/\mathbf{p}^{(\hat{k})}) \approx 1$ **then**
        $i \leftarrow \arg\max(\mathbf{w}^{(\hat{k})}/\mathbf{p}^{(\hat{k})})$
        $a_i \leftarrow \arg\max A^{(\hat{k})}(., i)$
        **for all** $e = (u, v) \in$ adjacent edges of $a_i$ **do**
            $A^{(\hat{k})}(a_i, i) \leftarrow 0$
            $A^{(\hat{k})}(v, i) \leftarrow 1$
            $\ell_*, (\mathbf{p}_*^{(\hat{k})}, \mathbf{w}_*^{(\hat{k})}, \bar{y}_*^{(\hat{k})}) \leftarrow \min$ (5), $\arg\min$ (5).
            **if** $(\mathbf{w}_*^{(\hat{k})}/\mathbf{p}_*^{(\hat{k})})(i) \not\approx 0$ **AND** $(\mathbf{w}_*^{(\hat{k})}/\mathbf{p}_*^{(\hat{k})})(i) \not\approx 1$ **then**
                $(\mathbf{p}^{(\hat{k})}, \mathbf{w}^{(\hat{k})}, \bar{y}^{(\hat{k})}) \leftarrow (\mathbf{p}_*^{(\hat{k})}, \mathbf{w}_*^{(\hat{k})}, \bar{y}_*^{(\hat{k})})$
    **if** $\min(\mathbf{w}^{(\hat{k})}/\mathbf{p}^{(\hat{k})}) \approx 0$ **then**
        $i \leftarrow \arg\min(\mathbf{w}^{(\hat{k})}/\mathbf{p}^{(\hat{k})})$
        $a_i \leftarrow \arg\max A^{(\hat{k})}(., i)$
        **for all** $v \in$ child nodes of $a_i$ **do**
            $A^{(\hat{k})}(a_i, i) \leftarrow 0$
            $A^{(\hat{k})}(v, i) \leftarrow 1$
            $\ell_*, (\mathbf{p}_*^{(\hat{k})}, \mathbf{w}_*^{(\hat{k})}, \bar{y}_*^{(\hat{k})}) \leftarrow \min$ (5), $\arg\min$ (5).
            **if** $(\mathbf{w}_*^{(\hat{k})}/\mathbf{p}_*^{(\hat{k})})(i) \not\approx 0$ **AND** $(\mathbf{w}_*^{(\hat{k})}/\mathbf{p}_*^{(\hat{k})})(i) \not\approx 1$ **then**
                $(\mathbf{p}^{(\hat{k})}, \mathbf{w}^{(\hat{k})}, \bar{y}^{(\hat{k})}) \leftarrow (\mathbf{p}_*^{(\hat{k})}, \mathbf{w}_*^{(\hat{k})}, \bar{y}_*^{(\hat{k})})$

#### C.4 Divide-and-conquer

Because the running time of DecoDiPhy is worse than linear in the number of leaves $n$ and quadratic in the number of consolidated placements $k$, applying it directly to large trees remains computationally expensive. To address this, we adopt a divide-and-conquer strategy for any reference tree with more than 1000 leaves. We partition the full tree $R$ into subtrees $R^1, \ldots, R^c$ of size approximately $n \approx 1000$ using TreeCluster [94] using its sum-branch mode (with all branch lengths set to 1). For the WoL-2 reference tree used in Experiment E3 and in our biological analyses, this led to $c = 16$ subtrees. Let $\mathcal{X}$ denote the set of (single or multi-) placements for all reads on $R$. Our current divide-and-conquer methods require a way to assign reads to subsets; when the input is read placements, this is trivial. Otherwise, other methods such as classification are available [151].

For each subtree $R^{(i)}$, we extract the subset of read placements that fall within that subtree,

$$\mathcal{X}^{(i)} = \mathcal{X} \cap E_{R^{(i)}}.$$

We then compute the corresponding estimated distance vector $\hat{\mathbf{d}}^{(i)}$ using the same procedure described in *Methods* using $\mathcal{X}^{(i)}$. Running DecoDiPhy independently on each pair $(R^{(i)}, \hat{\mathbf{d}}^{(i)})$ yields a set of consolidated placements $(\mathbf{e}^{(i)}, \mathbf{p}^{(i)}, \mathbf{x}^{(i)}, \bar{y}^{(i)})$. Because each subtree accounts for a fraction $\frac{|\mathcal{X}^{(i)}|}{|\mathcal{X}|}$ of all read placements, after we obtain the solutions, we normalize $\mathbf{p}^{(i)}$ so that its entries sum to that fraction. Finally, we combine the results across all subtrees to obtain the full set of placements:

$$\mathbf{e} = \bigcup_i \mathbf{e}^{(i)}, \qquad \mathbf{x} = \bigcup_i \mathbf{x}^{(i)}, \qquad \bar{y} = \sum_i \frac{|\mathcal{X}^{(i)}|}{|\mathcal{X}|}\, \bar{y}^{(i)}, \qquad \mathbf{p} = \bigcup_i \left( \frac{|\mathcal{X}^{(i)}|}{|\mathcal{X}|}\, \mathbf{p}^{(i)} \right).$$

#### C.5 estimating individual y values

We set $\mathbf{y}_q$ proportionally to the mean height of $C(a_q)$. Let's define the mean height of $C(a_q)$ as $H_{a_q} = \frac{1}{|C(a_q)|} \sum_{l \in C(a_q)} (\mathbf{x}_q + d(l, a_q))$. We set $\mathbf{y}_q = C \cdot H_{a_q}$, such that $\bar{y} = \sum_q \mathbf{p}_q \cdot \mathbf{y}_q = C \cdot \sum_q \mathbf{p}_q \cdot H_{a_q}$.

#### C.6 Supplementary Proofs and Claims

While all claims are presented before claim 4, we will use Eq. (4) from claim 4 in the proofs.

##### C.6.1 Proof of Claim 1

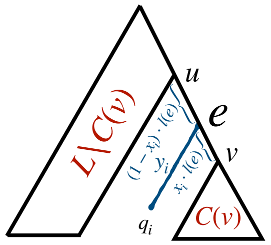

Figure S19: An example of a query $q_i$ on the edge $e = (u, v)$. The distance of $q_i$ to each of the nodes $u$ and $v$ can be shown with $(1 - x_i) \cdot l(e) + y_i$ and $x_i \cdot l(e) + y_i$, respectively.

For a placement of a query $q_i$ on tree $R$, we define $x_i$ as the relative position of the connecting point of $q_i$ on its placement edge $e = (u, v)$ to node $v$, and terminal distance $y_i$ as the distance of $q_i$ to its connecting point on $e$ (see Fig. S19). Let queries $\mathcal{Q} = q_1, q_2, \ldots, q'_k$ share the same placement edge $e = (u, v)$. Let $\mathbf{p} = \{p_1, p_2, \ldots, p'_k\}$ be the corresponding query weights, $\mathbf{x} = \{x_1, x_2, \ldots, x'_k\}$ be their relative position on the branch $e$, and $y = \{y_1, y_2, \ldots, y'_k\}$ be the terminal distances. Let $q'$ be a single query on the same edge with weight $p_{q'} = \sum_{i=1}^{k'} p_i$, relative position $x_{q'} = \frac{\sum_{i=1}^{k'} p_i \cdot x_i}{\sum_{i=1}^{k'} p_i}$, and terminal distance $y_{q'} = \frac{\sum_{i=1}^{k'} p_i \cdot y_i}{\sum_{i=1}^{k'} p_i}$. By definition, $0 \leq x_i \leq 1$ for all $i \in \{1, 2, \ldots, k'\}$, ensuring $0 \leq x_{q'} \leq 1$. Thus, $q'$ can be placed on the same edge $e$. The edge $e = (u, v)$ creates a bipartition of the set of leaves $\mathcal{R}$: The set of leaves under $v$ (i.e., $C(v)$) and the rest of the leaves ($\mathcal{R} \setminus C(v)$). We show that the average distance of $\mathcal{Q}$ to any leaf $l \in \mathcal{R} \setminus \mathcal{Q}$ equals the weighted distance of $q'$ to $l$, whether $l \in C(v)$ or $l \in \mathcal{R} \setminus C(v)$.

- **Case 1:** $l \in C(v)$

  For any $q_i \in \mathcal{Q}$, the distance to $l$ equals:

  $$d(q_i, l) = d(v, l) + x_i \cdot l(e) + y_i,$$

  where $d(v, l)$is the distance from $v$ to $l$ (see Fig. S19). The average distance of $\mathcal{Q}$ to $l$ is:

  $$d_l = \sum_{i=1}^{k'} p_i \cdot (d(v, l) + x_i \cdot l(e) + y_i).$$

  Expanding this expression:

  $$d_l = \sum_{i=1}^{k'} p_i \cdot d(v, l) + \sum_{i=1}^{k'} p_i \cdot x_i \cdot l(e) + \sum_{i=1}^{k'} p_i \cdot y_i.$$

  Using the definitions of $p_{q'}$, $x_{q'}$, and $y_{q'}$, this simplifies to:

  $$d_l = p_{q'} \cdot (d(v, l) + x_{q'} \cdot l(e) + y_{q'}).$$

  Thus, $d_l = d'_l$, the distance of $q'$ to $l$.

- **Case 2:** $l \in \mathcal{R} \setminus C(v)$

  For any $q_i \in \mathcal{Q}$, the distance to $l'$ is:

  $$d(q_i, l') = d(u, l) + (1 - x_i) \cdot l(e) + y_i,$$

  where $d(u, l')$ is the distance from $u$ to $l'$. The average distance of $\mathcal{Q}$ to $l'$ is:

  $$d'_l = \sum_{i=1}^{k'} p_i \cdot (d(u, l') + (1 - x_i) \cdot l(e) + y_i).$$

  Expanding this expression:

  $$d'_l = \sum_{i=1}^{k'} p_i \cdot d(u, l') + \sum_{i=1}^{k'} p_i \cdot (1 - x_i) \cdot l(e) + \sum_{i=1}^{k'} p_i \cdot y_i.$$

Rewriting $\sum_{i=1}^{k'} p_i \cdot (1 - x_i) \cdot l(e)$ as $p_{q'} \cdot (1 - x_{q'})$, this becomes:

$$d_{l'} = p_{q'} \cdot (d(u, l') + (1 - x_{q'}) \cdot l(e) + y_{q'}).$$

Thus, $d_{l'} = d'_{l'}$, the distance of $q'$ to $l'$.

Therefore, any set of queries $\mathcal{Q}$ on the same branch can be replaced by a single query $q'$ with:

$$p_{q'} = \sum_{i=1}^{k'} p_i, \qquad x_{q'} = \frac{\sum_{i=1}^{k'} p_i \cdot x_i}{\sum_{i=1}^{k'} p_i}, \qquad y_{q'} = \frac{\sum_{i=1}^{k'} p_i \cdot y_i}{\sum_{i=1}^{k'} p_i}.$$

□

##### C.6.2 Proof of Claim 2 (Unidentifiability of $x_i \in 0, 1$)

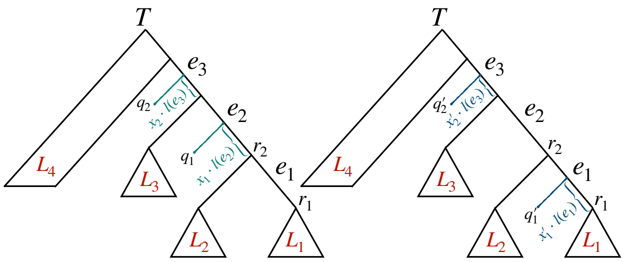

Figure S20: An example of two sets of placements leading to the same solution to Problem 1 if $x'_1 == 1$.

Consider the counterexample shown in Fig. S20. Let the placement of $q_1$ and $q_2$ on $R$ (left panel) correspond to the true placement of the query taxa, with corresponding relative positions $x_1$ and $x_2$, and proportions $p_1$ and $p_2$. Let $q'_1$ and $q'_2$ correspond to an alternative placement of these queries on tree $R$. If $d_l$ is the average distance of queries $q_1$ and $q_2$ to the leaf $l$, and $d'_l$ is the average distance of queries $q'_1$ and $q'_2$ to $l$. Therefore, $d_l = d'_l$ for any leaf $l$. For simplification, we assume the average terminal branch lengths for the two placements are equal ($\bar{y} = \bar{y'}$). Also, by definition, $p_1 + p_2 = p'_1 + p'_2 = 1$. Using the equations in claim 4, we can write the equations for leaf $l1 \in L_1$ and $l_2 \in L_2$:

$$\begin{aligned} l_1 : & p_1 \cdot (d(l_1, r_1) + l(e_1) + x_1 \cdot l(e_2)) + p_2 \cdot (d(l_1, r_1) + l(e_1) + l(e_2) + x_2 \cdot l(e_3)) = \\ & p'_1 \cdot (d(l_1, r_1) + x'_1 \cdot l(e_1)) + p'_2 \cdot (d(l_1, r_1) + l(e_1) + l(e_2) + x'_2 \cdot l(e_3)). \end{aligned}$$

$$\begin{aligned} l_2 : & p_1 \cdot (d(l_2, r_2) + x_1 \cdot l(e_2)) + p_2 \cdot (d(l_2, r_2) + l(e_2) + x_2 \cdot l(e_3)) = \\ & p'_1 \cdot (d(l_2, r_2) + l(e_1) - x'_1 \cdot l(e_1)) + p'_2 \cdot (d(l_2, r_2) + l(e_2) + x'_2 \cdot l(e_3)). \end{aligned}$$

By simplifying these two equations and using $p_1 + p_2 = p'_1 + p'_2$ , we get:

$$p_1 \cdot l(e_1) + p_2 \cdot l(e_1) = 2p'_1 \cdot x'_1 \cdot l(e_1) - p'_1 \cdot l(e_1) + p'_2 \cdot l(e_1)$$

Therefore:

$$p'_1 \cdot x'_1 \cdot l(e_1) = p'_1 \cdot l(e_1)$$

.

As a result, if $x_i = 1$ is allowed for any query $q_i$, we will lose identifiability. A similar example can be constructed for $x_i = 0$ (by rerooting the example tree $R$ on $l_1$). □

###### C.6.3 Proof of Claim 2 (Unidentifiability of individual y values)

Let $\mathcal{Q}$ be a set of $k$ sample queries with anchor nodes $a = \{a_1, a_2, \ldots, a_k\}$, weights $\mathbf{p} = \{p_1, p_2, \ldots, p_k\}$, position vector $x = \{x_1, x_2, \ldots, x_k\}$, and terminal branch lengths $y = \{y_1, y_2, \ldots, y_k\}$. Using Eq. (3) and claim 4, we can rewrite Eq. (3) as:

$$d_j = \sum_{i=1}^{k} p_i \cdot (d(a_i, r_j) + \mathbf{1}_{r_j \in C(a_i)} \cdot (x_i \cdot l(e_{a_i})) + y_i)$$

where $\mathbf{1}_{r_j \in C(a_i)}$ is the indicator function, equal to one if $r_j \in C(a_i)$, and 0 otherwise. This expression can be separated into two components:

$$d_j = \underbrace{\sum_{i=1}^{k} p_i \cdot (d(a_i, r_j) + \mathbf{1}_{r_j \in C(a_i)}(x_i \cdot l(e_{a_i})))}_{\text{first term}} + \bar{y}$$

where $\bar{y} = \sum_{i=1}^{k} p_i \cdot y_i$ is the weighted average of the terminal branch lengths. The first term in the expression depends only on the anchor nodes, the query positions, and the distances $d(a_i, r_j)$ and is independent from the specific values of $y$. The second term, $\bar{y}$, contributes a constant offset to $d_j$ for all $r_j$. This offset depends only on the weighted average $\bar{y}$, not on the individual values of $y_1, y_2, \ldots, y_k$. Since $\bar{y}$ is the only contribution from $y$ to the average distance $d_j$, any set of terminal branch lengths $y' = \{y'_1, y'_2, \ldots, y'_k\}$ that satisfies $\bar{y'} = \sum_{i=1}^{k} p_i \cdot y'_i = \bar{y}$ will result in the same average distances $d_j$ for all $j$. Therefore, it suffices to recover $\bar{y}$ rather than the individual values of $y$. □

###### C.6.4 Proof of Claim 3

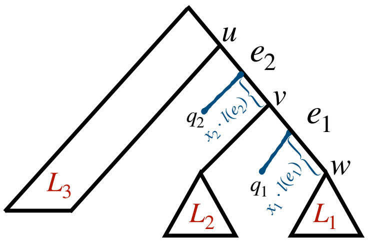

Figure S21: An example of two adjacent true placement edges $e_1$ and $e_2$. Queries $q_1$ and $q_2$ are placed on $e_1$ and $e_2$, respectively.

Let $\mathcal{E}$ be the set of true placement edges for the query sample $\mathcal{Q}$. Let the placement edges $e_1, e_2 \in \mathcal{E}$ be two adjacent placement edges and $v$ be their connecting node. Claim 3 states that there are infinite solutions to Problem 1, by keeping the placements edges $\mathcal{E}$ fixed, and only changing abundances $\mathbf{p}$ and placement positions $\mathbf{x}$.

**Proof.** Without losing generality, consider a rooting of the tree where $e_1$ is an outgoing edge of $v$ and $e_2$ is an incoming edge to $v$ (see Fig. S21). Let $q_1$ and $q_2$ be the queries on the edges $e_1$ and $e_2$, respectively. For any reference leaf $l$, assume $d(\mathcal{Q} \setminus \{q_1, q_2\}, l)$ is the true average distance from all the other query taxa to the leaf $l$. Also, let $\bar{y}_{1,2}$ be the true weighted average of the terminal branch lengths $y_1$ and $y_2$ corresponding to $q_1$ and $q_2$. Let $d'_l$ be the average distance vector after subtracting $d(\mathcal{Q} \setminus \{q_1, q_2\}, l)$ from $d_l$. According to Eq. (3) and claim 4, there exists a set of variables $x_1$, $x_2$, $p_1$, and $p_2$ that satisfies:

$$p_1 \cdot (d(w,l) + \mathbf{1}_{l \in C(w)}(x_1 \cdot l(e_1))) + p_2 \cdot (d(v,l) + \mathbf{1}_{l \in C(v)}(x_2 \cdot l(e_2))) = d'_l.$$

for all $l \in \mathcal{R}$. Using the same trick as Eq. (4), we can rewrite the above equation as:

$$p_1 \cdot d(w,l) + \mathbf{1}_{l \in C(w)}(w_1 \cdot l(e_1)) + p_2 \cdot d(v,l) + \mathbf{1}_{l \in C(v)}(w_2 \cdot l(e_2)) = d'_l.$$

where $w_1 = x_1 \cdot p_1$ and $w_2 = x_2 \cdot p_2$. Since this equation is linear with respect to the variables $p_1$, $p_2$, $w_1$, and $w_2$, we need at least four linearly independent equations to find unique solutions for each variable. Note that the introduction of $w_1$ and $w_2$ introduces new constraints that further limit the solution space; however, for the sake of this proof, we ignore these constraints. Since the true proportions of all the other queries are known, $p_1 + p_2 = 1 - \sum_{i \neq 1,2}^{k} p_i = c$ is a constant. Therefore, we need at least three more linearly independent equations that are also independent from the equation $p_1 + p_2 = c$. The set of leaves $\mathcal{R}$ on the tree is partitioned into three sets $L_1$, $L_2$, and $L_3$. Each leaf yields to an equation. Taking two leaves from the same set will not result in two linearly independent equations. We will use two leaves $l_1$ and $l'_1$ from $L_1$ as an example, but the rest follows:

$$l_1 : p_1 \cdot d(l_1, w) + l(e_1) \cdot w_1 + p_2 \cdot (d(l_1, w) + p_2 \cdot l(e_1) + l(e_2) \cdot w_2 = d'_{l_1} - \bar{y}_{1,2}$$

$$l'_1 : p_1 \cdot d(l'_1, w) + l(e_1) \cdot w_1 + p_2 \cdot (d(l'_1, w) + p_2 \cdot l(e_1) + l(e_2) \cdot w_2 = d'_{l'_1} - \bar{y}_{1,2}$$

By subtracting the two equations, we will get the following equation:

$$(p_1 + p_2) \cdot (d(l_1, w) - d(l'_1, w)) = d'_{l_1} - d'_{l'_1}$$

Since $p_1 + p_2 = c$ is a constant, the two equations are not linearly independent, and similarly, any two leaves from the same set will result in two linearly dependent equations.

Therefore, only leaves taken from different sets might result in linearly independent equations. Since there are three disjoint sets of leaves $L_1$, $L_2$, and $L_3$, we will take one leaf from each set: $l_1 \in L_1$, $l_2 \in L_2$, and $l_3 \in L_3$. We will write the equations corresponding to $l_1$, $l_2$, and $l_3$, respectively:

$$l_1 : p_1 \cdot d(l_1, w) + l(e_1) \cdot w_1 + p_2 \cdot (d(l_1, w) + p_2 \cdot l(e_1) + l(e_2) \cdot w_2 = d'_{l_1} - \bar{y}_{1,2}$$

$$l_2 : p_1 \cdot d(l_2, v) + l(e_1) \cdot p_1 - l(e_1) \cdot w_1 + p_2 \cdot (d(l_2, v) + l(e_2) \cdot w_2 = d'_{l_2} - \bar{y}_{1,2}$$

$$l_3 : p_1 \cdot d(l_3, u) + (l(e_1) + l(e_2)) \cdot p_1 - l(e_1) \cdot w_1 + p_2 \cdot (d(l_3, u) + l(e_2) \cdot p_2 - l(e_2) \cdot w_2 = d'_{l_3} - \bar{y}_{1,2}$$

Adding the first equation to the second equation results in the following equation:

$$(p_1 + P_2) \cdot (d(l_1, w) + d(l_2, v)) + (p_1 + p_2) \cdot l(e_2) = d'_{l_1} + d'_{l_2} - 2\bar{y}_{1,2}$$

Since $p_1 + p_2 = c$ is a constant, equations 2 and 3 are not linearly independent, and therefore, we have fewer equations than variables, and if a set of variables $p_1$, $p_2$, $w_1$, and $w_2$ is a solution to this system, this system has infinite solutions. □

##### C.6.5 Proof of claim 4

Let $D(F) \in \mathbb{R}^{n \times k}$ be the true distance matrix where $D(F)_{(r,q)} = d_F(q, r)$. Problem 1 is expressed as:

$$\mathbf{d} = D(F) \cdot \mathbf{p} \ .$$

The pairwise distance between a query taxon $q$ and a reference taxon $r$ can be rewritten as:

$$d(q, r) = \begin{cases} d(r, a_q) + \mathbf{x}_q \cdot l(e(a_q)) + \mathbf{y}_q, & \text{if } r \in C(a_q) \\ d(r, a_q) - \mathbf{x}_q \cdot l(e(a_q)) + \mathbf{y}_q, & \text{if } r \notin C(a_q) \end{cases}$$

Recall:

- $D \in \mathbb{R}^{n \times m}$: A distance matrix equivalent to tree $R$ where $D_{r,v} = d_R(r, v)$ for $r \in \mathcal{R}, v \in V_R$.
- $C \in \{-1, 1\}^{n \times m}$ is a matrix, where $C_{r,v} = 1$ iff $r \in C(v)$ ($r$ descends from $v$) and $C_{r,e} = -1$ otherwise.

Thus, $d_F(q, r)$ can be written as:

$$d_F(q, r) = D_{(r,a_q)} + C_{(r,a_q)} \cdot \mathbf{x}_q \cdot l(e(a_q)) + \mathbf{y}_q$$

Consequently:

$$D(F) = D \cdot A + C \cdot L \cdot A \cdot \mathbf{x} + \mathbf{y}$$

And therefore:

$$\mathbf{d} = D(F) \cdot \mathbf{p} = (D \cdot A + C \cdot L \cdot A \cdot \mathbf{x} + y) \cdot \mathbf{p}$$

Note that variables $\mathbf{x}$ and $\mathbf{p}$ are multiplied. To get to a linear formulation, we introduce $\mathbf{w} = \mathbf{x} \circ \mathbf{p}$ and due to lack of identifiability, we introduce $\bar{y} = \sum_{i=1}^{k} \mathbf{p}_i \cdot \mathbf{y}_i$ to get:

$$\mathbf{d} = (D \cdot A + C \cdot L \cdot A \cdot \mathbf{x} + y) \cdot \mathbf{p} = D \cdot A \cdot \mathbf{p} + C \cdot L \cdot A \cdot \mathbf{w} + \bar{y} \cdot \mathbf{1}_n,$$

Since $\mathbf{x}_i \in (0, 1)$ and $\mathbf{p}_i \in (0, 1)$, any $\mathbf{w}_i \leq \mathbf{p}_i$ would result in a valid $\mathbf{x}_i = \frac{\mathbf{w}_i}{\mathbf{p}_i}$. To ensure $\mathbf{w}_i \leq \mathbf{p}_i$, add the linear constraint $\mathbf{w}_i \leq \mathbf{p}_i$, $\forall i \in \{1, 2, \ldots, k\}$.

□

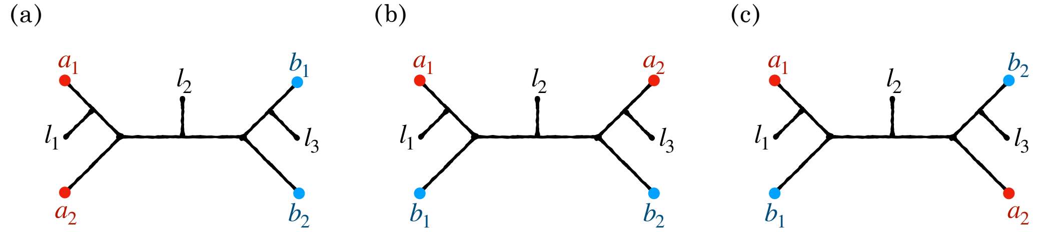

Figure S22: All possible placements of two sets of queries of size 2. The minimum number of leaves needed for the placement edges to be different for each query is shown on each tree.

###### C.6.6 Identifiability of Problem 1 for case $k = 2$

**Claim 5.** *For the case of $k = 2$ and $p_i = \frac{1}{2}$ for all queries $q_i$, the true placement edges can be uniquely recovered.*

*Proof.* Let $A = \{a_1, a_2\}$ and $B = \{b_1, b_2\}$ be two different placements for queries $q_1$ and $q_2$. Fig. S22 shows all possible configurations of placements $A$ and $B$. We assume that no two queries share the same placement edge; therefore, the minimum number of leaves needed for this assumption to be true (3 leaves) is added to each tree. Let $d(i, j)$ denote the distance between two nodes $i$ and $j$ on the tree, and let $d(A, l_i)$ and $d(B, l_i)$ denote the average distance of sets $A$ and $B$ to leaf $l_i$. More specifically, $d(A, l_i) = \frac{d(a_1,l_i)+d(a_2,l_i)}{2}$ and $d(B, l_i) = \frac{d(b_1,l_i)+d(b_2,l_i)}{2}$.

For each case, we first assume that the two sets of placements are both valid. In other words, $d(A, l_i) = d(B, l_i)$ for all $i$. We then use the four-point condition [81] between the leaves of the tree and the queries to prove otherwise.

- **Case 1:** Take leaves $l_1$ and $l_2$. We'll write the four-point condition once between $a_1$, $b_1$, $l_1$, and $l_2$, and once between $a_2$, $b_2$, $l_1$, and $l_2$:

$$d(a_1, l_1) + d(b_1, l_2) < d(a_1, l_2) + d(b_1, l_1)$$

$$d(a_2, l_1) + d(b_2, l_2) < d(a_2, l_2) + d(b_2, l_1)$$

  By summing these two inequalities, we'll get: $2d(A, l_1) + 2d(B, l_2) < 2d(A, l_2) + 2d(B, l_1)$, which is a contradiction to $d(A, l_1) = d(B, l_1)$ and $d(A, l_2) = d(B, l_2)$.

- **Case 2:** Take leaves $l_1$ and $l_2$. We'll write the four-point condition once between $a_1$, $b_1$, $l_1$, and $l_3$, and once between $a_2$, $b_2$, $l_1$, and $l_3$:

$$d(a_1, l_1) + d(b_1, l_2) < d(a_1, l_2) + d(b_1, l_1)$$

$$d(a_2, l_1) + d(b_2, l_2) = d(a_2, l_2) + d(b_2, l_1)$$

  By summing these two statements, we'll get: $2d(A, l_1) + 2d(B, l_2) < 2d(A, l_2) + 2d(B, l_1)$, which is a contradiction to $d(A, l_1) = d(B, l_1)$ and $d(A, l_2) = d(B, l_2)$.

- **Case 3:** Take leaves $l_1$ and $l_3$. We'll write the four-point condition once between $a_1$, $b_1$, $l_1$, and $l_3$, and once between $a_2$, $b_2$, $l_1$, and $l_3$:

$$d(a_1, l_1) + d(b_1, l_3) < d(a_1, l_3) + d(b_1, l_1)$$

$$d(a_2, l_1) + d(b_2, l_3) = d(a_2, l_3) + d(b_2, l_1)$$

  By summing these two inequalities, we'll get: $2d(A, l_1) + 2d(B, l_3) < 2d(A, l_3) + 2d(B, l_1)$, which is a contradiction to $d(A, l_1) = d(B, l_1)$ and $d(A, l_3) = d(B, l_3)$.

Therefore, no two separate sets of queries for the case of $k = 2$ and $p_i = \frac{1}{2}$ for all $i$ exist such that they both represent a possible placement of queries $q_1$ and $q_2$ on the tree. □

#### C.7 Experimental details

##### C.7.1 Runtime analysis

To evaluate the empirical running time of our algorithm, we used a 10,000-taxon simulated dataset studied by [102], which includes 10 replicates. These replicates feature gene trees that differ from the species tree due to both incomplete lineage sorting (ILS) and horizontal gene transfer (HGT), simulated using SimPhy. For each replicate, the first estimated gene tree was used as the backbone tree. We randomly selected $n \in \{50, 100, 200, 500, 1000\}$ taxa from the gene tree and pruned it to include only the selected $n$ taxa. For each replicate and $k$ ($k \in \{1, 2, \ldots, 10\}$ in this experiment), we generated 10 sets of query taxa and measured the running time of our method across different steps of the algorithm.

##### C.7.2 E1 Details

For each dataset, an available species tree was used as $F$; all these trees have branch lengths in units of substitutions per site, as assumed by DecoDiPhy.

**Drawing p.** We draw $k$ values from a uniform distribution, normalized them to add up to one, and dismissed replicates that lead to draws with $\mathbf{p}_q < {}^{0.1}/_{k^2}$ for any $q$. We use $k^2$ because this corresponds to the minimum of $k$ values, each drawn from an exponential distribution, followed by normalization.

**Add noise to true placements.** For each query species $q$ with true placement edge $\mathbf{e}_q$, we generated $\mathbf{p}_q \times 10^5$ simulated placements. Each such placement $x$ was on an edge $\mathbf{e}_x$ that could be potentially different from $\mathbf{e}_q$. First, the number of edges between $\mathbf{e}_x$ and $\mathbf{e}_q$ was drawn from an exponential distribution with scale ${}^1/_\lambda$ and rounded to an integer. Let the drawn distance be $d$. Among all edges at distance $d$ from $\mathbf{e}_q$, one was selected uniformly at random per $x$. If no such edge exists at distance $d$, we choose an edge among those that are furthest away from $\mathbf{e}_q$. Note that when $d = 0$, the read was placed on $\mathbf{e}_q$. For each placement, we set $\mathbf{x}_x = 0.5$ and $\mathbf{y}_x$ to the average height of the sister clade. The mixture distance vector $\hat{\mathbf{d}}$ was then computed as the mean of the three distances from these placements to each reference. We test ${}^1/_\lambda \in 1, 2$. Due to rounding and the limit on how far each node can move, this procedure leads to 0.6 and 1.56 edges of movement on average for our two $\lambda$ values.

**Alternative methods** The following are alternative methods that we designed and compared against. Each uses the small problem from DecoDiPhy, but to different extents.

- `reverse-search`: This approach solves the same small-problem as DecoDiPhy. Recall that for any fixed set of queries, we can solve the small-problem. This approach starts from $\mathbf{e} = E_R$, and solves the small problem, iteratively remove queries with $\mathbf{p}_q < 0.01$ from the fixed set of placement edges, until only $k$ queries remain. If $|\mathcal{Q}| > k$ and $min(\mathbf{p}_q) > 0.01$, it iteratively removes the query with minimum $\mathbf{p}_q$ and solves the small-problem for the rest of the queries until $|\mathcal{Q}| = k$.
- $k$`-nearest`: First, references are sorted based on their distance to the query, and $k$ nearest ones are selected as anchors. The small-problem is then solved on $\mathcal{Q}$ to estimate $\mathbf{p}$, $\mathbf{x}$, and $\bar{y}$.
- `Iterative` $k$`-nearest`: We run $k$ rounds of optimization. In each round, just like `k-nearest`, we select $k$ nearest references to the query and optimize for $\mathbf{p}$, $\mathbf{x}$, and $\bar{y}$. Then, the leaf $l$ with the highest abundance that is not already fixed in prior runs is added to the list of fixed anchors. Before the next

round starts, the distance vector $d$ is updated by subtracting the distance of $l$ to all reference taxa, setting its abundance to $\frac{1}{k}$ (computed $p_i$s are often 0 and not reliable). When $k$ anchors are fixed, a final optimization step is performed to get the final $\mathbf{p}$, $\mathbf{x}$, and $\bar{y}$.

##### C.7.3 E2: Impact of distance calculation.

We started from the WoL-1 10,575-leaf prokaryotic reference tree [152] and selected two substrees with diameter $\approx 0.3$ without a large polytomy (defined as fewer than 20% of branch lengths $< 0.001$). For each replicate of each $k$ of each subtree, we sample $\mathbf{p}$ from either uniform or exponential distributions ($\lambda = 1$). We ensured that $\mathbf{p}_q \geq {}^{0.1}/_{k^2}$ for all query taxa $i \in \{1, \ldots, k\}$

We computed $\hat{\mathbf{d}}$ in three ways: using sequence distances, single placements, and multi-placements. Both sequence distances and placements were computed using krepp (v.0.5.1), respectively, with commands:

- `krepp dist --no-filter --multi -i $KREPP_DB --filter -q $QUERY.fastq`
- `krepp place -i $KREPP_DB -t $BACKBONE.nwk --filter --no-multi -q $QUERY.fastq`
- `krepp place -i $KREPP_DB -t $BACKBONE.nwk --filter --multi -q $QUERY.fastq`

For all krepp computations, we used the same configuration for the indexing step:

- `krepp index -k 29 -w 35 -m 16 -r 15 --frac -h 14 -o $KREPP_DB -i $REFERENCE_PATHS`

For placements, we defined distances using tree paths. For multi-placements, we defined $d_{\hat{F}}(r, x)$ as the unweighted average of distances across all placements for each read before computing $\hat{\mathbf{d}}$. Since some references may receive too few mapped reads for reliable distance estimation, for sequence distances, we excluded any reference from $R$ with fewer than 30% of reads mapped.

##### C.7.4 E3: Realistic metagenomic simulations.

Human microbiome samples analyzed consist of the following seven body sites: stool ($s = 78$), tongue dorsum ($s = 42$), supragingival plaque ($s = 33$), buccal mucosa ($s = 28$), retroauricular crease ($s = 13$), posterior fornix ($s = 10$), and anterior nares ($s = 6$). We used OGU feature values computed using Woltka by Zhu *et al.* [12] to calculate an abundance profile over genomes in WoL-v2 [46] for each sample $i$, denoted by $\mathbf{p}^{(i)}$. For the high novelty case, we pruned all queries that have a non-zero abundance in $\mathbf{p}^{(i)}$ from the original WoL-v2 tree to create a backbone phylogeny for each replicate. Since these queries reflect empirical human microbiome profiles, they are expected to be correlated and aggregated in certain clades in the full phylogeny. This resulted in high novelty: A query whose close neighbors were also pruned lacks sufficient representation in the pruned phylogeny and becomes very novel (average distance of 0.16 to the closest taxon; Figure S3b). In this model condition, on average 90% of the queries did not have a genome from the same species present in the pruned phylogeny (Figure S3c). As an alternative to pruning all queries from the backbone, we created a low-novelty case by ensuring that at least one reference was retained in each small enough clade. This resulted in average distance of 0.04 to the closest taxon (Figure S3b), and 48% novelty in terms of species (Figure S3c). We achieved this by clustering leaves of the WoL-v2 phylogeny using TreeCluster [94] with the criterion that each subset should have a maximum diameter of 0.1 (`-m max` mode). For clusters (including singletons) that entirely appear in the query set of a replicate, we retained one randomly selected query in the backbone tree and only removed the rest. Thus, we guarantee that each query has at least one reference within a 0.1 distance radius (0.1 phylogenetic distance translates crudely to 11% Hamming distance). This procedure resulted in larger backbone trees containing some reference leaves overlapping with queries (Figure S3d) and significantly reduced the overall novelty (Figure S3b).

###### C.7.5 All methods used in biological analysis.

Using krepp (v0.4.4), we performed phylogenetic multi-placement of reads with default filtering parameters of krepp using the command below.

- `krepp place -i $KREPP_DB -t $BACKBONE.nwk --filter --multi -q $QUERY.fastq`

For all krepp computations, we used the following command to index references:

- `krepp index -k 29 -w 35 -m 16 -r $INDEX --frac -h 14 -o $KREPP_DB -i $REFERENCE_PATHS`

For DecoDiPhy, we divided the WoL2 reference tree into 16 subtrees based on Section C.4. We then compute the distance vector from krepp placements, and use them as input to DecoDiPhy:

- `decodiphy -d $KREPP_DISTANCES -t $REFERENCE_SUBTREE -o $OUTPUT`

We constructed the Kraken2 (v2.1.1) database and ran query search using the following commands with default parameters. The database comprised 16k WoL2 reference genomes with taxonomy exactly matching the WoL2 tree:

- `kraken2-build --build --db $KRKN_DB`
- `kraken2 --db $KRKN_DB $QUERY.fastq --output $OUTPUT.kraken --report $OUTPUT.report`

We generated the Bracken (v.2.5.3) database and ran Bracken for abundance estimation using the following commands:

- `bracken-build -d $KRKN_DB -t 10`
- `bracken -d $KRKN_DB -i $QUERY.fastq -o $OUTPUT.bracken -w $OUTPUT.report`

For SYLPH, we created a database of reference fasta files and ran query profiling using:

- `sylph sketch -g $REFERENCE_GENOMES_DIR/*.fna -o $SYLPH_DB -t 48`
- `sylph profile $SYLPH_DB $QUERY.fastq > $OUTPUT.tsv`

For sourmash, we utilized the following commands:

- We sketched reference genomes and built an indexed database from signatures using:
  - `sourmash sketch dna $QUERY.fastq -o $OUTPUT.sig.zip`

  We sketched query genomes and searched for the closest genomes in the reference database with:
  - `sourmash gather $OUTPUT.sig.zip $SMASH_DB.sbt.zip > $OUTPUT.tsv`

We aligned query reads with Bowtie2 (v.2.4.2) and summarized using Woltka:

- `bowtie2-build --large-index $INPUT.fasta $OUTPUT_DB`
- ```
  bowtie2 -p 48 -x $DB_INDEX -t -q -U $QUERY.fastq --xeq
  --very-sensitive --all --np 1 --mp "1,1" --rdg "0,1" --rfg "0,1"
  --score-min "L,0,-0.05" | woltka classify --no-demux -i - -o $OUTPUT.biom
  ```
  ```

##### C.7.6 Biological analyses

We selected 210 free-living samples from the Earth Microbiome Project with a balanced saline ($s = 106$) and non-saline ($s = 104$) composition. Among these, 91 and 67 samples were solid, and the rest were aqueous for non-saline and saline samples, respectively. In the IBD dataset, 164 of 220 samples were from diseased subjects, and 56 were from controls.

In calculating error metrics for evaluations, we ignore $\mathbf{x}$ and $\mathbf{y}$ since other methods do not produce them.

- Computing pairwise UniFrac and wUniFrac distances: For computing pairwise distances, a feature table input is provided where features are branches of the phylogeny, and values are predicted abundances for each feature.

```
qiime diversity beta-phylogenetic \
      --i-table $FEATURE_TABLE.qza \
      --i-phylogeny $TREE.qza \
      --p-metric [un]weighted_unifrac \
      --o-distance-matrix $DISTANCE_MATRIX.qza

qiime diversity beta-group-significance \
      --i-distance-matrix $DISTANCE_MATRIX.qza \
      --m-metadata-file $METADATA \
      --m-metadata-column $GROUP \
      --o-visualization $OUTPUT\
      --p-permutations 1000
```

- Computing pairwise Bray-Curtis distances:

```
qiime diversity beta \
  --i-table $FEATURE_TABLE.qza \
  --p-metric braycurtis \
  --o-distance-matrix $DISTANCE_MATRIX.qza

qiime diversity beta-group-significance \
  --i-distance-matrix $DISTANCE_MATRIX.qza \
  --m-metadata-file $METADATA \
  --m-metadata-column $GROUP \
  --o-visualization $OUTPUT\
  --p-permutations 1000
```

- Computing Faith's phylogenetic diversity:

```
qiime diversity alpha-phylogenetic \
  --i-table $FEATURE_TABLE.qza \
  --i-phylogeny $TREE.qza \
```

```
    --p-metric faith_pd \
    --o-alpha-diversity  $DIVERSITY_OUTPUT.qza

  qiime diversity alpha-group-significance \
    --i-alpha-diversity $DIVERSITY_OUTPUT.qza \
    --m-metadata-file $METADATA \
    --o-visualization $OUTPUT
```

- Differential abundance (DA) analysis with ANCOM-BC: For the DA analysis, the input is the same feature table as pairwise distance inputs, with the difference that values emulate the number of mapped reads to each feature instead of abundances. For each sample, we filter features with fewer than 100 reads mapped to them before performing ANCOM-BC.

```
qiime feature-table filter-features \
  --i-table $FEATURE_TABLE.qza\
  --p-min-frequency 100 \
  --o-filtered-table $FILTERED_FEATURE_TABLE.qza

qiime composition ancombc \
  --i-table $FILTERED_FEATURE_TABLE.qza\
  --m-metadata-file $METADATA \
  --p-formula $GROUP \
  --o-differentials $OUTPUT
```
